# The genetic origin of fetal growth restriction and mitochondrial complex I dysregulation

**DOI:** 10.1101/2025.06.19.660636

**Authors:** Joshua J. Fisher, Carol A. Wang, Veronica B. Botha, Siddharth Acharya, Heather C. Murray, John E. Schjenken, Craig E. Pennell, Roger Smith

## Abstract

**Background:** Mitochondria are organelles required for bioenergetic homeostasis, providing energy in the form of ATP to support cellular function, growth, and proliferation. Mitochondria synthesise ATP using the electron transport chain (ETC) to produce an electrochemical gradient that facilitates the conversion of ADP to ATP by ATP synthase. Mitochondrial function is governed by an intricate bidirectional relationship between the mitochondrial genome encoding 1-2% of proteins and the nuclear genome which encodes 98-99% of mitochondrial proteins. Pregnancy creates a unique environment whereby the mitochondrial genome of the placenta is maternally inherited, while the nuclear genome is comprised of both maternal and paternal contributions. Thus, mitochondrial structure and function are largely dependent on the adaptability of mitochondria to conform to the new mixed nuclear genome. In pregnancy, fetal growth restriction (FGR) is characterised by poor placental development, underlying trophoblast insufficiencies, and is associated with mitochondrial dysfunction within the placenta, although the mechanisms which underpin these changes remain unclear. This study investigated if mitochondrial dysfunction in FGR is programmed by genetic incompatibility arising from single nucleotide polymorphisms (SNPs).

**Methods and findings:** We performed a targeted meta-analysis of over 100 genome wide association studies within the early growth genetics (EGG) consortium, assessing 289 genes encoding mitochondrial function across the ETC and ATP synthase. This study identified 37 SNPs across 32 genes associated with low birthweight. Sanger sequencing was performed to validate the presence of 10 SNPs of interest located within the nuclear genome in a cohort of fetal growth restricted placentas. Of the 10 SNPs assessed we confirmed the presence of 6 in FGR, and identified an additional previously unidentified SNP, and detected 3 nucleotide deletions across numerous components of the electron transport chain. This analysis identified 5 mutations within genes that encode complex I. Subsequent analysis of the gene and proteins that correspond to complex I mutations was performed using PCR, proteomics, and western blotting. Notably, we identified significant downregulation of NDUFA6 at the gene and protein level in FGR, lower protein levels of NDUFS3 and NDUFS6, and confirmed the largest log2 fold change identified in NDUFS6 by western blot. Subsequent investigation of mitochondrial respiratory capacity identified decreased ATP-linked respiration in FGR using Seahorse XF Analysis.

**Conclusion:** Our results identify that there is an inheritable contribution of SNPs within the maternal and paternal nuclear genome in fetal growth restriction. These SNPs alter gene expression and protein abundance of mitochondrial complex I components, impacting structural assembly and subsequent bioenergetic function within the placenta. Collectively these findings suggest that SNPs that affect mitochondrial assembly and function are associated with low birthweight and fetal growth restriction.

## Introduction

Fetal growth restriction (FGR) occurs when the fetus fails to reach its predetermined growth potential. FGR is associated with an increased risk of death *in utero*, neonatal death, and programs lifelong cardiovascular, renal, metabolic diseases and neurodevelopmental delays that persist into adulthood (1). Fetal growth is monitored by *in utero* fetal ultrasound measurements charted against a population growth curve. Clinically, FGR is diagnosed when the estimated fetal weight is <3^rd^ percentile, or <10^th^ percentile with abnormal doppler ultrasound of the umbilical artery waveforms (2, 3). FGR placentas are smaller than healthy term placentas, and display underdeveloped placental villi, characterised by a reduced number of villi with decreased elongation, branching and reduced nutrient and waste exchange capacity (4). In the placentas of pregnancies complicated by FGR, changes have been documented in mitochondrial DNA, copy number, mitophagy pathways, enzyme function, bioenergetics in the form of membrane potential and ATP production, and mitochondrial morphology (5–7). A major role of mitochondria is the generation of energy for cellular function through the production of adenosine triphosphate (ATP). Mitochondria produce ATP through the shuttling of electrons along the electron transport chain (ETC) by a series of four complexes: NADH: ubiquinone oxidoreductase (complex I); succinate dehydrogenase (complex II); ubiquinol-cytochrome c oxidoreductase (complex III); and cytochrome c oxidase (complex IV) (8). The electron transport chain produces an electrochemical gradient that facilitates the generation of ATP via ATP synthase. In the placental trophoblast, in addition to the production of ATP, mitochondria participate in steroid hormone and phospholipid biosynthesis, substrate utilisation, control of calcium, cellular redox and reactive oxygen species signaling (9).

Mitochondria have a unique circular genome that is approximately 16.6kbp long and encodes 1-2% of the proteins required for mitochondrial function. The mitochondrial genome (mtDNA) codes for two ribosomal RNAs, 22 transfer RNAs, and 13 proteins within the ETC (10). Thereby the nuclear genome (nDNA) encodes the remaining 98-99% of structural, ancillary, metabolic and functional proteins of the mitochondria including most of the components of the ETC and ATP synthase. Primary mitochondrial disease refers to a group of disorders characterised by changes in genes found within the mitochondrial or the nuclear genomes which affect mitochondrial function (11). Primary mitochondrial diseases are often characterised by poor growth, learning and developmental delays, cardiac, kidney and liver disease (12–14). This group of diseases includes a subset that occurs as a result of single nucleotide polymorphisms (SNPs) which can alter protein expression within the ETC (15–17). Mutations within both nDNA and mtDNA that encode complex I, complex III, complex IV, and ATP synthase are known causes of mitochondrial diseases (10, 12).

We hypothesised that FGR may be due to SNPs causing dysregulation of nuclear genes encoding proteins that affect mitochondrial function. Building on improvements in genome wide association studies (GWAS), and the availability of summary data from meta-analysis we investigated the association between fetal growth and SNPs within nuclear gene regions that code for components of the ETC and ATP synthase. We identified candidate SNPs as targets for validation within a cohort of FGR placental tissue samples. Within FGR placental tissue samples we confirmed the presence of candidate SNPs affecting ETC components using Sanger sequencing. We then demonstrated that these SNPs were associated with differences in mRNA expression of the affected complex I components using quantitative PCR, and altered protein levels using proteomics and western blotting compared to placentas with normally grown fetuses. These data strongly suggest that SNPs in the nDNA coding for mitochondrial ETC components and ATP synthase generate mitochondrial dysfunction and affect placental trophoblast physiology, contributing to the aetiology of FGR.

## Methods

### Meta-analysis and Discovery Pipeline

An overview of this study is provided in Figure 1. Briefly, Gene Ontology (GO) enrichment was performed on two mitochondrial pathways, the ETC and ATP synthase, identifying 305 target genes. Summary results from the recent GWAS meta-analyses (18) of own birthweight (fetal genetic effect) and offspring birthweight (maternal effect) were examined to identify SNPs within the gene regions regulating these two mitochondrial pathways. Meta-analyses were performed in European-only and Trans-Ancestry (mix ancestry) datasets. Summary results were also available in the European-only cohorts for the fetal effect on own birthweight after adjusting for the correlated maternal genotype, and for maternal effect on offspring birthweight after adjusting for the correlated fetal genotype. Once identified, SNPs within these candidate genes associated with birthweight were filtered for a minor allele frequency ≥5% and multiple testing by Bonferroni correction was applied to identify SNPs for further consideration. SNPs for further consideration were examined by excluding SNPs unique to the trans-ancestry meta-analyses, thereby selecting for SNPs associated with birthweight regardless of ethnicity, before filtering by Linkage Disequilibrium (R^2^ ≤ 0.1) to identify independent SNPs. To examine the potential mechanisms, independent SNPs were mapped to genes before being annotated in the biological context using the GENE2FUNC module in FUMA software (19). This analysis examined the tissue specific gene expression patterns, enrichment of gene sets and associations to well-defined external biological databases. Gene set analyses were performed by assessing the hypergeometric tests of enrichment of the listed genes against MSigDB gene sets (20) including BioCarta, KEGG, Reactome and Gene Ontology. Multiple testing correction was accounted for using the Benjamini-Hochberg method with an adjusted P-value of 0.05 for the gene set analyses. A complete list of gene sets reported for each database is presented in Supplementary Tables 1-16.

**Figure 1:**
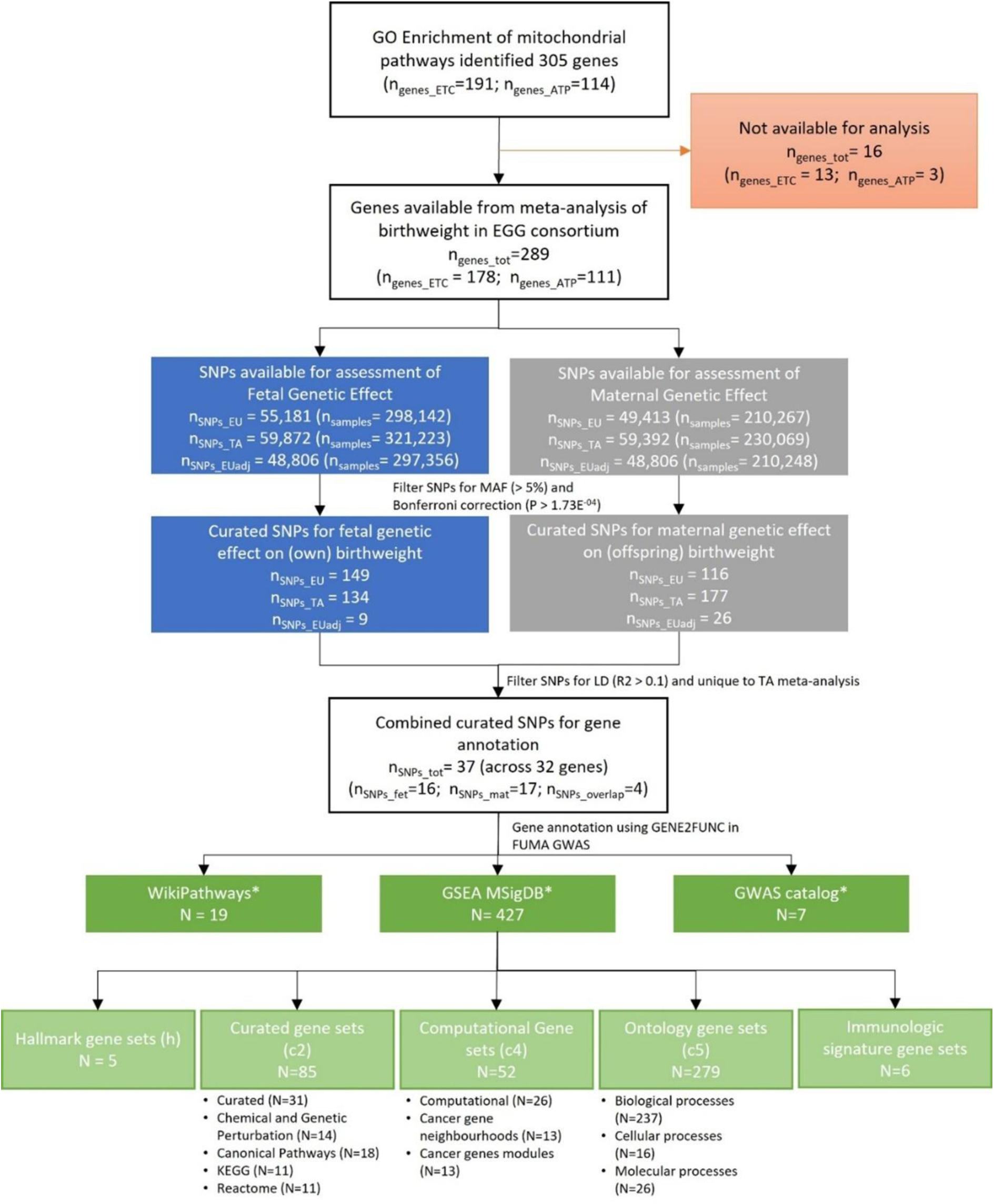
Summarises the results from GO enrichment analysis, the interrogation of the GWAS meta-analysis of birthweight in the EGG consortium and the gene annotation analysis. *A complete list of gene sets reported for each database is presented in Supplementary Tables 1-16. ngenes_tot denotes the total number of genes; ngenes_ETC denotes the number of genes within the ETC; ngenes_ATP denotes the number of genes within ATP synthase; nSNPs_EU denotes the number of SNPs in a meta-analysis of only European study participants; nSNPs_TA denotes the number of SNPs in a meta-analysis of study participants of all ethnicities (trans-ethnic); nSNPs_Euadj denotes the number of SNPs in a meta-analysis of only European study participants adjusting for maternal genotype when examining for fetal effect, and adjusting for fetal genotype when examining for maternal effect; nsamples denotes the number of study participants in the given analysis; nSNPs_tot denotes the total number of SNPs; nSNPS_fet denotes the number of SNPs unique to fetal effect; nSNPS_mat denotes the number of SNPs unique to maternal effect; nSNPs_overlap denotes the number of SNPs overlapping between maternal and fetal effects.

### Ethical Approval, Sample Collection and Preparation

Ethical approval was granted by the University of Newcastle Human Research Ethics Committee (H-382-0602), Hunter New England Health Human Research Ethics Committee (02/06/12/3.13), and Site-Specific Assessment (SSA/15/HNE/291) in compliance with the Declaration of Helsinki standards. Placental samples were collected with informed consent from healthy control pregnancies 37-40 weeks (n = 19), and placental samples with FGR defined by <10^th^ birth centile and abnormal fetal biometry in line with the Delphi consensus (3) (n = 18). Samples were excluded from both healthy and FGR groups if the pregnancy presented with any of the following; premature membrane rupture, hypertension, diabetes, placental abnormalities, chorioamnionitis, immune deficiencies, infectious diseases, smoking, and illicit drug use, to ensure an ideopathic FGR phenotype could be observed and compared.

### Sanger Sequencing

Sanger sequencing and primers used therein were designed, validated, and performed by the Australian Genome Research Facility (AGRF), relevant primers are presented in Table 1. Once performed, results were analysed using Geneious Prime software. Control forward and reverse sequences were trimmed with parameters set at an error probability of 0.05, followed by de novo synthesis. This process was repeated across all control (n=8) samples to facilitate the generation of a consensus strand with a threshold of “Highest Quality” selected at 60% and a Sanger Heterozygotes >50%. The control consensus was then utilised as a reference sequence to identify SNP variance within the placenta of FGR samples (n=10). FGR samples were assessed individually following trimming and de novo synthesis and any difference between the consensus and FGR strands were identified. Once a difference was determined between control and FGR, the corresponding nucleotide sequence was blasted (National Centre for Biotechnology Information, https://blast.ncbi.nlm.nih.gov/) against the human genome reference database (RefSeq Genome Database). SNPs were only considered apparent within FGR if validated against the corresponding gene location when blasted. Upon confirmation, sequences were run in Mutations Taster (https://www.mutationtaster.org/) to predict the severity of the polymorphism, the potential change of splice site or protein feature, as well as if the SNP is a known variant with the 1000 genomes project (TGP).

**Table 1.**
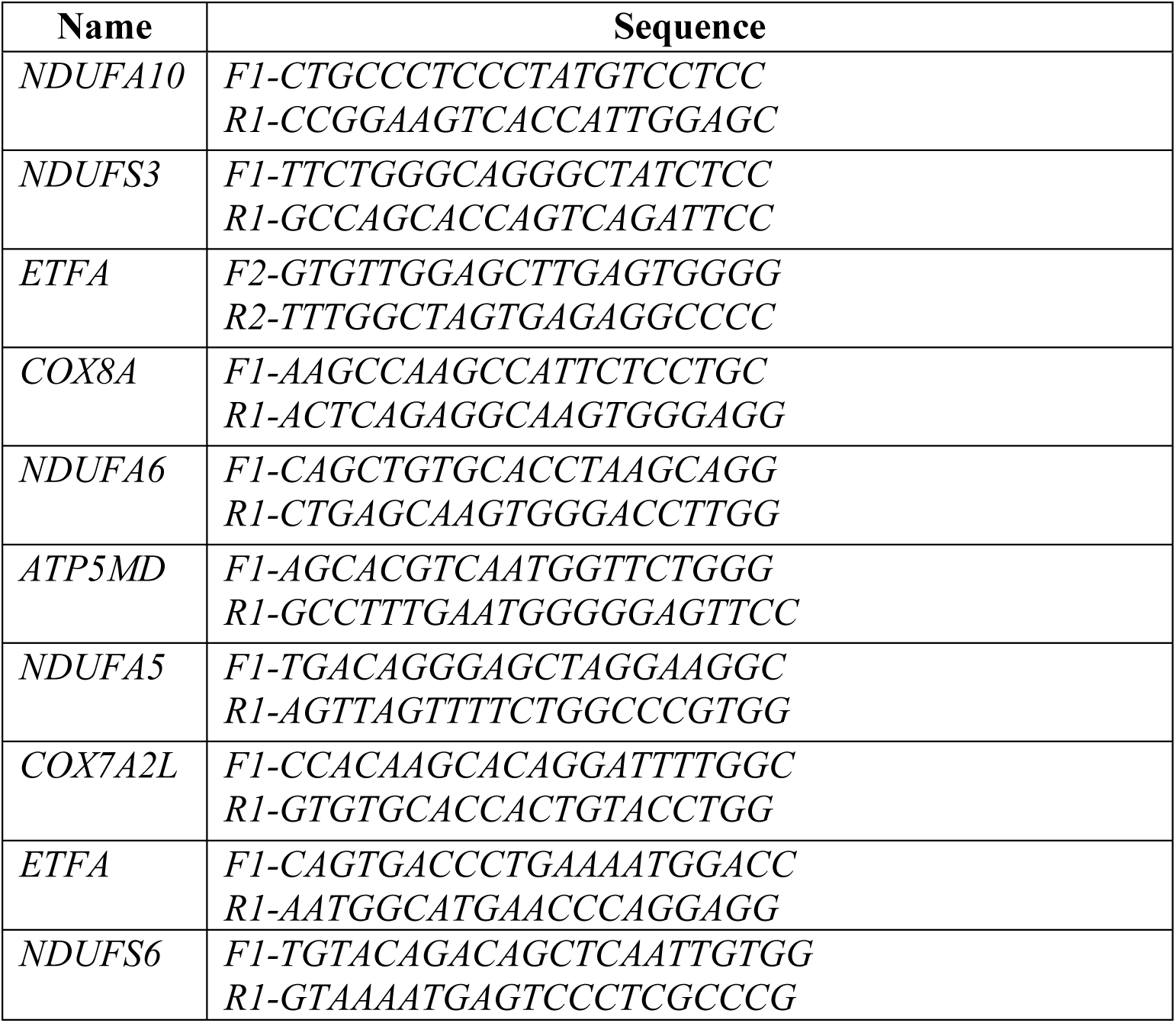
The Primer sequence generated and validated by the Australian Genome Research Facility (AGRF) for use within Sanger sequencing.

### Gene Expression Studies

Following sample collection, RNA was extracted from homogenised placental tissue using the Direct-zol^TM^ RNA miniprep kit (Zymo Research, California, USA) as outlined in the manufacturer’s instruction booklet. RNA quantification was assessed after extraction utilising NanoDrop One (ThermoFisher) and normalised to 1000ng/ul with SuperScript IV (ThermoFisher) reverse transcriptase utilised to synthesise cDNA. RT was performed as per the manufacturer’s specification, 65°C for 5 minutes, 55°C for 10 minutes, 80°C for 10 minutes, before adding E. coli H at 37°C for 20 minutes to remove remaining contaminating RNA. RT-qPCR was conducted using a QuantStudio6 Flex Real-Time PCR machine with the thermocycling parameters: initial activation 95°C for 2 minutes, followed by 40 repeated cycles of denaturation at 95°C for 5 seconds and a combined annealing and extension phase at 60°C for 10 seconds. Predesigned primers were supplied by Sigma-Aldrich (Table 2) in combination with SYBR Green Master Mix (Thermo Fisher Scientific). Relative gene expression was determined following geometric mean normalisation using TOP1 and TBP, with quantification via the 2^-ΔΔCT^ method. All samples were run in duplicate, with no product detected in the non-template controls and a single spike was indicated by melt curve analysis. All CT values over 34 were excluded from analysis to ensure reproducibility of the data in subsequent studies.

**Table 2:**
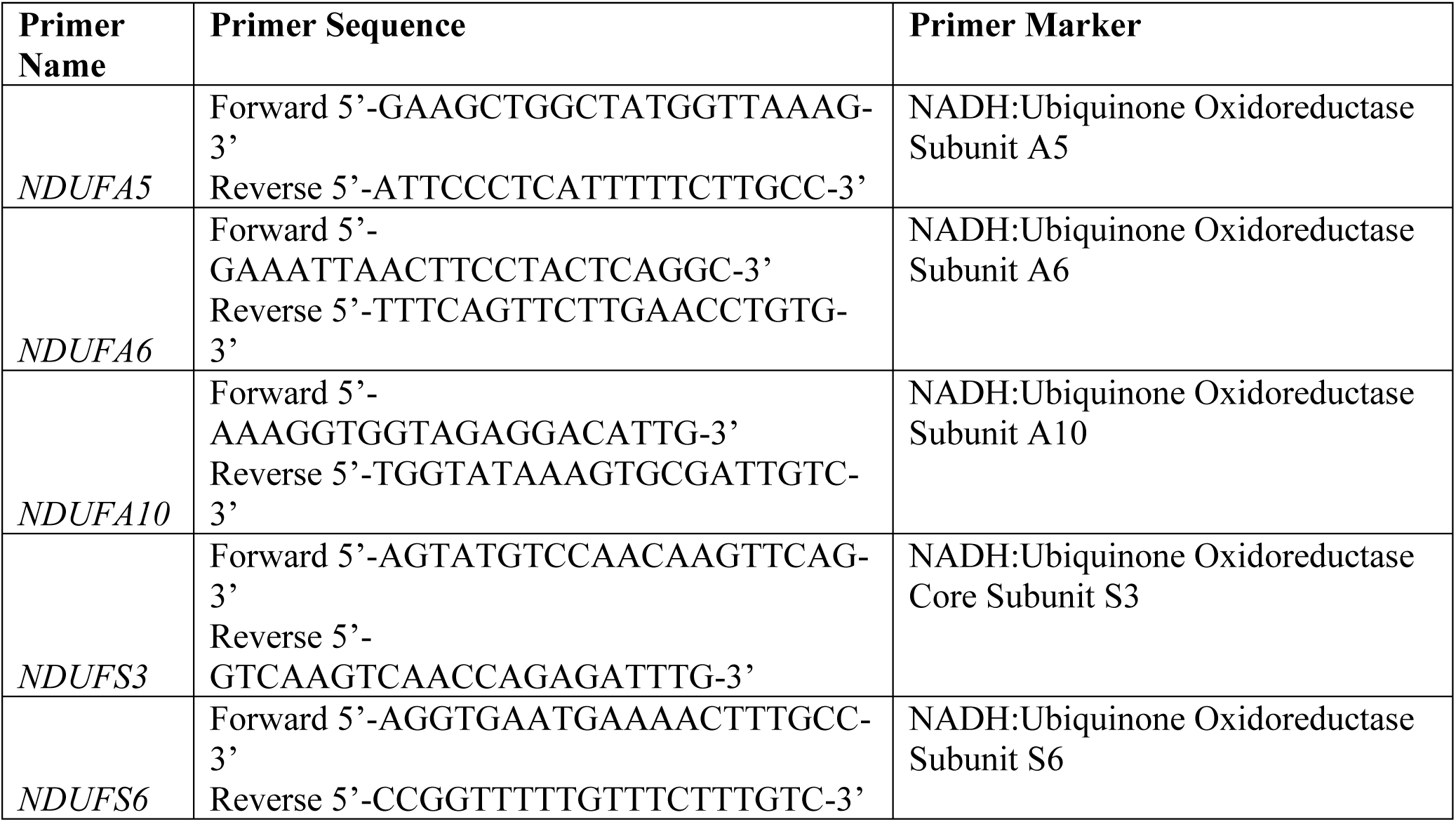
Primer sequences used to investigate complex I subunit gene expression.

### Proteomics

Placental tissue was crushed using a mortar and pestle, with liquid nitrogen and stored at −80°C until mass spectrometry (MS)-based proteomic analysis. Placental tissue samples were resuspended in chilled 4% w/V sodium deoxycholate (SDC), 100 mM Tris HCl (Ph.8.5), boiled at 95°C, for 10 min, and then passed through an 18G blunt needle prior to sonication. Protein concentration was determined by BCA assay, and 200ug of lysate was reduced and alkylated prior to overnight trypsin incubation at 37°C with 1500rpm shaking (1:30 enzyme:protein Trypsin-LysC (V5072; Promega Corporation, Madison, Wisconsin, USA)). Following trypsin digestion, samples were desalted using SDB-RPS Stage Tips. In brief, peptides were diluted to 1:1 with 100 mM Tris-HCl pH 8.5 (SDC < 1%) and vortexed briefly before adding 1:1 of 99% ethylacetate /1% Trifluoroacetic acid (TFA) and thoroughly vortexed for 2 min at 2000rpm using a ThermoMixer. Peptides were then desalted using styrenedivinylbenzene-reverse phase sulfonated StageTips and eluted using 5%NH_4_OH/80% acetonitrile and speed vacuumed to dryness. Lyophilised peptides were reconstituted in 2% acetonitrile/0.3% TFA. Separation of peptides was performed using an Aurora Ultimate C18 column (25 cm × 75 μm, IonOpticks, Collingwood, Australia) at a flow rate of 400nL/min for 90 min on a linear gradient from 3% to 41% of 80% acetonitrile in 0.1% formic acid. Mass spectrometry analysis was conducted on an Eclipse mass spectrometer (Thermo Fisher Scientific) with MS scans acquired with an automatic gain control (AGC) target of 100%, a maximum injection time of 50 ms, and a mass range of 375-1400m/z at a resolution of 60,000. MS/MS was performed at a resolution of 15,000, with normalised collision energy of 30%, and dynamic maximum injection time. Raw data was analysed using Proteome Discoverer 2.5 (Thermo Fisher) as previously published (21). We examined only proteins within complex I with a log2 fold change to determine directionality and abundance change as previously published (22).

### Western Blotting

Tissue was homogenised utilising the Precellys 24 (Bertin instruments) in RIPA buffer (50 mM Tris-HCl, 150 mM NaCl, 1 mM EDTA, 1% NP-40, 0.5% sodium deoxycholate, 100 nM sodium orthovanadate and Complete Mini Protease Inhibitor Cocktail tablets (Roche Diagnostics, North Ryde, Australia) and 1 nM PMSF). Samples were vortexed at 5000rpm in two 30 second bursts, prior to centrifugation at 13,000rpm at 4°C for 10 min and the supernatant was collected for western blotting and stored in a −80°C freezer until subsequent examination. Isolation of enriched mitochondrial populations was performed in accordance with previous publications (23). Briefly, tissue was homogenised in a glass dounce with 1 ml of chilled isolation media (250 mM sucrose, 0.5 mM Na2EDTA, 10 mM Tris, pH 7.4) before centrifugation at 1,500g for 10 mins at 4°C, supernatant was collected and spun at 4,000g for 15 mins to produce a cytotrophoblast mitochondrial pellet (Cyto-Mito), with the supernatant centrifuged again at 12,000g to produce a syncytiotrophoblast enriched mitochondrial pellet (Syncytio-Mito). Pellets were resuspended in either RIPA buffer for western blotting, isolation media for subsequent mitochondrial analysis.

Control and FGR placental tissue lysates were prepared, with concentrations normalised to 10 µg/µl via Pierce BCA Assay Kit (Thermo Fisher Scientific). Samples were loaded onto 12% NuPage Gels, and subsequently transferred to Polyvinylidene fluoride (PVDF) membrane. Following transfer, the membranes were blocked in Odyssey blocking buffer (LI-COR Bioscience) solution for 1 hour. The membranes were then left to incubate overnight with a primary antibody, targeting the protein interest. The following antibodies were used to target the proteins of interest, anti-NDUFS6 (1:500 ab195807) and, β actin (ACTB ab8227) at 1:1000. The following day, the membranes were washed extensively in PBS/Tween (PBST), before being incubated for 1 hour in the appropriate secondary antibody anti-rabbit (IRDye 680 Donkey, Licor, Lincoln, NE, USA), or anti-mouse (IRDye 680 goat, Licor, Lincoln, NE, USA) at 1:20,000 dilution. The membranes were washed in PBST before being imaged via infrared using a LI-COR Odyssey CLx imaging system and quantified via Image Studio v 5.2.

### Mitochondrial Function

To assess mitochondrial respiration Seahorse XF Analysis (Agilent Technologies, USA) was performed. To ensure functionality, enriched mitochondrial isolates were extracted as previously published (23). To facilitate assessment of isolated mitochondria, samples were suspended in mitochondrial isolation buffer and added into the plate in duplicate and spun at 200g for 10 minutes at 4°C. 130μl of pre warmed (37°C) mitochondrial assay solution (Sucrose 70mM, Mannitol 22nM, KH_2_PO_4_ 10mM, MgCl_2_ 5mM, HEPES 2mM, EGTA 1mM, Glutamate 10mM, Malate 5mM, Pyruvate 5mM, ADP 4mM adjusted to pH 7.4) were added and the plate was loaded following calibration in accordance to the manufacturer’s instructions in calibration media. The inclusion of glutamate, malate, and pyruvate to the media and ADP through Injection port A primed the mitochondria for subsequent steps ensuring no rate limiting with uncoupling. Port B administered FCCP at 40μM allowing maximum respiration (ATP linked respiration) to be assessed. Port C and D administered Rotenone at 0.5μM and Antimycin A at 5mM to inhibit complex I and III respectively, allowing residual oxygen concentration (non-mitochondrial respiration) to be calculated. All measurements were normalised to protein loaded as per previous publications, and non-mitochondrial respiration subtracted to exclude all non-mitochondrial oxygen consumption (23).

### Statistical Analysis for gene and protein examination

Statistical analysis for gene expression and western blotting applied Grubb’s test and Route testing to ensure no outliers were present within the dataset. All p values were compared with an alpha value of 0.05 (α = 0.05) and expressed as mean ±SD. Student’s t-tests were performed between groups to establish significance between healthy and FGR placentae. A two-way ANOVA was performed for further analysis of mitochondria between the two cell lineages in addition to comparing control and FGR groups. The Šídák post hoc analysis method was chosen to best examine across the multiple groups.

## Results

### Single Nucleotide Polymorphisms are associated with birthweight

Meta-analyses on data from 321,223 individuals within the Early Growth Genetic (EGG) consortium (http://egg-consortium.org/) have been published identifying SNPs that are linked to birthweight centiles (24). We interrogated this public repository for SNPs present in genes known to encode proteins that affect mitochondrial structure and function. GO enrichment of mitochondrial pathways identified 305 targets (178 from the electron transport chain and 111 from ATP synthase) of which data were available for 289 genes across all summary datasets. The analysis of own birthweight (effect of fetal genetics) SNP data was available for three groups of genetic origin within the database 55,181 European-only individuals, 59,872 Trans-Ancestry individuals, and 48,806 individuals to evaluate fetal effect adjusting for the correlated maternal genotype. After filtering for allelic frequency and applying a Bonferroni correction, 149, 134 and 9 SNPs linked to birthweight centiles and affecting ETC and ATP synthase genes were identified from the three respective datasets. In the meta-analysis of maternal contribution to offspring birthweight, SNP data was available from summary results for 49,413 European-only, 59,392 Trans-Ancestry, and 48,806 SNPs adjusting for correlated fetal genotype. Upon filtering for allelic frequency and Bonferroni correction, 116, 177 and 26 SNPs from each respective dataset remained. After filtering for linkage disequilibrium and excluding SNPs unique to a single meta-analyses of own birthweight and offspring birthweight, thereby selecting for SNPs associated with birthweight regardless of ethnicity, a total of 37 SNPs across 32 genes remained significant across the six datasets (summarised in Figure 1). From these 37 SNPs, four SNPs overlapped between the meta-analyses of own birthweight (fetal effect) and offspring birthweight (maternal effect), 17 SNPs were unique to the meta-analysis of offspring birthweight, and 16 SNPs were unique to the meta-analysis of own birthweight (Table 3). The mitochondrial gene regions affecting ETC and ATP synthase did not overlap with SNPs previously associated with decreased birthweight (supplementary Figure 1).

**Table 3:**
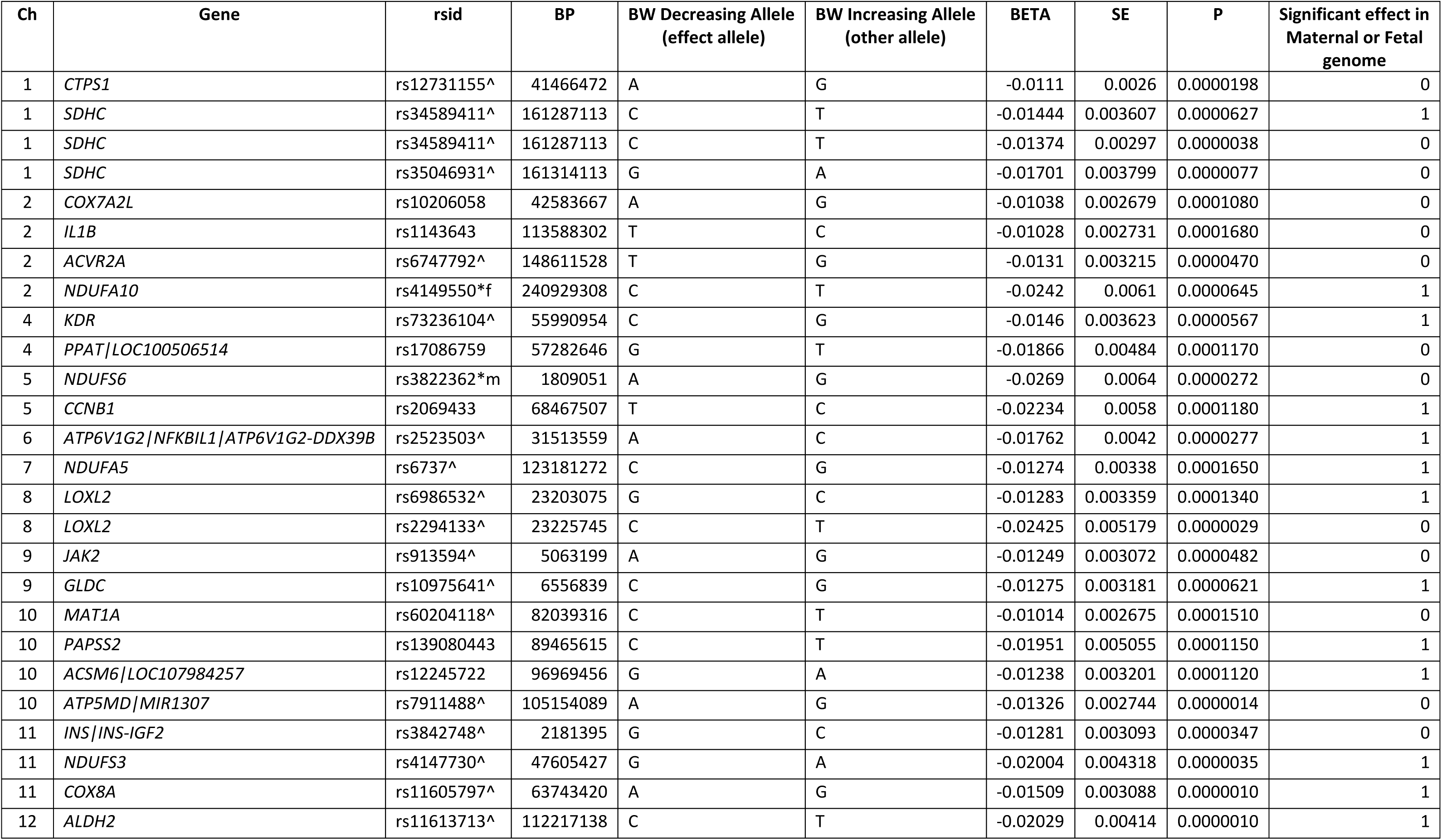
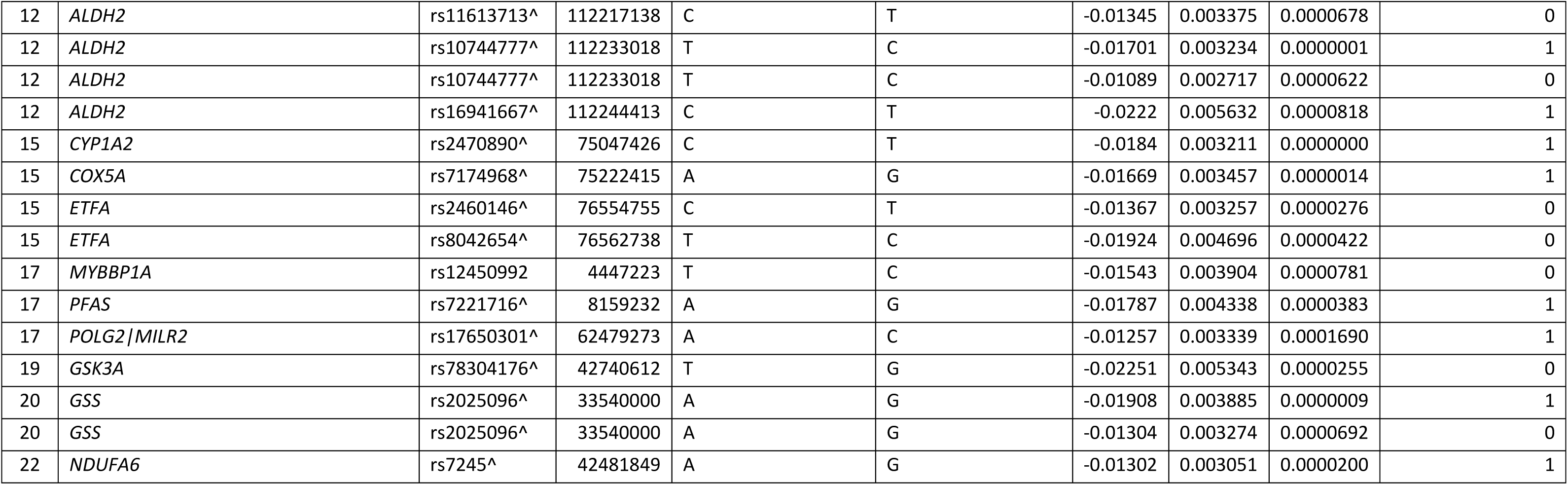
Presents all SNPs found to be significantly associated with birthweight, the chromosomes (Ch) they are present on, the location as a reference SNP (rsid), the gene which corresponds to the gene region and known aliases (Gene), the significance level of the association (P), the size of the gene region (BP) and effect allele associated with an increase or decrease in birthweight (BW decreasing, increasing allele). SE and Beta values are presented in their respective columns. Significant maternal effect or significant fetal effect (1 =Maternal 0=Fetal genome). Significance was established at p < 0.000173 (0.05/289 independent regions). ^ denotes SNP is significant in Trans-ethnic meta-analysis and European-ancestry meta-analysis. *m denotes SNP remaining significant after adjusting for maternal SNP. *f denotes SNP remain significant after adjusting for fetal SNP.

| Ch | Gene | rsid | BP | BW Decreasing Allele<br>(effect allele) | BW Increasing Allele<br>(other allele) | BETA | SE | P | Significant effect in<br>Maternal or Fetal<br>genome |
| --- | --- | --- | --- | --- | --- | --- | --- | --- | --- |
| 1 | *CTPS1* | rs12731155^ | 41466472 | A | G | -0.0111 | 0.0026 | 0.0000198 | 0 |
| 1 | *SDHC* | rs34589411^ | 161287113 | C | T | -0.01444 | 0.003607 | 0.0000627 | 1 |
| 1 | *SDHC* | rs34589411^ | 161287113 | C | T | -0.01374 | 0.00297 | 0.0000038 | 0 |
| 1 | *SDHC* | rs35046931^ | 161314113 | G | A | -0.01701 | 0.003799 | 0.0000077 | 0 |
| 2 | *COX7A2L* | rs10206058 | 42583667 | A | G | -0.01038 | 0.002679 | 0.0001080 | 0 |
| 2 | *IL1B* | rs1143643 | 113588302 | T | C | -0.01028 | 0.002731 | 0.0001680 | 0 |
| 2 | *ACVR2A* | rs6747792^ | 148611528 | T | G | -0.0131 | 0.003215 | 0.0000470 | 0 |
| 2 | *NDUFA10* | rs4149550*f | 240929308 | C | T | -0.0242 | 0.0061 | 0.0000645 | 1 |
| 4 | *KDR* | rs73236104^ | 55990954 | C | G | -0.0146 | 0.003623 | 0.0000567 | 1 |
| 4 | *PPAT/LOC100506514* | rs17086759 | 57282646 | G | T | -0.01866 | 0.00484 | 0.0001170 | 0 |
| 5 | *NDUFS6* | rs3822362*m | 1809051 | A | G | -0.0269 | 0.0064 | 0.0000272 | 0 |
| 5 | *CCNB1* | rs2069433 | 68467507 | T | C | -0.02234 | 0.0058 | 0.0001180 | 1 |
| 6 | *ATP6V1G2/NFKBIL1/ATP6V1G2-DDX39B* | rs2523503^ | 31513559 | A | C | -0.01762 | 0.0042 | 0.0000277 | 1 |
| 7 | *NDUFA5* | rs6737^ | 123181272 | C | G | -0.01274 | 0.00338 | 0.0001650 | 1 |
| 8 | *LOXL2* | rs6986532^ | 23203075 | G | C | -0.01283 | 0.003359 | 0.0001340 | 1 |
| 8 | *LOXL2* | rs2294133^ | 23225745 | C | T | -0.02425 | 0.005179 | 0.0000029 | 0 |
| 9 | *JAK2* | rs913594^ | 5063199 | A | G | -0.01249 | 0.003072 | 0.0000482 | 0 |
| 9 | *GLDC* | rs10975641^ | 6556839 | C | G | -0.01275 | 0.003181 | 0.0000621 | 1 |
| 10 | *MAT1A* | rs60204118^ | 82039316 | C | T | -0.01014 | 0.002675 | 0.0001510 | 0 |
| 10 | *PAPSS2* | rs139080443 | 89465615 | C | T | -0.01951 | 0.005055 | 0.0001150 | 1 |
| 10 | *ACSM6/LOC107984257* | rs12245722 | 96969456 | G | A | -0.01238 | 0.003201 | 0.0001120 | 1 |
| 10 | *ATP5MD/MIR1307* | rs7911488^ | 105154089 | A | G | -0.01326 | 0.002744 | 0.0000014 | 0 |
| 11 | *INS/INS-IGF2* | rs3842748^ | 2181395 | G | C | -0.01281 | 0.003093 | 0.0000347 | 0 |
| 11 | *NDUFS3* | rs4147730^ | 47605427 | G | A | -0.02004 | 0.004318 | 0.0000035 | 1 |
| 11 | *COX8A* | rs11605797^ | 63743420 | A | G | -0.01509 | 0.003088 | 0.0000010 | 1 |
| 12 | *ALDH2* | rs11613713^ | 112217138 | C | T | -0.02029 | 0.00414 | 0.0000010 | 1 |
| 12 | *ALDH2* | rs11613713^ | 112217138 | C | T | -0.01345 | 0.003375 | 0.0000678 | 0 |
| 12 | *ALDH2* | rs10744777^ | 112233018 | T | C | -0.01701 | 0.003234 | 0.0000001 | 1 |
| 12 | *ALDH2* | rs10744777^ | 112233018 | T | C | -0.01089 | 0.002717 | 0.0000622 | 0 |
| 12 | *ALDH2* | rs16941667^ | 112244413 | C | T | -0.0222 | 0.005632 | 0.0000818 | 1 |
| 15 | *CYP1A2* | rs2470890^ | 75047426 | C | T | -0.0184 | 0.003211 | 0.0000000 | 1 |
| 15 | *COX5A* | rs7174968^ | 75222415 | A | G | -0.01669 | 0.003457 | 0.0000014 | 1 |
| 15 | *ETFA* | rs2460146^ | 76554755 | C | T | -0.01367 | 0.003257 | 0.0000276 | 0 |
| 15 | *ETFA* | rs8042654^ | 76562738 | T | C | -0.01924 | 0.004696 | 0.0000422 | 0 |
| 17 | *MYBBP1A* | rs12450992 | 4447223 | T | C | -0.01543 | 0.003904 | 0.0000781 | 0 |
| 17 | *PFAS* | rs7221716^ | 8159232 | A | G | -0.01787 | 0.004338 | 0.0000383 | 1 |
| 17 | *POLG2/MILR2* | rs17650301^ | 62479273 | A | C | -0.01257 | 0.003339 | 0.0001690 | 1 |
| 19 | *GSK3A* | rs78304176^ | 42740612 | T | G | -0.02251 | 0.005343 | 0.0000255 | 0 |
| 20 | *GSS* | rs2025096^ | 33540000 | A | G | -0.01908 | 0.003885 | 0.0000009 | 1 |
| 20 | *GSS* | rs2025096^ | 33540000 | A | G | -0.01304 | 0.003274 | 0.0000692 | 0 |
| 22 | *NDUFA6* | rs7245^ | 42481849 | A | G | -0.01302 | 0.003051 | 0.0000200 | 1 |

### Presence of polymorphisms in fetal growth restriction

Following the identification of SNPs associated with birthweight, a cohort of placental samples from healthy control and FGR pregnancies was collected for the validation of polymorphisms associated with FGR. Clinical characteristics are presented in Table 4. When assessed using fetal measurements and biometry the FGR group were born significantly smaller with decreased placental weight, abdominal circumference, femur length, and head circumference compared to term controls (Table 2). No significant difference was observed between maternal age or BMI of the mothers in the two groups. A significant difference was observed based on gestation at delivery, with three FGR collected <32 weeks.

**Table 4:**
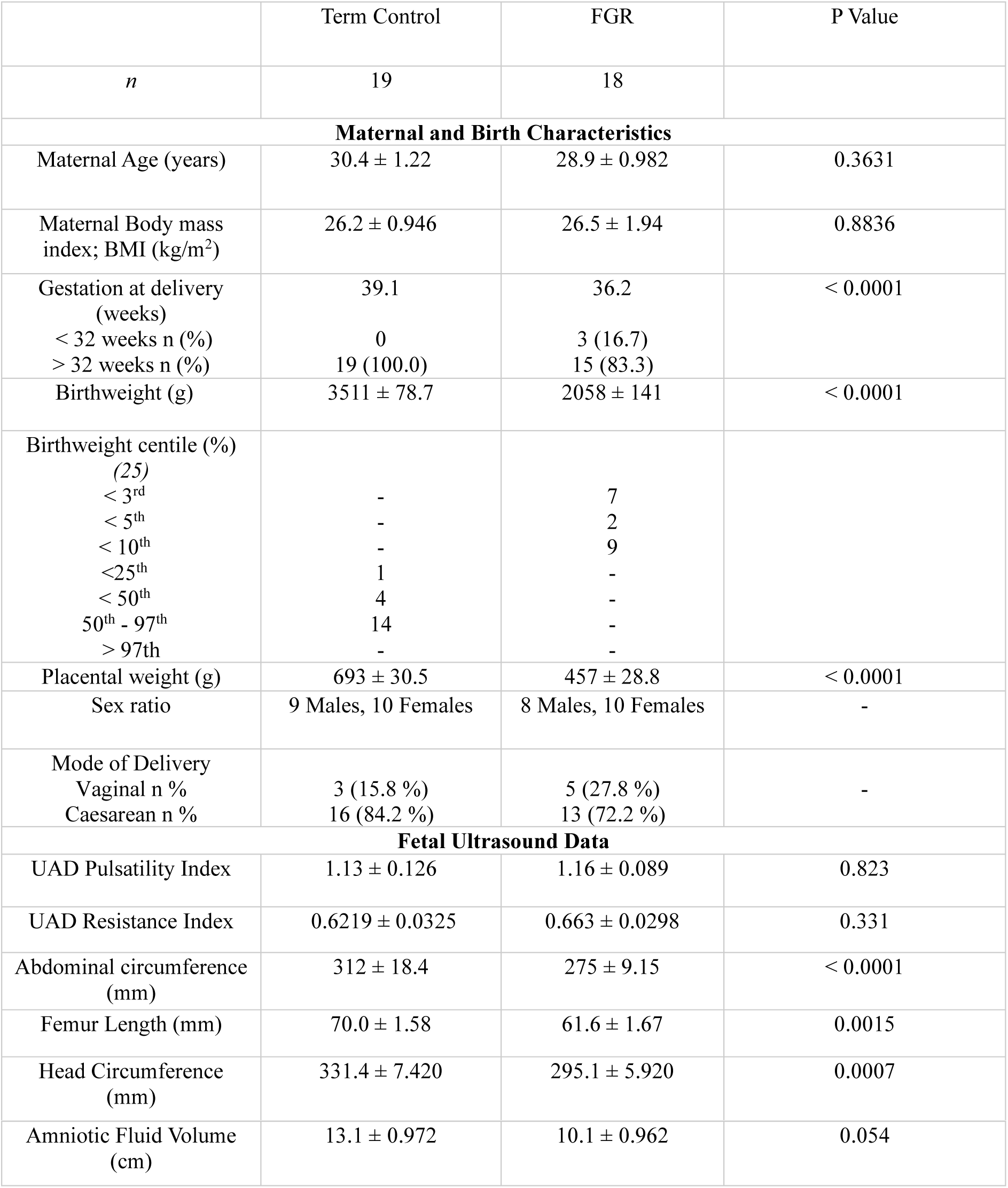
Clinical characteristics of the samples.

### Validation of single nucleotide polymorphisms in FGR placentas

To confirm if the SNPs identified to be associated with birthweight within our GWAS study were present within our cohort of FGR placentas (n=10) targeted Sanger sequencing was performed. SNPs were deemed present in a FGR sample if they resulted in, or changed from, the identified effect allele at the corresponding rsid location determined by our GWAS analysis. Of the 10 SNPs assessed across 9 genes, six were identified within FGR placentas (Table 5), with three of the SNPs (NDUFS6 - rs3822362 (Figure 2), NDUFA5 - rs6737, COX8A - rs11605797) found at a disproportionally greater percentage compared to their expected population prevalence on gnomAD. The remaining three were found at expected (NDUFS3 - rs4147730) or underrepresented (COX7A2L - rs10206058, ETFA - rs8042654) levels within our FGR placentas. Subsequent assessment of the pathogenicity of the SNPs was assessed in silico using Mutation Taster (https://www.mutationtaster.org/), with three of the SNPs (NDUFS6, NDUFA5, and COX8A) predicted to change splice sites in NDUFS6, NDUFA5, and COX8A allele changes. Further, COX8A SNPs were predicted to result in protein feature changes.

**Figure 2:**
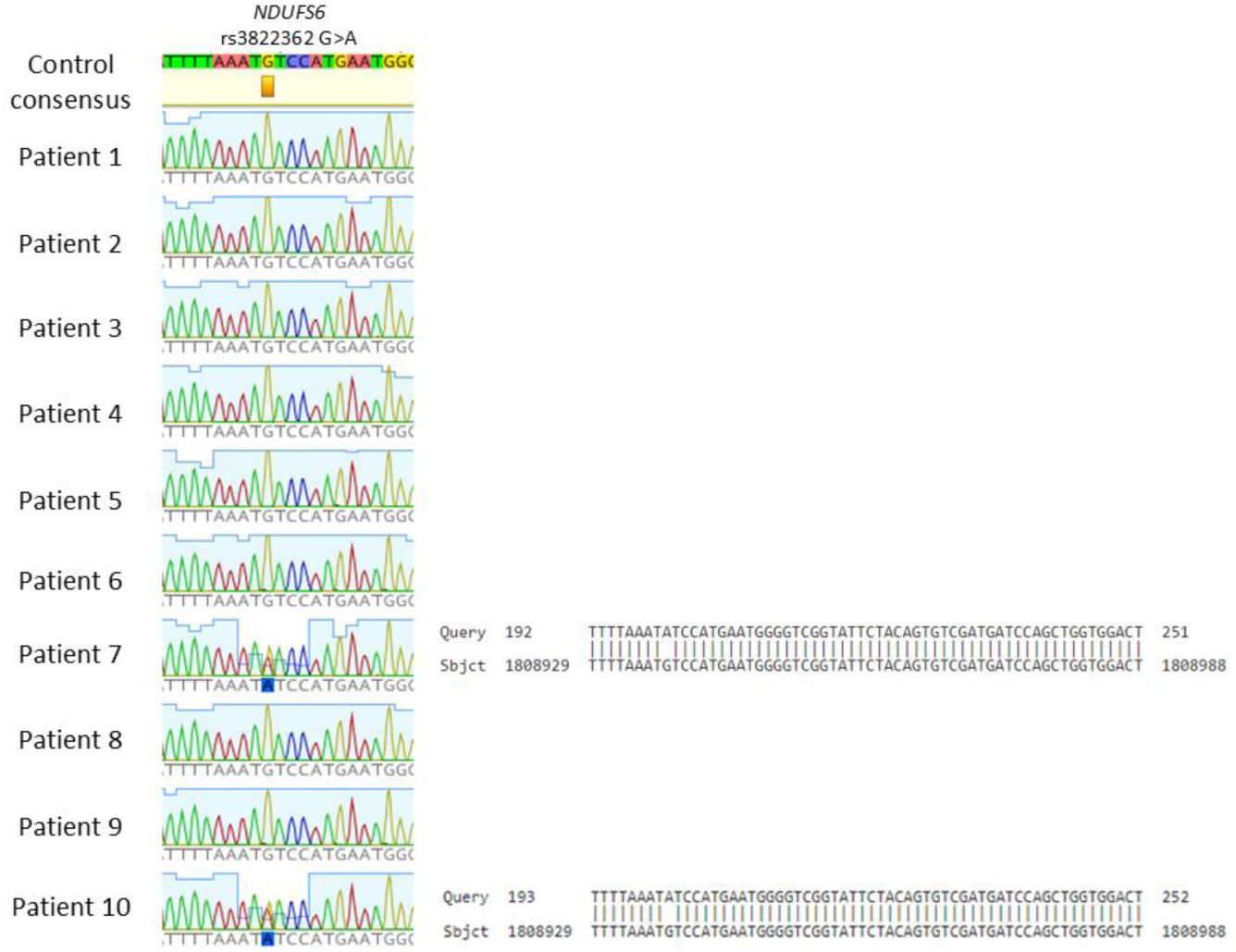
Shows the Sanger sequencing results in which a G>A SNP is present at rs3822362 in 2 out of 10 FGR placentae when run against a consensus strand from n=8 healthy term control placentas. Once identified, SNPs were validated against the NCBI reference genome sequence ID NC_000005.10 to determine the location of the SNP.

**Table 5:**
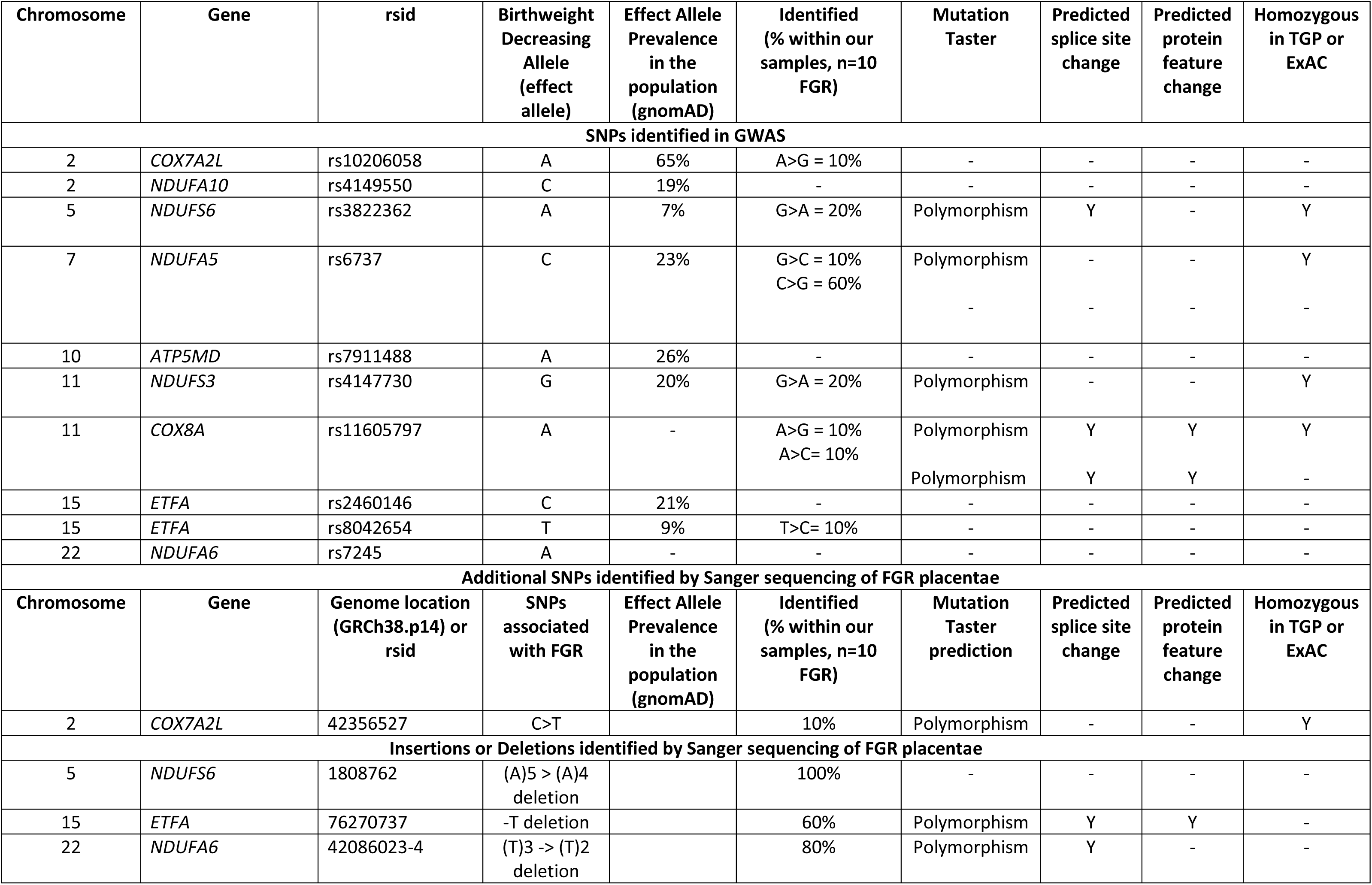
Shows the prevalence of SNPs associated with low birthweight in a primary cohort of placentas collected from fetal growth restricted pregnancies. This table highlights the corresponding chromosome of SNPs, the location within the genome with reference to their respective rsid or GRCH38, the known occurrence within the population (genomAD), effect alleles identified in our GWAS study to be associated with low birthweight, directional shifts in alleles, and the percentage of our primary FGR samples (n=10) that were identified with the SNP. In addition, table 2 shows novel SNPs and deletions identified within the same gene regions as our GWAS SNPs of interest, with Mutation Taster used to assess variant pathogenicity, predicted splice site and protein changes in silico. Further information regarding these classifications can be found at https://www.mutationtaster.org/. Y denotes a predicted change in the associated column, no predicted change or an undetermined value in the FGR samples or Mutation Taster is represented by a dash (-). The table addresses both known SNPs found in our GWAS, and novel SNPs identified in the gene regions of interest by Sanger Sequencing.

| Chromosome | Gene | rsid | Birthweight Decreasing Allele (effect allele) | Effect Allele Prevalence in the population (gnomAD) | Identified (% within our samples, n=10 FGR) | Mutation Taster | Predicted splice site change | Predicted protein feature change | Homozygous in TGP or ExAC |
| --- | --- | --- | --- | --- | --- | --- | --- | --- | --- |
| SNPs identified in GWAS |  |  |  |  |  |  |  |  |  |
| 2 | COX7A2L | rs10206058 | A | 65% | A>G = 10% | - | - | - | - |
| 2 | NDUFA10 | rs4149550 | C | 19% | - | - | - | - | - |
| 5 | NDUFS6 | rs3822362 | A | 7% | G>A = 20% | Polymorphism | Y | - | Y |
| 7 | NDUFA5 | rs6737 | C | 23% | G>C = 10%<br>C>G = 60% | Polymorphism<br>- | -<br>- | -<br>- | Y<br>- |
| 10 | ATP5MD | rs7911488 | A | 26% | - | - | - | - | - |
| 11 | NDUFS3 | rs4147730 | G | 20% | G>A = 20% | Polymorphism | - | - | Y |
| 11 | COX8A | rs11605797 | A | - | A>G = 10%<br>A>C = 10% | Polymorphism<br>Polymorphism | Y<br>Y | Y<br>Y | Y<br>- |
| 15 | ETFA | rs2460146 | C | 21% | - | - | - | - | - |
| 15 | ETFA | rs8042654 | T | 9% | T>C = 10% | - | - | - | - |
| 22 | NDUFA6 | rs7245 | A | - | - | - | - | - | - |
| Additional SNPs identified by Sanger sequencing of FGR placentae |  |  |  |  |  |  |  |  |  |
| Chromosome | Gene | Genome location (GRCh38.p14) or rsid | SNPs associated with FGR | Effect Allele Prevalence in the population (gnomAD) | Identified (% within our samples, n=10 FGR) | Mutation Taster prediction | Predicted splice site change | Predicted protein feature change | Homozygous in TGP or ExAC |
| 2 | COX7A2L | 42356527 | C>T |  | 10% | Polymorphism | - | - | Y |
| Insertions or Deletions identified by Sanger sequencing of FGR placentae |  |  |  |  |  |  |  |  |  |
| 5 | NDUFS6 | 1808762 | (A)5 > (A)4 deletion |  | 100% | - | - | - | - |
| 15 | ETFA | 76270737 | -T deletion |  | 60% | Polymorphism | Y | Y | - |
| 22 | NDUFA6 | 42086023-4 | (T)3 -> (T)2<br>deletion |  | 80% | Polymorphism | Y | - | - |

### Novel identification of single nucleotide polymorphisms in FGR placentas

In addition to validating SNPs identified in our GWAS analysis in a primary cohort of FGR, several allele changes were identified by Sanger sequencing in our FGR samples within the same gene region (Table 5). COX7A2L - 42356527 C > T, was identified to be present in one of our FGR samples and predicted by Mutation Taster as “polymorphisms”. Further, our study identified three deletions present in FGR placentas. ETFA – 76270737 was identified in 60% of our FGR samples with predicted splice site and protein feature changes. Four of our FGR samples contained a (T)3 -> (T)2 deletion at NDUFA6 - 42086023 while a further four contained the same SNP at NDUFA6 - 42086024, with a predicted “polymorphism” likely to result in splice site changes in 80% of our FGR. The remaining (A)5 > (A)4 deletion was identified in 100% of our FGR samples NDUFS6 – 1808762 but did not return any predictions when run through Mutation Taster.

### Gene and protein changes in FGR placentas

Given the disproportionate occurrence of SNPs, both identified in our GWAS to be associated with low birthweight, and confirmed in our Sanger sequencing of FGR placentas located within complex I, we investigated the transcriptional and translational changes that may arise in the genes and proteins associated with our identified complex I SNPs. We assessed gene expression of FGR samples relative to control placentas and observed significant increases in NDUFA5, a significant decrease in NDUFA6 and no change in NDUFA10 (Figure 3 A, B, C). No change was observed in NDUFS3 or NDUFS6 (Figure 3 D, E). Gene expression was not influenced by the severity of FGR within our group. Proteomics analysis was performed on placentas from five FGR samples <3^rd^ centile, and compared to four control placentas. Control samples were used as a baseline “normal” to establish a ratio based on average peptide hits and transformed to show directionality by a log2ratio. A positive log2 fold change (yellow) represents a protein that is higher in controls and decreased in the FGR group. Conversely, a negative value log2 fold change (blue) represents a protein that is less abundant in the control samples and increased in FGR. Our observations of increased gene expression of NDUFA5 and decreased NDUFA6 in FGR were conserved when assessing protein levels (Figure 3F). Similarly, no change was observed in NDUFS10 at a protein level, while NDUFS3 and NDUFS6 protein levels were greater in healthy controls than FGR placentae (Figure 3F). Given NDUFS6 showed the greatest change at the protein level between our groups, subsequent western blot validation was performed in whole tissue (Figure 3G) and in isolated mitochondria from two trophoblast lineages, cytotrophoblast and syncytiotrophoblast (Figure 3H). Western blot analysis confirmed our proteomics results showing that NDUFS6 was significantly reduced in FGR. This finding was conserved when assessing isolated mitochondria from the cytotrophoblast (Cyto-Mito) and syncytiotrophoblast (Syncytio-Mito). Expectedly, given previously published differences between these mitochondrial subtypes (22, 23), we identified a decline in NDUFS6 in healthy controls between the Cyto-Mito and Syncytio-Mito (Figure 3H), although this was not present in FGR. When assessing the functional implication of these structural changes we observed significantly reduced capacity to produce ATP in FGR tissue compared to controls (Figure 3I). Expectedly we observed a significant reduction in both the control and FGR syncytiotrophoblast mitochondria relative to the cytotrophoblast population.

**Figure 3:**
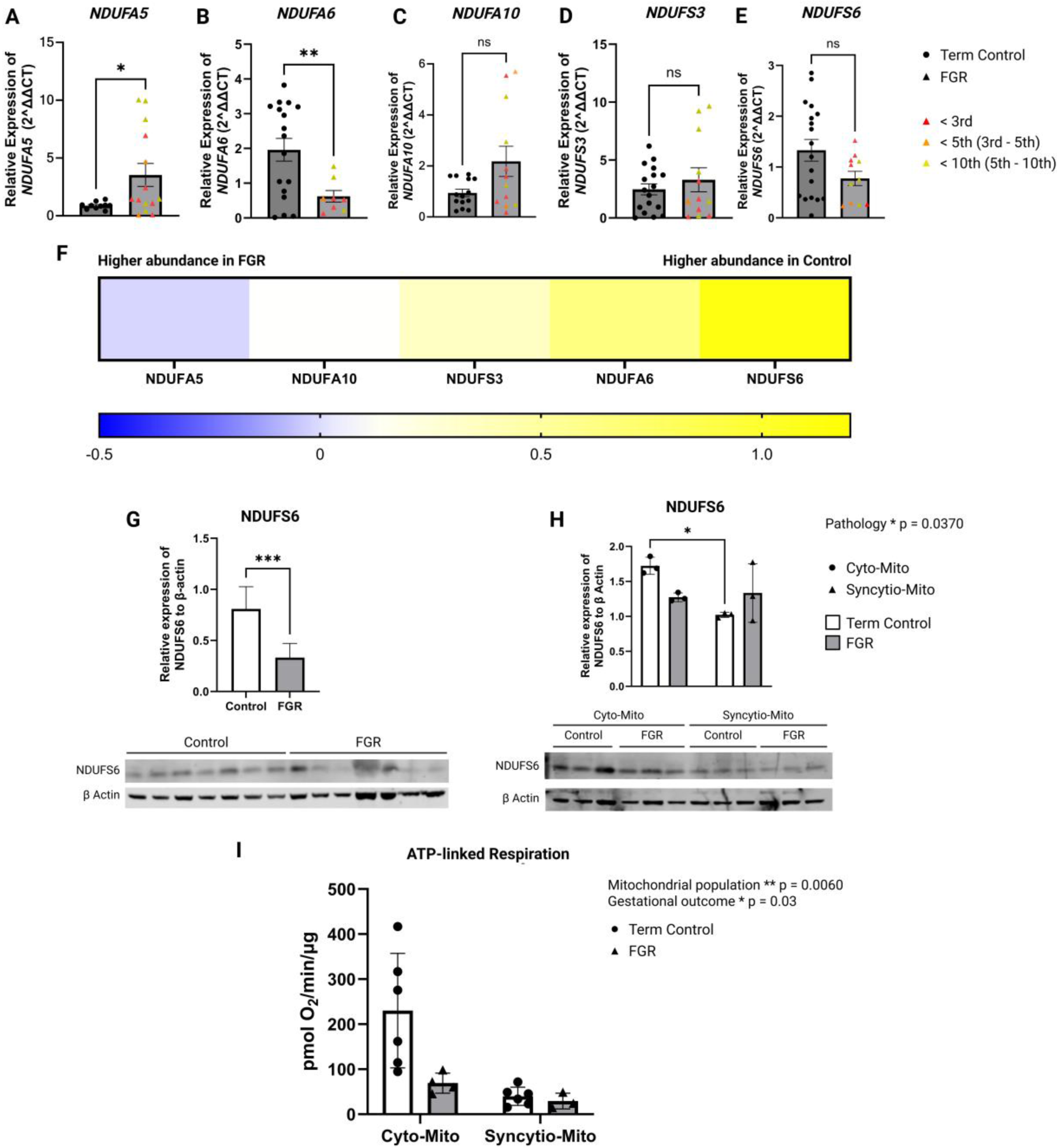
Shows the relative change in gene expression assessed by PCR across complex I subunits. n = 10C – 14 F (A), 17C – 8F (B), 14C – 12F (C), 17C – 12 (D), 18C-11F (E) 18C-11F. Controls shown as circles on the graph, FGR samples depicted as triangles. Variance in sample size occurred due to exclusion of samples with cycle threshold (CT) values in excess of 34 in accordance with SYBR green manufacturers optimum conditions to ensure reproducibility of the data and accurate interpretation across all primers. FGR samples are visually separated to illustrate a range of pathology with less than the 3^rd^ centile shown as red, 3^rd^-5^th^ shown as a dark orange, and 5^th^ – 10^th^ represented as yellow (no significance or linearity was observed when stratifying by birthweight centile). Proteomics data is shown as log2 fold change (F), with a positive number (yellow) representing a greater abundance in control samples. Conversely a negative number (blue) represents a greater abundance in FGR. Proteins that did not express a change are represented in (white). Western blotting (G) was used on a spread of FGR samples (<3^rd^-<10^th^ centile) to validate our proteomics findings and assess whether the overall effect was maintained across a broad range of FGR severity. Further analysis was performed (H) to assess isolated mitochondria from the cytotrophoblast (Cyto-Mito) represented as circles, and syncytiotrophoblast mitochondria (Syncytio-Mito) represented by triangles, in both control (white) and FGR (grey). (I) Functional assessment of mitochondrial maximum respiratory capacity (ATP linked respiration) of healthy term placenta in white and FGR in grey. Data are expressed as mean ±SD. Statistical analysis was performed by students t-test, 2-way ANOVA with Šídák post hoc analysis. *p < 0.05, **p < 0.01, ***p < 0.001, ****p < 0.0001.

## Discussion

We examined a meta-analysis of more than 100 study cohorts contained within the EGG consortium targeting 289 genes encoding mitochondrial function. Upon curation, we determined that 372 SNPs were associated with fetal birthweight, with subsequent filtering identifying 37 SNPs across 32 genes and 16 chromosomes that reached significance (Table 3). Sanger sequencing confirmed the presence of six SNPs identified in our GWAS analysis in a new cohort of FGR placentas. We discovered one additional SNP and three deletions within the same gene regions not previously associated with disease. All confirmed SNPs had the predicted potential to alter splice sites, change protein features, or both, with the exception of NDUFS6 – 1808762, which was, however, identified in all of our FGR placental samples. Given the disproportional number of SNPs identified in complex I by our GWAS analysis and confirmed by Sanger sequencing, we conducted an investigation of complex I subunits by PCR, proteomics, and western blotting, which demonstrated significant alterations in gene expression and protein levels in numerous subunits. All of the genes presented in this study are encoded by the nDNA, and our results suggest that the presence of SNPs within the parental genome has the potential to affect mitochondrial function, and subsequently affect placental, and thereby fetal development, supporting a role for mitochondrial nuclear genetics in the aetiology of fetal growth restriction.

### Structural implications of single nucleotide polymorphisms within nuclear genes that encode the electron transport chain

We observed significant associations between birthweight and SNPs within gene regions encoding for structural and functional components of the ETC across complex I, complex II, complex IV, and ATP synthase. The largest number of SNPs was present within gene regions encoding complex I (NADH: Ubiquinone oxidoreductase - NDUF). The assembly of complex I is an intricate multistep process that varies between species and is extensively reviewed elsewhere (26–29). We identified SNPs within regions that encode genes responsible for mitochondrial complex I assembly, including NDUFA6, an accessory subunit required for complex I assembly, and NDUFA10, an accessory subunit of complex I that likely plays a role in assembly (29). In addition to identifying SNPs associated with assembly, our GWAS analysis identified several SNPs within regions that encode functional components of complex I. Notably, NDUFA5 which, while required for the formation and stability of the extramembrane arm of complex I, is also thought to primarily aid the transfer of electrons through complex I, with a depleted cell line showing up to 80% reduction in complex I activity (30). Further, we identified two SNPs within gene regions that encode two of the seven crucial iron-sulfur proteins, NDUFS6 and NDUFS3, essential for the transfer of electrons through the complex, the latter encoding the seventh and last core iron-sulfur complex within complex I (31).

Our analyses also identified associations between birthweight and SNPs within SDHC, which encodes one of two proteins that anchor complex II (succinate dehydrogenase -SDH) to the inner mitochondrial membrane and form the catalytic core of succinate dehydrogenase. SNPs within complex IV (cytochrome c oxidase - COX) were found to be associated with low birthweight, including COX5A, a subunit of complex IV that, in conjunction with the COX4 subunit, forms a heterodimer that binds with the COX1 subunit, an essential process for complex IV biogenesis. Similarly, we identified a SNP within the region encoding COX7A2L, also known as supercomplex assembly factor 1 (SCAF1), a subunit essential for respiratory supercomplex formation (the association of complex I, III, and IV) (32, 33). In addition, our data identified a SNP in a gene region coding for the ATP synthase subunit ATP5MD. ATP5MD (known by the aliases ATP5MK, Diabetes-associated protein in insulin-sensitive tissues (DAPIT), and Up-regulated during skeletal muscle growth protein 5 (USMG5)) is a membrane subunit that extends into the intermembrane space, required for the dimerization of ATP synthase, a process essential for membrane curvature and cristae formation (34, 35). Collectively these findings suggest that SNPs that affect mitochondrial assembly and function are associated with low birthweight outcomes. Our data highlights the role of nuclear genetics in mitochondrial structure and function. Excitingly, this analysis also identified that SNPs can be observed in the maternal genome, the fetal genome (and thereby contributed by the paternal genome), and in the case of rs34589411 the SNP is found to be associated with birthweight in both the maternal and fetal genomes (Table 3). These data suggest that there is an inheritable contribution from maternal and paternal nuclear genes that affect placental mitochondrial function, that is associated with impaired fetal growth.

### The role of SNPs in the pathology of fetal growth restriction

While our GWAS analysis allowed the discovery of SNPs that lie within the maternal or fetal nuclear genomes encoding proteins that affect mitochondrial structure and function are associated with birthweight is novel, in isolation it does not demonstrate that these drive pathology. To investigate potential protein consequences of these SNPs we collected placentas from a cohort of pregnancies affected by fetal growth restriction (Table 4). Clinically our FGR samples were diagnosed in-utero in line with the Delphi consensus and estimated birthweight (3). Fetal biometry of our FGR cohort showed a 12% reduction in abdominal circumference and femur length and an 11% reduction in head circumference in our FGR compared to health pregnancies (Table 4). Diagnosis of FGR was confirmed upon birth and categorised by birthweight centiles, with our FGR cohort born on average 1,453g smaller than healthy term controls. In line with the general consensus that FGR results from placental insufficiency and poor placental development our FGR placentas were 236g (33%) smaller than the placentas from the healthy control group.

Of the 10 SNPs identified in our GWAS and assessed in FGR placentas, six were confirmed by Sanger sequencing (Table 5). Our analysis determined that three of our SNPS reside within genes that encode complex I of the electron transport chain NDUFA5, NDUFS3, and NDUFS6. Further, two of these SNPs were present at a disproportionately greater percentage compared to their expected population prevalence on gnomAD. With NDUFS6 - rs3822362 present at only 7% in the population, we identified a G>A in 20% of our FGR cohort. While NDUFA5 - rs6737 is expected at 23% our FGR samples identified a G>C SNP in 10% and a C>G change in 60%, collectively observing an allele shift in 70% of our FGR samples. NDUFS3 did not exceed population prevalence and was identified in 20% of our FGR samples, in line with the expected prevalence. These findings identify SNPs within nuclear genes that encode mitochondrial components found within the placenta. Further these SNPs are present at significantly greater proportion than expected in placentas associated with FGR. Moreover, our results identified an additional SNP (n=1) not predicted within our GWAS analysis within the gene region encoding COX7A2L, and two deletions within NDUFA6 and NDUFS6. Upon subsequent analysis of our SNPs, mutation taster predicted “polymorphism” or “probably harmless”, but these do not fall into the “polymorphism automatic” prediction which are “known to be harmless” (Table 5).

Our findings suggest that SNPs within the nuclear genome may be responsible for mitochondrial dysfunction in FGR and our identified SNPs are potentially deleterious. Indeed, this is not the first study to associate nuclear encoded SNPs with the aetiology of disease, others have shown that mutations within NDUFS6 resulting from splice variants lead to complex I deficiency, and are linked to complex I deficiency-specific cardiomyopathy, and lethal neonatal mitochondrial complex I deficiency (36, 37). Further, NDUFS6 mutation is known to contribute to the primary mitochondrial disease Leigh syndrome (38), and has been shown to be hypomethylated in cord blood from growth restricted fetuses when compared to those that are appropriate for gestational age (39). In addition, mutations in NDUFS3, another iron-sulfur cluster coding gene, has been shown by denaturing high-performance liquid chromatography and RT-PCR to induce the delayed onset of mitochondrial complex I deficiency, optic atrophy and Leigh syndrome due to amino acid alterations (31). Moreover, mutations within NDUFA10, although identified in our GWAS analysis, but not confirmed in our FGR cohort, has been identified in association with mitochondrial deficiencies in Leigh syndrome (40).

In addition to complex I, we identified an association between SNPs and birthweight (Table 3) in several complex IV gene regions with SNPs present in the gene regions of COX5A, COX7A2L, and COX8A, with the latter predicted to result in splice changes and protein feature alterations (Table 5). Like complex I, mutations within complex IV have been known to drive pathologies, with splice mutations in COX8A shown to underpin primary mitochondrial disease and to present as a Leigh-like syndrome (41). The majority of studies examining the role of COX7A2L have been performed in mice, with pathological variants yet to be definitively identified in humans. Our study identified an association between COX7A2L - rs10206058 and birthweight in our GWAS analysis, and confirmed its presence within a FGR placenta. Although absent from our GWAS analysis we subsequently identified the presence of an additional SNP COX7A2L - 42356527 in FGR. These data suggest there may be a role of COX7A2L in human diseases, specifically FGR.

Given the similarities between our SNPs and those known to induce mitochondrial disease, Leigh-syndrome or Leigh-like syndrome, we suggest that like these pathologies, FGR has an inheritable component and is driven by mitochondrial dysfunction. These findings are supported by the identification of SNPs in nuclear genes associated with birthweight (Table 3).

### Programmed mitochondrial complex I dysfunction in fetal growth restriction

Given that 50% of the SNPs confirmed within our FGR cohort were present in complex I we examined the expression of genes and corresponding protein levels of NDUFA5, NDUFA6, NDUFA10, NDUFS3, and NDUFS6 (Figure 3). We identified dysregulation of complex I assembly components in FGR placentas and an altered structural ability to transfer electrons to ubiquinone. Specifically, we observed an increase in NDUFA5 at both the gene and protein level in FGR. NDUSFA5 is a supernumerary subunit, located within the inner mitochondrial membrane required for the formation of the Q module in the extramembrane arm of complex I, in addition to facilitating the transfer of electrons to ubiquinone (30). Our findings showed no change in accessory subunit NDUFA10, and a conserved decrease in gene and protein of supernumerary subunit NDUFA6 in FGR compared to controls. Similarly, we identified that two of the seven crucial iron-sulfur proteins, NDUFS3 and NDUFS6, were downregulated at the protein level, with NDUFS6 the most severely dysregulated in FGR. This was somewhat expected given that our SNPs were identified within NDUFS6 in 20% of FGR samples and a deletion was present in 100% of our samples, with predicted splice changes corresponding with G>A rs3822362 (Table 5). Subsequent validation by western blot confirmed our proteomics data and showed significantly decreased NDUFS6 protein level in FGR placentas when compared to healthy controls.

As structure is so crucial to the functional capacity of mitochondria, we can surmise that the decline in assembly capacity due to altered protein levels observed in NDUFA6, in combination with impaired electron flux resulting from a decline in NDUFS3 and supernumerary subunit NDUFS6 would profoundly impact bioenergetics, trophoblast homeostasis, and influence the ability of the FGR placenta to develop, and thereby support fetal growth. This concept is supported by our observations in a subset of FGR placentas (Figure 3I), where maximum ATP capacity was reduced in comparison to controls. The functional consequences of reduced iron-sulfur proteins NDUFS3 and NDUFS6 may include an increase in electron leak from complex I due to an inability of electrons to move efficiently across complex I and an insufficiency in the terminal exchange with ubiquinone. These data provide an explanation for the well-documented increase in reactive oxygen species observed in FGR placentas (42). In addition to electron transfer, NDUFS6 is responsible for the assembly and stabilisation of complex I, where, in association with NDUFS4, NDUFA12, and NDUFAF2, it forms the machinery for the attachment of the N module in complex I biogenesis (43, 44). Our observations of Q module biogenesis are less clear, given the contrasting effects of NDUFA5 increasing, and NDUFS3 decreasing in FGR. The implications of these changes are difficult to discern given the current understanding within the literature on the accessory subunit NDUFA5. Although the role of the core subunit NDUFS3 in early Q-module synthesis is presumed to precede NDUFA5, studies have shown that depletion of the core unit NDUFS3 still facilitates a small amount of complex I assembly, albeit in supercomplexes, with Q module reductions preceding N-module decline (45). These findings may explain our observed decline in N module proteins in FGR, while concurrently detecting NDUFA5 at a greater rate than in control placentas. A decline in mitochondrial assembly may be supported by observations of decreased mitochondrial content in FGR within the literature (5, 46, 47).

Given the complex nature of placental formation and the central role cytotrophoblast cells play in the development and sustained growth of the placenta through fusing to form syncytiotrophoblast we examined the expression level of NDUFS6 within cytotrophoblast (Cyto-Mito) and syncytiotrophoblast mitochondria (Syncytio-Mito) (Figure 3) in FGR. Examination of NDUFS6 from isolated mitochondria from both of these cell types showed a significant decrease in FGR. Collectively these findings may provide a rationale for poor cytotrophoblast function and proliferation rates, and the reduced number of cytotrophoblast and terminally differentiated syncytiotrophoblast observed in FGR (4). The resulting underdeveloped and reduced number of villi, limit the surface area for placental exchange in FGR (37). Our data suggests these placental developmental changes are a result of SNPs programming mitochondrial dysfunction, and thus trophoblast dysfunction in FGR. This interpretation is supported by our clinical findings of significantly smaller placentas in FGR (Table 4).

In combination these data provide evidence that SNPs within nuclear genes that encode mitochondrial structure and function result in programmed dysregulation of complex I formation in fetal growth restriction. We propose these changes underpin poor functioning trophoblasts, resulting in placental insufficiency, reduced placental weight, and the reduced fetal weight observed in FGR.

### Limitations

To our knowledge, this study is the first to investigate targeted mitochondrial pathways regulated by the nuclear genome in association with birthweight. While significant SNPs were identified using the EGG consortium, no information on pathology of these samples or rates of FGR pregnancies was available. This led to the collection and analysis of FGR samples in a new cohort. We employed strict inclusion-exclusion criteria for healthy control and FGR samples to ensure that to the best of our ability we assessed the pathology of FGR and not comorbidities or lifestyle factors influencing our observations. Exclusion criteria are expanded upon in the methods. It should be noted that while SNPs were observed and confirmed in FGR, we believe that for mitochondrial dysfunction to result in pathology an effect threshold must be reached in which the mutated mitochondria are the dominant population. Ordinarily, this would correspond to the severity of primary mitochondrial disease and is dependent on heteroplasmy, referring to two or more populations of mtDNA present within a cell (48). This effect has been shown to be dependent on tissue type, with a lower disease threshold required in more oxidative tissues due to their higher vulnerability to mutations, and more frequent replication of mitochondria (49, 50). The list of recognised influences on heteroplasmy has recently expanded beyond mtDNA, to include a heritable nuclear component (51). Our findings align with these data, identifying SNPs encoded within the nuclear genome, and therefore expected to exert their effect on all mitochondria within the tissue. While heteroplasmy may not be involved, we suggest that a threshold effect is still present, whereby, if functional consequences exceed a viability threshold pregnancy would not progress due to a failure of trophoblast cells and the placenta. This may account for the variability in pathology, thus, while we have identified significant mitochondrial dysfunction predominantly in complex I in FGR corresponding with our identified SNPs, to confirm clinical causation of any single SNP requires a validation cohort. Given the nature of this project to assess and confirm SNPs in the nuclear genome of FGR, proteomics and western blotting was performed on whole tissue homogenates comprising of various cell lineages to best determine the presence and effect of SNPs. Although complicated by the complex and unique subpopulations of mitochondria present in the placenta, addressed in Figure 3, future studies should build upon our findings and consider the contribution of SNPs programing mitochondrial dysfunction in specific cell types within the same tissue.

## Conclusion

Our findings provide a novel insight into the role of mitochondria in pregnancy, concluding 37 SNPs were significantly associated with birthweight. All SNPs were found within the nuclear genome that encode mitochondrial structure or function, and to our knowledge have not previously been identified in other large GWAS meta-analyses examining the regulation of birthweight and adult disease (Figure 1) (24). Of the genes examined in a refined analysis of two mitochondrial pathways encoding the electron transport chain and ATP synthase (289 genes), 12.8% contained SNPs associated with birthweight, residing in gene regions which encode subunits of the electron transport chain, NDUFA5, NDUFA6, NDUFA10, NDUFS3, NDUFS6, SDHC, COX5A, COX7A2L, COX8A and ATP synthase, ATP5MD. For the first time we identified SNPs found to be unique to maternal and fetal summary data, enabling us to ascertain a paternal contribution to fetal birthweight. This was possible as the 17 SNPs present in gene regions unique to the fetus and absent from the maternal genome (Table 3) can only be attributed to the paternal genome. Thus, we provide evidence that mitochondrial inheritance and the maternal genome, while important, and the predominant influence behind known pathologies, is not the sole contributor to mitochondrial insufficiency. Our confirmation of the presence of these SNPs within placentas from a primary dataset of fetal growth restriction demonstrates the role of SNPs in the pathology. Further, our investigation determined that the SNPs present in FGR had the predicted potential to alter gene splicing or protein function. Our subsequent investigation identified that these SNPs corresponded to gene expression and protein changes in complex I. Specifically, these changes were observed in proteins that permit complex I assembly and the transfer of electrons to ubiquinone, thereby impeding electron flux and ATP generating capacity. In combination, our findings suggest that it is highly likely that a mismatch or incompatibility in nuclear maternal-paternal genomes dysregulate complex I assembly and structure, programming mitochondrial dysfunction in FGR. We suggest that these arise in response to SNPs in a manner akin to other primary mitochondrial diseases. Further, we suggest that this programmed dysfunction occurs predominantly within the cytotrophoblast cell lineage of FGR, underpinning poor placental growth, resulting in placental insufficiency and ultimately a decreased placental weight and fetal birthweight. Our study demonstrates the complex nature of mitochondrial function and establishes the premise for genetic complementarity in gestational outcomes identifying the need for further investigation into the physiological and pathophysiological role of SNPs in gestational and placental pathologies.

## Acknowledgements

Data on birthweight trait has been contributed by the EGG Consortium using the UK Biobank Resource and has been downloaded from www.egg-consortium.org. The Australian Genome Research Facility (AGRF) https://www.agrf.org.au/ was utilized for the design and validation of primers used in Sanger Sequencing, sample processing and data generation. This research was supported by National Health & Medical Research Council Investigator Grant GNT2026065.

