## Supplementary Data for "The genetic origin of fetal growth restriction and mitochondrial complex I dysregulation"

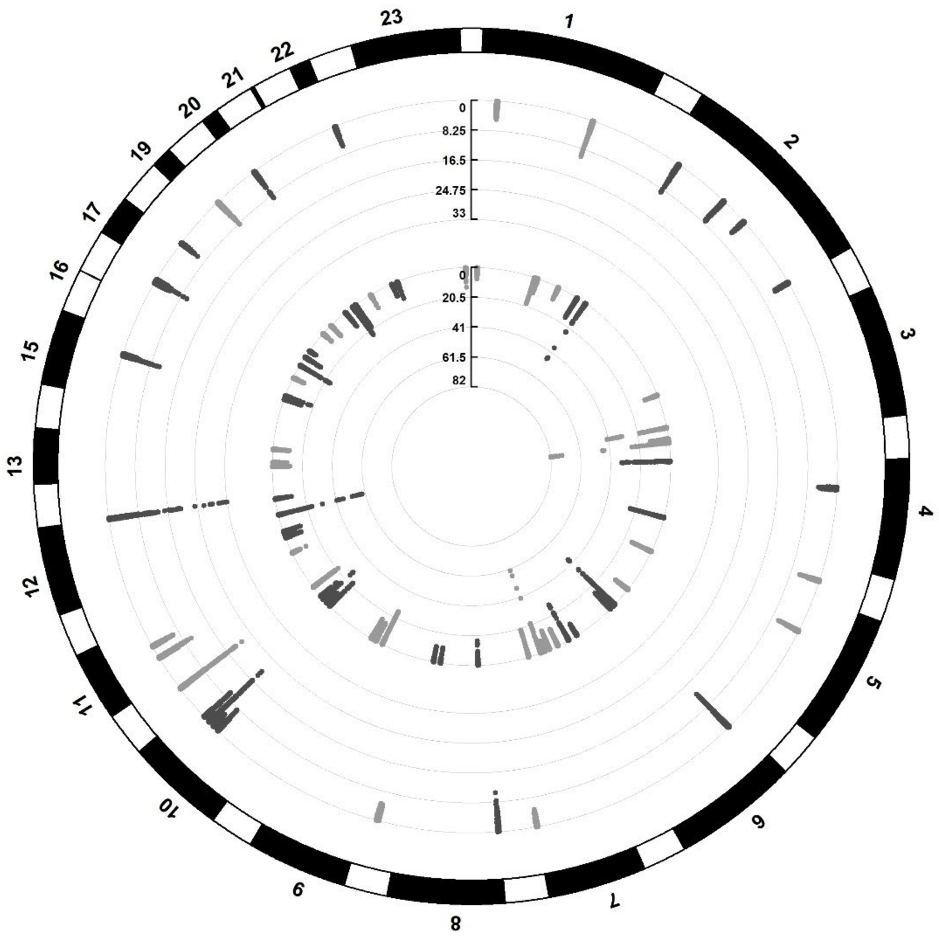


Supplementary Figure 1: Shows a circular Manhattan plot with mitochondrial gene regions that are significantly associated with birthweight; the inner ring contains SNPs previously associated with decreased birthweight. The outer ring shows significant regions unique to mitochondrial gene regions regulating the ETC and ATP synthase. Numbers around the outside represent the chromosome. GWAS significance is established at (p <5 x10-8).

| Gene Set | Brief description | N | n | P-value | adj. P-value | Genes |
|---|---|---|---|---|---|---|
| ELECTRON TRANSPORT CHAIN | A process in which a series of electron carriers operate together to transfer electrons from donors to any of several different terminal electron acceptors. | 173 | 12 | 1.79E-20 | 1.00E-16 | SDHC:NDUFS3:COX8A:ALDH2:CYP1A2:MYBBP1A:POLG2:COX7A2L:NDUFA6:CCNB1:NDUFA5:GLDC |
| GENERATION OF PRECURSOR METABOLITES AND ENERGY | The chemical reactions and pathways resulting in the formation of precursor metabolites, substances from which energy is derived, and any process involved in the liberation of energy from these substances | 488 | 15 | 2.73E-20 | 1.00E-16 | SDHC:IGF2:INS:NDUFS3:COX8A:ALDH2:CYP1A2:MYBBP1A:POLG2:GSK3A:COX7A2L:NDUFA6:CCNB1:NDUFA5:GLDC |
| ENERGY DERIVATION BY OXIDATION OF ORGANIC COMPOUNDS | The chemical reactions and pathways by which a cell derives energy from organic compounds; results in the oxidation of the compounds from which energy is released | 269 | 13 | 4.22E-20 | 1.03E-16 | SDHC:IGF2:INS:NDUFS3:COX8A:CYP1A2:MYBBP1A:POLG2:GSK3A:COX7A2L:NDUFA6:CCNB1:NDUFA5 |
| RESPIRATORY ELECTRON TRANSPORT CHAIN | A process in which a series of electron carriers operate together to transfer electrons from donors such as NADH and FADH2 to any of several different terminal electron acceptors to generate a transmembrane electrochemical gradient. | 103 | 9 | 2.13E-16 | 3.92E-13 | SDHC:NDUFS3:COX8A:MYBBP1A:POLG2:COX7A2L:NDUFA6:CCNB1:NDUFA5 |
| CELLULAR RESPIRATION | The enzymatic release of energy from inorganic and organic compounds (especially carbohydrates and fats) which either requires oxygen (aerobic respiration) or does not (anaerobic respiration) | 179 | 10 | 3.69E-16 | 5.42E-13 | SDHC:NDUFS3:COX8A:CYP1A2:MYBBP1A:POLG2:COX7A2L:NDUFA6:CCNB1:NDUFA5 |
| OXIDATION REDUCTION PROCESS | A metabolic process that results in the removal or addition of one or more electrons to or from a substance, with or without the concomitant removal or addition of a proton or protons. | 954 | 15 | 5.77E-16 | 7.06E-13 | SDHC:IGF2:INS:NDUFS3:COX8A:ALDH2:CYP1A2:MYBBP1A:POLG2:GSK3A:COX7A2L:NDUFA6:CCNB1:NDUFA5:GLDC |
| ATP SYNTHESIS COUPLED ELECTRON TRANSPORT | The transfer of electrons through a series of electron donors and acceptors, generating energy that is ultimately used for synthesis of ATP. | 84 | 7 | 8.67E-13 | 9.10E-10 | SDHC:NDUFS3:COX8A:COX7A2L:NDUFA6:CCNB1:NDUFA5 |

| | | | | | | |
|---|---|---|---|---|---|---|
| OXIDATIVE PHOSPHORYLATION | The phosphorylation of ADP to ATP that accompanies the oxidation of a metabolite through the operation of the respiratory chain. Oxidation of compounds establishes a proton gradient across the membrane, providing the energy for ATP synthesis. | 126 | 7 | 1.58E-11 | 1.45E-08 | SDHC:NDUFS3:COX8A:COX7A2L:NDUFA6:CCNB1:NDUFA5 |
| RESPONSE TO OXYGEN CONTAINING COMPOUND | Any process that results in a change in state or activity of a cell or an organism (in terms of movement, secretion, enzyme production, gene expression, etc.) as a result of an oxygen-containing compound stimulus. | 1603 | 12 | 4.90E-09 | 4.00E-06 | IGF2:INS:CYP1A2:GSK3A:IL1B:GSS:PPAT:CCNB1:ATP6V1G2:NFKBIL1:JAK2:GLDC |
| CELLULAR RESPONSE TO PEPTIDE HORMONE STIMULUS | Any process that results in a change in state or activity of a cell (in terms of movement, secretion, enzyme production, gene expression, etc.) as a result of a peptide hormone stimulus. A peptide hormone is any of a class of peptides that are secreted into the blood stream and have endocrine functions in living animals. | 315 | 7 | 9.51E-09 | 6.99E-06 | IGF2:INS:GSK3A:IL1B:PPAT:ATP6V1G2:JAK2 |
| DE NOVO IMP BIOSYNTHETIC PROCESS | The chemical reactions and pathways resulting in the formation of IMP, inosine monophosphate, by the stepwise assembly of a purine ring on ribose 5-phosphate. | 6 | 3 | 1.24E-08 | 8.29E-06 | PFAS:PPAT:PAICS |
| CELLULAR RESPONSE TO INSULIN STIMULUS | Any process that results in a change in state or activity of a cell (in terms of movement, secretion, enzyme production, gene expression, etc.) as a result of an insulin stimulus. Insulin is a polypeptide hormone produced by the islets of Langerhans of the pancreas in mammals, and by the homologous organs of other organisms | 213 | 6 | 3.00E-08 | 1.84E-05 | IGF2:INS:GSK3A:IL1B:PPAT:ATP6V1G2 |
| CELLULAR RESPONSE TO PEPTIDE | Any process that results in a change in state or activity of a cell (in terms of movement, secretion, enzyme production, gene | 377 | 7 | 3.26E-08 | 1.84E-05 | IGF2:INS:GSK3A:IL1B:PPAT:ATP6V1G2:JAK2 |

| | | | | | | |
|---|---|---|---|---|---|---|
| | expression, etc.) as a result of a peptide stimulus. | | | | | |
| RESPONSE TO PEPTIDE HORMONE | Any process that results in a change in state or activity of a cell or an organism (in terms of movement, secretion, enzyme production, gene expression, etc.) as a result of a peptide hormone stimulus. A peptide hormone is any of a class of peptides that are secreted into the blood stream and have endocrine functions in living animals. | 425 | 7 | 7.37E-08 | 3.64E-05 | IGF2:INS:GSK3A:IL1B:PPAT:ATP6V1G2:JAK2 |
| IMP BIOSYNTHETIC PROCESS | The chemical reactions and pathways resulting in the formation of IMP, inosine monophosphate. | 10 | 3 | 7.43E-08 | 3.64E-05 | PFAS:PPAT:PAICS |
| RESPONSE TO NITROGEN COMPOUND | Any process that results in a change in state or activity of a cell or an organism (in terms of movement, secretion, enzyme production, gene expression, etc.) as a result of a nitrogen compound stimulus. | 968 | 9 | 1.03E-07 | 4.49E-05 | IGF2:INS:GSK3A:IL1B:GSS:PPAT:ATP6V1G2:JAK2:GLDC |
| SMALL MOLECULE METABOLIC PROCESS | The chemical reactions and pathways involving small molecules, any low molecular weight, monomeric, non-encoded molecule. | 1685 | 11 | 1.04E-07 | 4.49E-05 | CTPS1:IGF2:INS:ALDH2:CYP1A2:PFAS:GSK3A:IL1B:GSS:PPAT:GLDC |
| RESPONSE TO INSULIN | Any process that results in a change in state or activity of a cell or an organism (in terms of movement, secretion, enzyme production, gene expression, etc.) as a result of an insulin stimulus. Insulin is a polypeptide hormone produced by the islets of Langerhans of the pancreas in mammals, and by the homologous organs of other organisms. | 269 | 6 | 1.19E-07 | 4.87E-05 | IGF2:INS:GSK3A:IL1B:PPAT:ATP6V1G2 |
| INSULIN RECEPTOR SIGNALING PATHWAY | The series of molecular signals generated as a consequence of the insulin receptor binding to insulin. | 139 | 5 | 1.41E-07 | 5.45E-05 | IGF2:INS:GSK3A:IL1B:ATP6V1G2 |
| REGULATION OF GENERATION OF | Any process that modulates the frequency, rate or extent of the chemical reactions and | 145 | 5 | 1.74E-07 | 6.39E-05 | IGF2:INS:GSK3A:COX7A2L:CCNB1 |

| | | | | | | |
|---|---|---|---|---|---|---|
| PRECURSOR METABOLITES AND ENERGY | pathways resulting in the formation of precursor metabolites, substances from which energy is derived, and the processes involved in the liberation of energy from these substances. | | | | | |
| CELLULAR AMINO ACID METABOLIC PROCESS | The chemical reactions and pathways involving amino acids, carboxylic acids containing one or more amino groups, as carried out by individual cells. | 290 | 6 | 1.86E-07 | 6.50E-05 | CTPS1:INS:PFAS:GSS:PPAT:GLDC |
| RESPONSE TO PEPTIDE | Any process that results in a change in state or activity of a cell or an organism (in terms of movement, secretion, enzyme production, gene expression, etc.) as a result of a peptide stimulus. | 503 | 7 | 2.31E-07 | 7.71E-05 | IGF2:INS:GSK3A:IL1B:PPAT:ATP6V1G2:JAK2 |
| MITOCHONDRION ORGANIZATION | A process that is carried out at the cellular level which results in the assembly, arrangement of constituent parts, or disassembly of a mitochondrion; includes mitochondrial morphogenesis and distribution, and replication of the mitochondrial genome as well as synthesis of new mitochondrial components. | 512 | 7 | 2.60E-07 | 8.31E-05 | NDUFS3:POLG2:GSK3A:COX7A2L:NDUFA6:KDR:NDUFA5 |
| CELLULAR RESPONSE TO OXYGEN CONTAINING COMPOUND | Any process that results in a change in state or activity of a cell (in terms of movement, secretion, enzyme production, gene expression, etc.) as a result of an oxygen-containing compound stimulus. | 1118 | 9 | 3.47E-07 | 0.000106 | IGF2:INS:GSK3A:IL1B:PPAT:CCNB1:ATP6V1G2:NFKBIL1:JAK2 |
| REGULATION OF CELLULAR RESPONSE TO INSULIN STIMULUS | Any process that modulates the frequency, rate or extent of cellular response to insulin stimulus. | 70 | 4 | 4.36E-07 | 0.000128 | IGF2:INS:GSK3A:IL1B |
| RIBOSE PHOSPHATE BIOSYNTHETIC PROCESS | The chemical reactions and pathways resulting in the formation of ribose phosphate, any phosphorylated ribose sugar. | 183 | 5 | 5.52E-07 | 0.000156 | CTPS1:PAPSS2:PFAS:PPAT:PAICS |

| | | | | | | |
|---|---|---|---|---|---|---|
| NUCLEOBASE BIOSYNTHETIC PROCESS | The chemical reactions and pathways resulting in the formation of a nucleobase, a nitrogenous base that is a constituent of a nucleic acid. | 19 | 3 | 5.97E-07 | 0.000162 | CTPS1:PPAT:PAICS |
| PURINE NUCLEOSIDE MONOPHOSPHATE BIOSYNTHETIC PROCESS | The chemical reactions and pathways resulting in the formation of purine nucleoside monophosphate, a compound consisting of a purine base linked to a ribose or deoxyribose sugar esterified with phosphate on the sugar. [GOC:go_curators, | 20 | 3 | 7.01E-07 | 0.000184 | PFAS:PPAT:PAICS |
| CELLULAR RESPONSE TO NITROGEN COMPOUND | Any process that results in a change in state or activity of a cell (in terms of movement, secretion, enzyme production, gene expression, etc.) as a result of a nitrogen compound stimulus. | 632 | 7 | 1.06E-06 | 0.00027 | IGF2:INS:GSK3A:IL1B:PPAT:ATP6V1G2:JAK2 |
| POSITIVE REGULATION OF CELL CYCLE | Any process that activates or increases the rate or extent of progression through the cell cycle. | 402 | 6 | 1.25E-06 | 0.000306 | IGF2:INS:MYBBP1A:IL1B:CCNB1:DDX39B |
| RESPONSE TO HORMONE | Any process that results in a change in state or activity of a cell or an organism (in terms of movement, secretion, enzyme production, gene expression, etc.) as a result of a hormone stimulus. | 973 | 8 | 1.50E-06 | 0.000356 | IGF2:INS:CYP1A2:GSK3A:IL1B:PPAT:ATP6V1G2:JAK2 |
| CELLULAR RESPONSE TO HORMONE STIMULUS | Any process that results in a change in state or activity of a cell (in terms of movement, secretion, enzyme production, gene expression, etc.) as a result of a hormone stimulus. | 689 | 7 | 1.89E-06 | 0.000433 | IGF2:INS:GSK3A:IL1B:PPAT:ATP6V1G2:JAK2 |
| TRANSMEMBRANE RECEPTOR PROTEIN TYROSINE KINASE SIGNALING PATHWAY | Any process that initiates the activity of the inactive transmembrane receptor protein tyrosine kinase activity. | 712 | 7 | 2.34E-06 | 0.000511 | IGF2:INS:GSK3A:IL1B:KDR:ATP6V1G2:JAK2 |
| ENZYME LINKED RECEPTOR PROTEIN | Any series of molecular signals initiated by the binding of an extracellular ligand to a | 1034 | 8 | 2.36E-06 | 0.000511 | IGF2:INS:GSK3A:IL1B:ACVR2A:KDR:ATP6V1G2:JAK2 |

| | | | | | | |
|---|---|---|---|---|---|---|
| SIGNALING PATHWAY | receptor on the surface of the target cell, where the receptor possesses catalytic activity or is closely associated with an enzyme such as a protein kinase, and ending with regulation of a downstream cellular process, e.g. transcription. | | | | | |
| REACTIVE OXYGEN SPECIES BIOSYNTHETIC PROCESS | The chemical reactions and pathways resulting in the formation of reactive oxygen species, any molecules or ions formed by the incomplete one-electron reduction of oxygen | 112 | 4 | 2.88E-06 | 0.000604 | INS:CYP1A2:IL1B:JAK2 |
| RIBONUCLEOSIDE MONOPHOSPHATE BIOSYNTHETIC PROCESS | The chemical reactions and pathways resulting in the formation of a ribonucleoside monophosphate, a compound consisting of a nucleobase linked to a ribose sugar esterified with phosphate on the sugar. | 33 | 3 | 3.33E-06 | 0.00068 | PFAS:PPAT:PAICS |
| REGULATION OF GLYCOGEN METABOLIC PROCESS | Any process that modulates the frequency, rate or extent of the chemical reactions and pathways involving glycogen. | 34 | 3 | 3.65E-06 | 0.000725 | IGF2:INS:GSK3A |
| NUCLEOSIDE PHOSPHATE BIOSYNTHETIC PROCESS | The chemical reactions and pathways resulting in the formation of a nucleoside phosphate. | 273 | 5 | 3.93E-06 | 0.00076 | CTPS1:PAPSS2:PFAS:PPAT:PAICS |
| HISTONE PHOSPHORYLATION | The modification of histones by addition of phosphate groups | 38 | 3 | 5.13E-06 | 0.000968 | IL1B:CCNB1:JAK2 |
| POSITIVE REGULATION OF CELL CYCLE PROCESS | Any process that increases the rate, frequency or extent of a cellular process that is involved in the progression of biochemical and morphological phases and events that occur in a cell during successive cell replication or nuclear replication events. | 296 | 5 | 5.82E-06 | 0.00107 | IGF2:INS:MYBBP1A:IL1B:CCNB1 |
| REGULATION OF POLYSACCHARIDE METABOLIC PROCESS | Any process that modulates the frequency, rate or extent of the chemical reactions and pathways involving polysaccharides. | 42 | 3 | 6.97E-06 | 0.00125 | IGF2:INS:GSK3A |

| | | | | | | |
|---|---|---|---|---|---|---|
| POSITIVE REGULATION OF PROTEIN METABOLIC PROCESS | Any process that activates or increases the frequency, rate or extent of the chemical reactions and pathways involving a protein. | 1615 | 9 | 7.19E-06 | 0.001259 | IGF2:INS:GSK3A:IL1B:ACVR2A:KDR:CCNB1:DDX39B:JAK2 |
| POSITIVE REGULATION OF GLYCOGEN STARCH SYNTHASE ACTIVITY | Any process that activates or increases the frequency, rate or extent of glycogen (starch) synthase activity. | 5 | 2 | 7.52E-06 | 0.00127 | IGF2:GSK3A |
| POSITIVE REGULATION OF PROTEIN MODIFICATION PROCESS | Any process that activates or increases the frequency, rate or extent of the covalent alteration of one or more amino acid residues within a protein | 1212 | 8 | 7.61E-06 | 0.00127 | IGF2:INS:GSK3A:IL1B:ACVR2A:KDR:CCNB1:JAK2 |
| RESPONSE TO ENDOGENOUS STIMULUS | Any process that results in a change in state or activity of a cell or an organism (in terms of movement, secretion, enzyme production, gene expression, etc.) as a result of a stimulus arising within the organism. | 1634 | 9 | 7.91E-06 | 0.001283 | IGF2:INS:CYP1A2:GSK3A:IL1B:ACVR2A:PPAT:ATP6V1G2:JAK2 |
| NUCLEOSIDE MONOPHOSPHATE BIOSYNTHETIC PROCESS | The chemical reactions and pathways resulting in the formation of a nucleoside monophosphate, a compound consisting of a nucleobase linked to a deoxyribose or ribose sugar esterified with phosphate on the sugar. | 44 | 3 | 8.03E-06 | 0.001283 | PFAS:PPAT:PAICS |
| GLYCOGEN BIOSYNTHETIC PROCESS | The chemical reactions and pathways resulting in the formation of glycogen, a polydisperse, highly branched glucan composed of chains of D-glucose residues. | 46 | 3 | 9.20E-06 | 0.001408 | IGF2:INS:GSK3A |
| MITOCHONDRIAL ELECTRON TRANSPORT NADH TO UBIQUINONE | The transfer of electrons from NADH to ubiquinone that occurs during oxidative phosphorylation. | 46 | 3 | 9.20E-06 | 0.001408 | NDUFS3:NDUFA6:NDUFA5 |
| ORGANIC ACID METABOLIC PROCESS | The chemical reactions and pathways involving organic acids, any acidic compound containing carbon in covalent linkage. | 901 | 7 | 1.09E-05 | 0.00164 | CTPS1:INS:CYP1A2:PFAS:GSS:PPAT:GLDC |

| | | | | | | |
|---|---|---|---|---|---|---|
| APOPTOTIC SIGNALING PATHWAY | A series of molecular signals which triggers the apoptotic death of a cell. The pathway starts with reception of a signal, and ends when the execution phase of apoptosis is triggered. | 591 | 6 | 1.14E-05 | 0.00167 | INS:NDUFS3:MYBBP1A:GSK3A:IL1B:JAK2 |
| REGULATION OF NITRIC OXIDE SYNTHASE ACTIVITY | Any process that modulates the activity of the enzyme nitric-oxide synthase. | 51 | 3 | 1.26E-05 | 0.001813 | INS:IL1B:ACVR2A |
| POSITIVE REGULATION OF MITOTIC CELL CYCLE | Any process that activates or increases the rate or extent of progression through the mitotic cell cycle. | 166 | 4 | 1.37E-05 | 0.001933 | IGF2:INS:IL1B:CCNB1 |
| ALPHA AMINO ACID METABOLIC PROCESS | The chemical reactions and pathways involving an alpha-amino acid | 167 | 4 | 1.40E-05 | 0.001942 | CTPS1:PFAS:PPAT:GLDC |
| POSITIVE REGULATION OF ORGANELLE ORGANIZATION | Any process that increases the frequency, rate or extent of a process involved in the formation, arrangement of constituent parts, or disassembly of an organelle. | 619 | 6 | 1.48E-05 | 0.001981 | IGF2:INS:GSK3A:IL1B:KDR:CCNB1 |
| POSITIVE REGULATION OF MITOTIC NUCLEAR DIVISION | Any process that activates or increases the frequency, rate or extent of mitosis. | 54 | 3 | 1.50E-05 | 0.001981 | IGF2:INS:IL1B |
| POSITIVE REGULATION OF MULTICELLULAR ORGANISMAL PROCESS | Any process that activates or increases the frequency, rate or extent of an organismal process, any of the processes pertinent to the function of an organism above the cellular level; includes the integrated processes of tissues and organs. | 1771 | 9 | 1.51E-05 | 0.001981 | IGF2:INS:GSK3A:IL1B:ACVR2A:KDR:CCNB1:DDX39B:JAK2 |
| NEGATIVE REGULATION OF GLYCOGEN METABOLIC PROCESS | Any process that stops, prevents, or reduces the frequency, rate or extent of the chemical reactions and pathways involving glycogen. | 7 | 2 | 1.58E-05 | 0.002034 | INS:GSK3A |
| POSITIVE REGULATION OF CELL POPULATION PROLIFERATION | Any process that activates or increases the rate or extent of cell proliferation. | 965 | 7 | 1.70E-05 | 0.002157 | IGF2:INS:IL1B:KDR:CCNB1:DDX39B:JAK2 |

| | | | | | | |
|---|---|---|---|---|---|---|
| POSITIVE REGULATION OF TRANSFERASE ACTIVITY | Any process that activates or increases the frequency, rate or extent of transferase activity, the catalysis of the transfer of a group, e.g. a methyl group, glycosyl group, acyl group, phosphorus-containing, or other groups, from a donor compound to an acceptor. | 643 | 6 | 1.83E-05 | 0.002264 | IGF2:INS:GSK3A:IL1B:CCNB1:JAK2 |
| NADH DEHYDROGENASE COMPLEX ASSEMBLY | The aggregation, arrangement and bonding together of a set of components to form an NADH dehydrogenase complex. | 58 | 3 | 1.86E-05 | 0.002264 | NDUFS3:NDUFA6:NDUFA5 |
| CELLULAR RESPONSE TO ENDOGENOUS STIMULUS | Any process that results in a change in state or activity of a cell (in terms of movement, secretion, enzyme production, gene expression, etc.) as a result of a stimulus arising within the organism. | 1373 | 8 | 1.88E-05 | 0.002264 | IGF2:INS:GSK3A:IL1B:ACVR2A:PPAT:ATP6V1G2:JAK2 |
| CARBOHYDRATE DERIVATIVE BIOSYNTHETIC PROCESS | The chemical reactions and pathways resulting in the formation of carbohydrate derivative. | 655 | 6 | 2.03E-05 | 0.002377 | CTPS1:PAPSS2:PFAS:IL1B:PPAT:PAICS |
| POSITIVE REGULATION OF DEVELOPMENTAL PROCESS | Any process that activates or increases the rate or extent of development, the biological process whose specific outcome is the progression of an organism over time from an initial condition (e.g. a zygote, or a young adult) to a later condition (e.g. a multicellular animal or an aged adult). | 1393 | 8 | 2.09E-05 | 0.002377 | IGF2:INS:IL1B:ACVR2A:KDR:CCNB1:DDX39B:JAK2 |
| REGULATION OF GLYCOGEN STARCH SYNTHASE ACTIVITY | Any process that modulates the frequency, rate or extent of glycogen (starch) synthase activity. | 8 | 2 | 2.10E-05 | 0.002377 | IGF2:GSK3A |
| POSITIVE REGULATION OF HISTONE PHOSPHORYLATION | Any process that activates or increases the frequency, rate or extent of the addition of one or more phosphate groups to a histone protein. | 8 | 2 | 2.10E-05 | 0.002377 | IL1B:CCNB1 |
| RESPONSE TO DRUG | Any process that results in a change in state or activity of a cell or an organism (in terms | 1008 | 7 | 2.25E-05 | 0.002509 | CTPS1:CYP1A2:PFAS:IL1B:KDR:PPAT:CCNB1 |

| | | | | | | |
|---|---|---|---|---|---|---|
| | of movement, secretion, enzyme production, gene expression, etc.) as a result of a drug stimulus. A drug is a substance used in the diagnosis, treatment or prevention of a disease. | | | | | |
| REGULATION OF APOPTOTIC SIGNALING PATHWAY | Any process that modulates the frequency, rate or extent of apoptotic signaling pathway. | 396 | 5 | 2.37E-05 | 0.002595 | INS:NDUFS3:GSK3A:IL1B:JAK2 |
| REGULATION OF MONOOXYGENASE ACTIVITY | Any process that modulates the activity of a monooxygenase. | 64 | 3 | 2.50E-05 | 0.002667 | INS:IL1B:ACVR2A |
| REGULATION OF GROWTH | Any process that modulates the frequency, rate or extent of the growth of all or part of an organism so that it occurs at its proper speed, either globally or in a specific part of the organism's development. | 680 | 6 | 2.50E-05 | 0.002667 | IGF2:INS:NDUFS3:GSK3A:CCNB1:DDX39B |
| PROTEIN CONTAINING COMPLEX ASSEMBLY | The aggregation, arrangement and bonding together of a set of macromolecules to form a protein-containing complex. | 1891 | 9 | 2.54E-05 | 0.002668 | INS:NDUFS3:COX7A2L:NDUFA6:PPAT:CCNB1:DDX39B:NDUFA5:JAK2 |
| PURINE CONTAINING COMPOUND BIOSYNTHETIC PROCESS | The chemical reactions and pathways resulting in the formation of a purine-containing compound, i.e. any compound that contains purine or a formal derivative thereof. | 196 | 4 | 2.62E-05 | 0.002717 | PAPSS2:PFAS:PPAT:PAICS |
| POSITIVE REGULATION OF NUCLEAR DIVISION | Any process that activates or increases the frequency, rate or extent of nuclear division, the partitioning of the nucleus and its genetic information. | 66 | 3 | 2.74E-05 | 0.002796 | IGF2:INS:IL1B |
| REGULATION OF NUCLEAR DIVISION | Any process that modulates the frequency, rate or extent of nuclear division, the partitioning of the nucleus and its genetic information. | 206 | 4 | 3.19E-05 | 0.003209 | IGF2:INS:IL1B:CCNB1 |
| REGULATION OF CELLULAR | Any process that modulates the frequency, rate, or extent of the chemical reactions and | 10 | 2 | 3.37E-05 | 0.00333 | INS:GSK3A |

| | | | | | | |
|---|---|---|---|---|---|---|
| CARBOHYDRATE CATABOLIC PROCESS | pathways resulting in the breakdown of carbohydrates, carried out by individual cells. | | | | | |
| POSITIVE REGULATION OF PHOSPHORUS METABOLIC PROCESS | Any process that increases the frequency, rate or extent of the chemical reactions and pathways involving phosphorus or compounds containing phosphorus. | 1075 | 7 | 3.40E-05 | 0.00333 | IGF2:INS:IL1B:ACVR2A:KDR:CCNB1:JAK2 |
| GLUTAMINE FAMILY AMINO ACID METABOLIC PROCESS | The chemical reactions and pathways involving amino acids of the glutamine family, comprising arginine, glutamate, glutamine and proline. | 72 | 3 | 3.56E-05 | 0.00344 | CTPS1:PFAS:PPAT |
| GLUCAN METABOLIC PROCESS | The chemical reactions and pathways involving glucans, polysaccharides consisting only of glucose residues, occurring at the level of an individual cell. | 73 | 3 | 3.71E-05 | 0.003494 | IGF2:INS:GSK3A |
| REGULATION OF GLUCOSE METABOLIC PROCESS | Any process that modulates the rate, frequency or extent of glucose metabolism. Glucose metabolic processes are the chemical reactions and pathways involving glucose, the aldohexose gluco-hexose. | 73 | 3 | 3.71E-05 | 0.003494 | IGF2:INS:GSK3A |
| POLYSACCHARIDE BIOSYNTHETIC PROCESS | The chemical reactions and pathways resulting in the formation of a polysaccharide, a polymer of many (typically more than 10) monosaccharide residues linked glycosidically. | 74 | 3 | 3.86E-05 | 0.003593 | IGF2:INS:GSK3A |
| PURINE NUCLEOBASE BIOSYNTHETIC PROCESS | The chemical reactions and pathways resulting in the formation of purine nucleobases, one of the two classes of nitrogen-containing ring compounds found in DNA and RNA, which include adenine and guanine. | 11 | 2 | 4.12E-05 | 0.003787 | PPAT:PAICS |
| REGULATION OF GLUCOSE TRANSMEMBRANE TRANSPORT | Any process that modulates the frequency, rate or extent of glucose transport across a membrane. Glucose transport is the directed movement of the hexose monosaccharide glucose into, out of or within a cell, or | 79 | 3 | 4.70E-05 | 0.004262 | INS:GSK3A:IL1B |

| | | | | | | |
|---|---|---|---|---|---|---|
| | between cells, by means of some agent such as a transporter or pore. | | | | | |
| ORGANELLE FISSION | The creation of two or more organelles by division of one organelle. | 460 | 5 | 4.83E-05 | 0.004325 | IGF2:INS:IL1B:KDR:CCNB1 |
| REGULATION OF HEART GROWTH | Any process that modulates the rate or extent of heart growth. Heart growth is the increase in size or mass of the heart. | 81 | 3 | 5.06E-05 | 0.004482 | GSK3A:CCNB1:DDX39B |
| AEROBIC RESPIRATION | The enzymatic release of energy from inorganic and organic compounds (especially carbohydrates and fats) which requires oxygen as the terminal electron acceptor. | 82 | 3 | 5.25E-05 | 0.00454 | SDHC:COX8A:COX7A2L |
| CELLULAR CARBOHYDRATE BIOSYNTHETIC PROCESS | The chemical reactions and pathways resulting in the formation of carbohydrates, any of a group of organic compounds based of the general formula Cx(H2O)y, carried out by individual cells. | 82 | 3 | 5.25E-05 | 0.00454 | IGF2:INS:GSK3A |
| POSITIVE REGULATION OF NITRIC OXIDE SYNTHASE BIOSYNTHETIC PROCESS | Any process that activates or increases the frequency, rate or extent of the chemical reactions and pathways resulting in the formation of a nitric oxide synthase enzyme. | 13 | 2 | 5.84E-05 | 0.004933 | KDR:JAK2 |
| REGULATION OF HISTONE PHOSPHORYLATION | Any process that modulates the frequency, rate or extent of the addition of one or more phosphate groups to a histone protein. | 13 | 2 | 5.84E-05 | 0.004933 | IL1B:CCNB1 |
| ENERGY RESERVE METABOLIC PROCESS | The chemical reactions and pathways by which a cell derives energy from stored compounds such as fats or glycogen. | 86 | 3 | 6.05E-05 | 0.004999 | IGF2:INS:GSK3A |
| POSITIVE REGULATION OF PHOSPHATIDYLINOSITOL 3 KINASE SIGNALING | Any process that activates or increases the frequency, rate or extent of signal transduction mediated by the phosphatidylinositol 3-kinase cascade. | 86 | 3 | 6.05E-05 | 0.004999 | INS:KDR:JAK2 |
| CARDIAC CELL DEVELOPMENT | The process whose specific outcome is the progression of a cardiac cell over time, from | 90 | 3 | 6.93E-05 | 0.005599 | GSK3A:CCNB1:DDX39B |

| | | | | | | |
|---|---|---|---|---|---|---|
| | its formation to the mature state. A cardiac cell is a cell that will form part of the cardiac organ of an individual. | | | | | |
| MITOCHONDRIAL RESPIRATORY CHAIN COMPLEX ASSEMBLY | The aggregation, arrangement and bonding together of a set of components to form a mitochondrial respiratory chain complex. | 90 | 3 | 6.93E-05 | 0.005599 | NDUFS3:NDUFA6:NDUFA5 |
| REGULATION OF REACTIVE OXYGEN SPECIES BIOSYNTHETIC PROCESS | Any process that modulates the frequency, rate or extent of reactive oxygen species biosynthetic process. | 92 | 3 | 7.40E-05 | 0.005913 | INS:IL1B:JAK2 |
| REGULATION OF PHOSPHORUS METABOLIC PROCESS | Any process that modulates the frequency, rate or extent of the chemical reactions and pathways involving phosphorus or compounds containing phosphorus. | 1677 | 8 | 7.79E-05 | 0.006157 | IGF2:INS:COX7A2L:IL1B:ACVR2A:KDR:CCNB1:JAK2 |
| PROTEIN KINASE B SIGNALING | A series of reactions, mediated by the intracellular serine/threonine kinase protein kinase B (also called AKT), which occurs as a result of a single trigger reaction or compound. | 263 | 4 | 8.23E-05 | 0.006435 | IGF2:INS:IL1B:KDR |
| POSITIVE REGULATION OF GROWTH | Any process that activates or increases the rate or extent of growth, the increase in size or mass of all or part of an organism. | 266 | 4 | 8.60E-05 | 0.006565 | IGF2:INS:CCNB1:DDX39B |
| REGULATION OF CARBOHYDRATE BIOSYNTHETIC PROCESS | Any process that modulates the frequency, rate or extent of the chemical reactions and pathways resulting in the formation of carbohydrates. | 97 | 3 | 8.66E-05 | 0.006565 | IGF2:INS:GSK3A |
| REGULATION OF CARDIAC MUSCLE TISSUE DEVELOPMENT | Any process that modulates the frequency, rate or extent of cardiac muscle tissue development. | 97 | 3 | 8.66E-05 | 0.006565 | GSK3A:CCNB1:DDX39B |
| POSITIVE REGULATION OF MAPK CASCADE | Any process that activates or increases the frequency, rate or extent of signal transduction mediated by the MAPK cascade. | 537 | 5 | 0.0001 | 0.007466 | IGF2:INS:IL1B:KDR:JAK2 |

| | | | | | | |
|---|---|---|---|---|---|---|
| AEROBIC ELECTRON TRANSPORT CHAIN | A process in which a series of electron carriers operate together to transfer electrons from donors such as NADH and FADH2 to oxygen to generate a transmembrane electrochemical gradient. | 17 | 2 | 0.000102 | 0.007466 | COX8A:COX7A2L |
| NITRIC OXIDE SYNTHASE BIOSYNTHETIC PROCESS | The chemical reactions and pathways resulting in the formation of a nitric-oxide synthase, an enzyme which catalyzes the reaction L-arginine + n NADPH + n H+ + m O2 = citrulline + nitric oxide + n NADP+. | 17 | 2 | 0.000102 | 0.007466 | KDR:JAK2 |
| STRIATED MUSCLE CELL DIFFERENTIATION | The process in which a relatively unspecialized cell acquires specialized features of a striated muscle cell; striated muscle fibers are divided by transverse bands into striations, and cardiac and voluntary muscle are types of striated muscle. | 279 | 4 | 0.000103 | 0.00752 | IGF2:GSK3A:CCNB1:DDX39B |
| MITOTIC NUCLEAR DIVISION | A mitotic cell cycle process comprising the steps by which the nucleus of a eukaryotic cell divides; the process involves condensation of chromosomal DNA into a highly compacted form. Canonically, mitosis produces two daughter nuclei whose chromosome complement is identical to that of the mother cell. | 280 | 4 | 0.000105 | 0.00755 | IGF2:INS:IL1B:CCNB1 |
| POSITIVE REGULATION OF SMALL MOLECULE METABOLIC PROCESS | Any process that activates or increases the frequency, rate or extent of a small molecule metabolic process. | 105 | 3 | 0.00011 | 0.007742 | IGF2:INS:IL1B |
| HEART GROWTH | The increase in size or mass of the heart. | 105 | 3 | 0.00011 | 0.007742 | GSK3A:CCNB1:DDX39B |
| INTRINSIC APOPTOTIC SIGNALING PATHWAY | A series of molecular signals in which an intracellular signal is conveyed to trigger the apoptotic death of a cell. The pathway starts with reception of an intracellular signal (e.g. DNA damage, endoplasmic reticulum stress, oxidative stress etc.), and ends when the | 284 | 4 | 0.000111 | 0.007742 | INS:NDUFS3:MYBBP1A:JAK2 |

| | | | | | | |
|---|---|---|---|---|---|---|
| | execution phase of apoptosis is triggered. The intrinsic apoptotic signaling pathway is crucially regulated by permeabilization of the mitochondrial outer membrane (MOMP). | | | | | |
| POLYSACCHARIDE METABOLIC PROCESS | The chemical reactions and pathways involving a polysaccharide, a polymer of many (typically more than 10) monosaccharide residues linked glycosidically. [ISBN:0198547684] | 106 | 3 | 0.000113 | 0.007742 | IGF2:INS:GSK3A |
| REGULATION OF OXIDOREDUCTASE ACTIVITY | Any process that modulates the frequency, rate or extent of oxidoreductase activity, the catalysis of an oxidation-reduction (redox) reaction, a reversible chemical reaction in which the oxidation state of an atom or atoms within a molecule is altered. One substrate acts as a hydrogen or electron donor and becomes oxidized, while the other acts as hydrogen or electron acceptor and becomes reduced. | 106 | 3 | 0.000113 | 0.007742 | INS:IL1B:ACVR2A |
| POSITIVE REGULATION OF GLYCOGEN METABOLIC PROCESS | Any process that activates or increases the frequency, rate or extent of the chemical reactions and pathways involving glycogen. | 18 | 2 | 0.000114 | 0.007742 | IGF2:INS |
| ORGANOPHOSPHATE BIOSYNTHETIC PROCESS | The chemical reactions and pathways resulting in the biosynthesis of deoxyribose phosphate, the phosphorylated sugar 2-deoxy-erythro-pentose. | 553 | 5 | 0.000115 | 0.007742 | CTPS1:PAPSS2:PFAS:PPAT:PAICS |
| REGULATION OF ORGAN GROWTH | Any process that modulates the frequency, rate or extent of growth of an organ of an organism. | 109 | 3 | 0.000122 | 0.008184 | GSK3A:CCNB1:DDX39B |
| RESPONSE TO LIPID | Any process that results in a change in state or activity of a cell or an organism (in terms of movement, secretion, enzyme production, gene expression, etc.) as a result of a lipid stimulus. | 915 | 6 | 0.000129 | 0.008562 | CYP1A2:IL1B:CCNB1:NFKBIL1:JAK2:GLDC |

| | | | | | | |
|---|---|---|---|---|---|---|
| POSITIVE REGULATION OF KINASE ACTIVITY | Any process that activates or increases the frequency, rate or extent of kinase activity, the catalysis of the transfer of a phosphate group, usually from ATP, to a substrate molecule. | 569 | 5 | 0.000131 | 0.008608 | IGF2:INS:IL1B:CCNB1:JAK2 |
| ORGANONITROGEN COMPOUND BIOSYNTHETIC PROCESS | The chemical reactions and pathways resulting in the formation of organonitrogen compound. | 1814 | 8 | 0.000135 | 0.008759 | CTPS1:PAPSS2:PFAS:IL1B:GSS:PPAT:PAICS:DDX39B |
| REGULATION OF PROTEIN MODIFICATION PROCESS | Any process that modulates the frequency, rate or extent of the covalent alteration of one or more amino acid residues within a protein. | 1826 | 8 | 0.000141 | 0.009088 | IGF2:INS:GSK3A:IL1B:ACVR2A:KDR:CCNB1:JAK2 |
| REGULATION OF CELLULAR CARBOHYDRATE METABOLIC PROCESS | Any process that modulates the rate, extent or frequency of the chemical reactions and pathways involving carbohydrates, any of a group of organic compounds based of the general formula Cx(H2O)y, as carried out by individual cells. | 115 | 3 | 0.000144 | 0.009164 | IGF2:INS:GSK3A |
| CARBOHYDRATE DERIVATIVE METABOLIC PROCESS | The chemical reactions and pathways involving carbohydrate derivative. | 934 | 6 | 0.000145 | 0.009164 | CTPS1:PAPSS2:PFAS:IL1B:PPAT:PAICS |
| REGULATION OF CELL DIFFERENTIATION | Any process that modulates the frequency, rate or extent of cell differentiation, the process in which relatively unspecialized cells acquire specialized structural and functional features. | 1844 | 8 | 0.000151 | 0.009408 | IGF2:INS:GSK3A:IL1B:ACVR2A:KDR:DDX39B:JAK2 |
| CARBOHYDRATE TRANSMEMBRANE TRANSPORT | The process in which a carbohydrate is transported across a membrane. | 117 | 3 | 0.000151 | 0.009408 | INS:GSK3A:IL1B |
| GLUCOSE METABOLIC PROCESS | The chemical reactions and pathways involving glucose, the aldohexose gluco-hexose. D-glucose is dextrorotatory and is sometimes known as dextrose; it is an important source of energy for living | 118 | 3 | 0.000155 | 0.009507 | IGF2:INS:GSK3A |

| | | | | | | |
|---|---|---|---|---|---|---|
| | organisms and is found free as well as combined in homo- and hetero-oligosaccharides and polysaccharides. | | | | | |
| NEGATIVE REGULATION OF GLUCOSE TRANSMEMBRANE TRANSPORT | Any process that decreases the frequency, rate or extent of glucose transport across a membrane. Glucose transport is the directed movement of the hexose monosaccharide glucose into, out of or within a cell, or between cells, by means of some agent such as a transporter or pore. | 21 | 2 | 0.000157 | 0.009507 | GSK3A:IL1B |
| GLUTAMINE METABOLIC PROCESS | The chemical reactions and pathways involving glutamine, 2-amino-4-carbamoylbutanoic acid. | 21 | 2 | 0.000157 | 0.009507 | CTPS1:PFAS |
| REGULATION OF TRANSFERASE ACTIVITY | Any process that modulates the frequency, rate or extent of transferase activity, the catalysis of the transfer of a group, e.g. a methyl group, glycosyl group, acyl group, phosphorus-containing, or other groups, from one compound (generally regarded as the donor) to another compound (generally regarded as the acceptor). Transferase is the systematic name for any enzyme of EC class 2. | 956 | 6 | 0.000164 | 0.009888 | IGF2:INS:GSK3A:IL1B:CCNB1:JAK2 |
| REGULATION OF PHOSPHATIDYLINOSITOL 3 KINASE SIGNALING | Any process that modulates the frequency, rate or extent of signal transduction mediated by the phosphatidylinositol 3-kinase cascade. | 122 | 3 | 0.000171 | 0.010118 | INS:KDR:JAK2 |
| NEGATIVE REGULATION OF LIPID CATABOLIC PROCESS | Any process that stops, prevents, or reduces the frequency, rate or extent of the chemical reactions and pathways resulting in the breakdown of lipids. | 22 | 2 | 0.000172 | 0.010118 | INS:IL1B |
| REGULATION OF CELL GROWTH INVOLVED IN | Any process that modulates the rate, frequency, or extent of the growth of a cardiac muscle cell, where growth contributes to the progression of the cell over | 22 | 2 | 0.000172 | 0.010118 | GSK3A:DDX39B |

| | | | | | | |
|---|---|---|---|---|---|---|
| CARDIAC MUSCLE CELL DEVELOPMENT | time from its initial formation to its mature state. | | | | | |
| POSITIVE REGULATION OF CELL DIFFERENTIATION | Any process that activates or increases the frequency, rate or extent of cell differentiation. | 970 | 6 | 0.000178 | 0.010359 | INS:IL1B:ACVR2A:KDR:DDX39B:JAK2 |
| MUSCLE ADAPTATION | A process in which muscle adapts, with consequent modifications to structural and/or functional phenotypes, in response to a stimulus. Stimuli include contractile activity, loading conditions, substrate supply, and environmental factors. These adaptive events occur in both muscle fibers and associated structures (motoneurons and capillaries), and they involve alterations in regulatory mechanisms, contractile properties and metabolic capacities. | 125 | 3 | 0.000184 | 0.010627 | GSK3A:IL1B:DDX39B |
| GROWTH | The increase in size or mass of an entire organism, a part of an organism or a cell. | 979 | 6 | 0.000187 | 0.010649 | IGF2:INS:NDUFS3:GSK3A:CCNB1:DDX39B |
| CARDIAC MUSCLE CELL DIFFERENTIATION | The process in which a cardiac muscle precursor cell acquires specialized features of a cardiac muscle cell. Cardiac muscle cells are striated muscle cells that are responsible for heart contraction. | 126 | 3 | 0.000188 | 0.010649 | GSK3A:CCNB1:DDX39B |
| POSITIVE REGULATION OF CELLULAR RESPONSE TO INSULIN STIMULUS | Any process that activates or increases the frequency, rate or extent of cellular response to insulin stimulus. | 23 | 2 | 0.000188 | 0.010649 | IGF2:INS |
| POSITIVE REGULATION OF PROTEIN SERINE THREONINE KINASE ACTIVITY | Any process that activates or increases the frequency, rate or extent of cellular response to insulin stimulus. | 331 | 4 | 0.000199 | 0.011163 | IGF2:IL1B:CCNB1:JAK2 |

| | | | | | | |
|---|---|---|---|---|---|---|
| POSITIVE REGULATION OF GLUCOSE METABOLIC PROCESS | Any process that increases the rate, frequency or extent of glucose metabolism. Glucose metabolic processes are the chemical reactions and pathways involving glucose, the aldohexose gluco-hexose. | 24 | 2 | 0.000205 | 0.011435 | IGF2:INS |
| RESPONSE TO ACID CHEMICAL | Any process that results in a change in state or activity of a cell or an organism (in terms of movement, secretion, enzyme production, gene expression, etc.) as a result of a stimulus by the chemical structure of the anion portion of a dissociated acid (rather than the acid acting as a proton donor). The acid chemical may be in gaseous, liquid or solid form. | 336 | 4 | 0.000211 | 0.011642 | GSS:KDR:CCNB1:GLDC |
| RESPONSE TO MOLECULE OF BACTERIAL ORIGIN | Any process that results in a change in state or activity of an organism (in terms of movement, secretion, enzyme production, gene expression, etc.) as a result of a stimulus by molecules of bacterial origin such as peptides derived from bacterial flagellin. | 337 | 4 | 0.000213 | 0.011646 | CYP1A2:IL1B:NFKBIL1:JAK2 |
| POSITIVE REGULATION OF INTRACELLULAR SIGNAL TRANSDUCTION | Any process that activates or increases the frequency, rate or extent of intracellular signal transduction. | 1005 | 6 | 0.000215 | 0.011646 | IGF2:INS:GSK3A:IL1B:KDR:JAK2 |
| REGULATION OF DEVELOPMENTAL GROWTH | Any process that modulates the frequency, rate or extent of developmental growth. | 338 | 4 | 0.000215 | 0.011646 | IGF2:GSK3A:CCNB1:DDX39B |
| PROTEIN PHOSPHORYLATION | Any process that increases the rate, frequency, or extent of protein serine/threonine kinase activity. | 1949 | 8 | 0.000221 | 0.011841 | IGF2:INS:GSK3A:IL1B:ACVR2A:KDR:CCNB1:JAK2 |
| POSITIVE REGULATION OF GENE EXPRESSION | Any process that increases the frequency, rate or extent of gene expression. Gene expression is the process in which a gene's coding sequence is converted into a mature gene product (protein or RNA). | 1955 | 8 | 0.000225 | 0.012005 | IGF2:INS:MYBBP1A:GSK3A:IL1B:ACVR2A:CCNB1:DDX39B |

| | | | | | | |
|---|---|---|---|---|---|---|
| REGULATION OF SMALL MOLECULE METABOLIC PROCESS | Any process that modulates the rate, frequency or extent of a small molecule metabolic process. | 343 | 4 | 0.000228 | 0.01205 | IGF2:INS:GSK3A:IL1B |
| REGULATION OF OXIDATIVE PHOSPHORYLATION | Any process that modulates the frequency, rate or extent of the chemical reactions and pathways resulting in the phosphorylation of ADP to ATP that accompanies the oxidation of a metabolite through the operation of the respiratory chain. Oxidation of compounds establishes a proton gradient across the membrane, providing the energy for ATP synthesis. | 26 | 2 | 0.000242 | 0.012682 | COX7A2L:CCNB1 |
| REGULATION OF HISTONE MODIFICATION | Any process that modulates the frequency, rate or extent of the covalent alteration of a histone. | 140 | 3 | 0.000256 | 0.013362 | IGF2:IL1B:CCNB1 |
| PHOSPHATIDYLINOSITOL 3 KINASE SIGNALING | A series of reactions within the signal-receiving cell, mediated by the intracellular phosphatidylinositol 3-kinase (PI3K). Many cell surface receptor linked signaling pathways signal through PI3K to regulate numerous cellular functions. | 146 | 3 | 0.00029 | 0.015008 | INS:KDR:JAK2 |
| MUSCLE CELL DIFFERENTIATION | The process in which a relatively unspecialized cell acquires specialized features of a muscle cell. | 367 | 4 | 0.000295 | 0.015144 | IGF2:GSK3A:CCNB1:DDX39B |
| POSITIVE REGULATION OF MONOOXYGENASE ACTIVITY | Any process that activates or increases the activity of a monooxygenase. | 29 | 2 | 0.000301 | 0.015296 | INS:IL1B |
| CARBOHYDRATE TRANSPORT | The directed movement of carbohydrate into, out of or within a cell, or between cells, by means of some agent such as a transporter or pore. Carbohydrates are a group of organic compounds based of the general formula Cx(H2O)y. | 148 | 3 | 0.000302 | 0.015296 | INS:GSK3A:IL1B |

| | | | | | | |
|---|---|---|---|---|---|---|
| PROTON TRANSMEMBRANE TRANSPORT | The directed movement of a proton across a membrane. | 152 | 3 | 0.000326 | 0.016426 | COX8A:COX7A2L:ATP6V1G2 |
| MONOSACCHARIDE METABOLIC PROCESS | The chemical reactions and pathways involving monosaccharides, the simplest carbohydrates. They are polyhydric alcohols containing either an aldehyde or a keto group and between three to ten or more carbon atoms. They form the constitutional repeating units of oligo- and polysaccharides. | 153 | 3 | 0.000333 | 0.016631 | IGF2:INS:GSK3A |
| REGULATION OF MUSCLE ORGAN DEVELOPMENT | Any process that modulates the frequency, rate or extent of muscle development. | 154 | 3 | 0.000339 | 0.016837 | GSK3A:CCNB1:DDX39B |
| CELLULAR PROTEIN CONTAINING COMPLEX ASSEMBLY | Any process that modulates the frequency, rate or extent of protein complex assembly. | 1112 | 6 | 0.00037 | 0.01824 | NDUFS3:COX7A2L:NDUFA6:DDX39B:NDUFA5:JAK2 |
| CARDIOCYTE DIFFERENTIATION | The process in which a relatively unspecialized cell acquires the specialized structural and/or functional features of a cell that will form part of the cardiac organ of an individual. | 159 | 3 | 0.000372 | 0.01824 | GSK3A:CCNB1:DDX39B |
| PHYSIOLOGICAL CARDIAC MUSCLE HYPERTROPHY | The enlargement or overgrowth of all or part of the heart muscle due to an increase in size of cardiac muscle cells without cell division. This process contributes to the developmental growth of the heart. | 33 | 2 | 0.000391 | 0.018871 | GSK3A:DDX39B |
| REGULATION OF CELL CYCLE CHECKPOINT | Any process that modulates the frequency, rate or extent of cell cycle checkpoint. | 33 | 2 | 0.000391 | 0.018871 | CCNB1:DDX39B |
| SMALL MOLECULE CATABOLIC PROCESS | The chemical reactions and pathways resulting in the breakdown of small molecules, any low molecular weight, monomeric, non-encoded molecule. | 396 | 4 | 0.000393 | 0.018871 | ALDH2:GSK3A:PPAT:GLDC |

| | | | | | | |
|---|---|---|---|---|---|---|
| POSITIVE REGULATION OF CELL ADHESION | Any process that activates or increases the frequency, rate or extent of cell adhesion. | 397 | 4 | 0.000397 | 0.018927 | IGF2:IL1B:KDR:JAK2 |
| REGULATION OF MAPK CASCADE | Any process that modulates the frequency, rate or extent of signal transduction mediated by the MAPK cascade. | 751 | 5 | 0.000471 | 0.022354 | IGF2:INS:IL1B:KDR:JAK2 |
| POSITIVE REGULATION OF CATABOLIC PROCESS | Any process that activates or increases the frequency, rate or extent of the chemical reactions and pathways resulting in the breakdown of substances. | 418 | 4 | 0.000482 | 0.022686 | INS:GSK3A:IL1B:KDR |
| NEGATIVE REGULATION OF CELLULAR RESPONSE TO INSULIN STIMULUS | Any process that stops, prevents or reduces the frequency, rate or extent of cellular response to insulin stimulus. | 37 | 2 | 0.000492 | 0.022909 | GSK3A:IL1B |
| REGULATION OF MUSCLE CELL DIFFERENTIATION | Any process that modulates the frequency, rate or extent of muscle cell differentiation. | 175 | 3 | 0.000492 | 0.022909 | IGF2:GSK3A:DDX39B |
| REGULATION OF DNA BINDING TRANSCRIPTION FACTOR ACTIVITY | Any process that modulates the frequency, rate or extent of the activity of a transcription factor, any factor involved in the initiation or regulation of transcription. | 425 | 4 | 0.000512 | 0.023691 | INS:IL1B:NFKBIL1:JAK2 |
| MUSCLE CELL DEVELOPMENT | The process whose specific outcome is the progression of a muscle cell over time, from its formation to the mature structure. Muscle cell development does not include the steps involved in committing an unspecified cell to the muscle cell fate. | 178 | 3 | 0.000517 | 0.02377 | GSK3A:CCNB1:DDX39B |
| POSITIVE REGULATION OF CELLULAR COMPONENT ORGANIZATION | Any process that activates or increases the frequency, rate or extent of a process involved in the formation, arrangement of constituent parts, or disassembly of cell structures, including the plasma membrane and any external encapsulating structures such as the cell wall and cell envelope. | 1188 | 6 | 0.000525 | 0.023863 | IGF2:INS:GSK3A:IL1B:KDR:CCNB1 |

| | | | | | | |
|---|---|---|---|---|---|---|
| T CELL PROLIFERATION | The expansion of a T cell population by cell division. Follows T cell activation. | 179 | 3 | 0.000526 | 0.023863 | CTPS1:IGF2:IL1B |
| POSITIVE REGULATION OF DEVELOPMENTAL GROWTH | Any process that activates, maintains or increases the rate of developmental growth. | 181 | 3 | 0.000543 | 0.024177 | IGF2:CCNB1:DDX39B |
| REGULATION OF CELL POPULATION PROLIFERATION | Any process that modulates the frequency, rate or extent of cell proliferation. | 1684 | 7 | 0.000546 | 0.024177 | IGF2:INS:IL1B:KDR:CCNB1:DDX39B:JAK2 |
| REGULATION OF CHROMATIN ORGANIZATION | Any process that modulates the frequency, rate or extent of chromatin organization. | 182 | 3 | 0.000552 | 0.024177 | IGF2:IL1B:CCNB1 |
| INOSITOL LIPID MEDIATED SIGNALING | A series of molecular signals in which a cell uses an inositol-containing lipid to convert a signal into a response. Inositol lipids include the phosphoinositides (phosphatidylinositol and its phosphorylated derivatives), ceramides containing inositol, and inositol glycolipids. | 182 | 3 | 0.000552 | 0.024177 | INS:KDR:JAK2 |
| REGULATION OF CELL CYCLE | Any process that modulates the rate or extent of progression through the cell cycle. | 1201 | 6 | 0.000556 | 0.024177 | IGF2:INS:MYBBP1A:IL1B:CCNB1:DDX39B |
| REGULATION OF RESPONSE TO EXTERNAL STIMULUS | Any process that modulates the frequency, rate or extent of a response to an external stimulus. | 779 | 5 | 0.000557 | 0.024177 | INS:IL1B:KDR:NFKBIL1:JAK2 |
| GLAND DEVELOPMENT | The process whose specific outcome is the progression of a gland over time, from its formation to the mature structure. A gland is an organ specialised for secretion. | 435 | 4 | 0.000559 | 0.024177 | IGF2:PPAT:DDX39B:JAK2 |
| RESPONSE TO OXIDATIVE STRESS | Any process that results in a change in state or activity of a cell or an organism (in terms of movement, secretion, enzyme production, gene expression, etc.) as a result of oxidative stress, a state often resulting from exposure to high levels of reactive oxygen species, e.g. | 435 | 4 | 0.000559 | 0.024177 | INS:GSS:NDUFA6:JAK2 |

| | | | | | | |
|---|---|---|---|---|---|---|
| | superoxide anions, hydrogen peroxide (H2O2), and hydroxyl radicals. | | | | | |
| POSITIVE REGULATION OF ESTABLISHMENT OF PROTEIN LOCALIZATION | Any process that activates or increases the frequency, rate or extent of establishment of protein localization. | 436 | 4 | 0.000564 | 0.024243 | INS:GSK3A:IL1B:JAK2 |
| REGULATION OF CELL DEATH | Any process that modulates the rate or frequency of cell death. Cell death is the specific activation or halting of processes within a cell so that its vital functions markedly cease, rather than simply deteriorating gradually over time, which culminates in cell death. | 1697 | 7 | 0.000572 | 0.024443 | INS:NDUFS3:MYBBP1A:GSK3A:IL1B:KDR:JAK2 |
| REGULATION OF CELL CYCLE PROCESS | Any process that modulates a cellular process that is involved in the progression of biochemical and morphological phases and events that occur in a cell during successive cell replication or nuclear replication events. | 785 | 5 | 0.000576 | 0.024484 | IGF2:INS:MYBBP1A:IL1B:CCNB1 |
| PEPTIDYL AMINO ACID MODIFICATION | The alteration of an amino acid residue in a peptide. | 1219 | 6 | 0.000602 | 0.025419 | IGF2:GSK3A:IL1B:KDR:CCNB1:JAK2 |
| POSITIVE REGULATION OF NITRIC OXIDE BIOSYNTHETIC PROCESS | Any process that activates or increases the frequency, rate or extent of the chemical reactions and pathways resulting in the formation of nitric oxide. | 42 | 2 | 0.000634 | 0.026642 | IL1B:JAK2 |
| REGENERATION | The regrowth of a lost or destroyed body part, such as an organ or tissue. This process may occur via renewal, repair, and/or growth alone (i.e. increase in size or mass). | 192 | 3 | 0.000645 | 0.026813 | PPAT:CCNB1:JAK2 |
| REGULATION OF CELLULAR CATABOLIC PROCESS | Any process that modulates the frequency, rate or extent of the chemical reactions and pathways resulting in the breakdown of substances, carried out by individual cells. | 805 | 5 | 0.000646 | 0.026813 | INS:GSK3A:IL1B:KDR:ATP6V1G2 |

| | | | | | | |
|---|---|---|---|---|---|---|
| INTRINSIC APOPTOTIC SIGNALING PATHWAY IN RESPONSE TO OXIDATIVE STRESS | A series of molecular signals in which an intracellular signal is conveyed to trigger the apoptotic death of a cell. The pathway is induced in response to oxidative stress, a state often resulting from exposure to high levels of reactive oxygen species, and ends when the execution phase of apoptosis is triggered. | 43 | 2 | 0.000665 | 0.027456 | INS:JAK2 |
| T CELL ACTIVATION | The change in morphology and behavior of a mature or immature T cell resulting from exposure to a mitogen, cytokine, chemokine, cellular ligand, or an antigen for which it is specific. | 457 | 4 | 0.000672 | 0.027614 | CTPS1:IGF2:INS:IL1B |
| COVALENT CHROMATIN MODIFICATION | The alteration of DNA or protein in chromatin by the covalent addition or removal of chemical groups. | 458 | 4 | 0.000678 | 0.027685 | IGF2:IL1B:CCNB1:JAK2 |
| ORGAN GROWTH | The increase in size or mass of an organ. Organs are commonly observed as visibly distinct structures, but may also exist as loosely associated clusters of cells that function together as to perform a specific function. | 196 | 3 | 0.000685 | 0.027797 | GSK3A:CCNB1:DDX39B |
| POSITIVE REGULATION OF CELLULAR CARBOHYDRATE METABOLIC PROCESS | Any process that increases the rate, extent or frequency of the chemical reactions and pathways involving carbohydrates, any of a group of organic compounds based of the general formula Cx(H2O)y, as carried out by individual cells. | 44 | 2 | 0.000696 | 0.028039 | IGF2:INS |
| REGULATION OF ORGANELLE ORGANIZATION | Any process that modulates the frequency, rate or extent of a process involved in the formation, arrangement of constituent parts, or disassembly of an organelle. | 1254 | 6 | 0.000698 | 0.028039 | IGF2:INS:GSK3A:IL1B:KDR:CCNB1 |
| REGULATION OF CARBOHYDRATE METABOLIC PROCESS | Any process that modulates the frequency, rate or extent of the chemical reactions and pathways involving carbohydrates. | 199 | 3 | 0.000715 | 0.028468 | IGF2:INS:GSK3A |

| | | | | | | |
|---|---|---|---|---|---|---|
| ORGANONITROGEN COMPOUND CATABOLIC PROCESS | The chemical reactions and pathways resulting in the breakdown of organonitrogen compound. | 1263 | 6 | 0.000725 | 0.028468 | INS:GSK3A:IL1B:PPAT:CCNB1:GLDC |
| CELLULAR CARBOHYDRATE CATABOLIC PROCESS | The chemical reactions and pathways resulting in the breakdown of carbohydrates, any of a group of organic compounds based of the general formula Cx(H2O)y, as carried out by individual cells. | 45 | 2 | 0.000728 | 0.028468 | INS:GSK3A |
| NEGATIVE REGULATION OF CARBOHYDRATE METABOLIC PROCESS | Any process that stops, prevents, or reduces the frequency, rate or extent of the chemical reactions and pathways involving carbohydrate. | 45 | 2 | 0.000728 | 0.028468 | INS:GSK3A |
| REGULATION OF ACUTE INFLAMMATORY RESPONSE | Any process that modulates the frequency, rate, or extent of an acute inflammatory response. | 45 | 2 | 0.000728 | 0.028468 | INS:IL1B |
| CELL GROWTH | The process in which a cell irreversibly increases in size over time by accretion and biosynthetic production of matter similar to that already present. | 468 | 4 | 0.000735 | 0.028578 | INS:NDUFS3:GSK3A:DDX39B |
| REGULATION OF CARDIAC MUSCLE CELL DIFFERENTIATION | Any process that modulates the frequency, rate or extent of cardiac muscle cell differentiation. | 46 | 2 | 0.000761 | 0.029433 | GSK3A:DDX39B |
| REGULATION OF PEPTIDE HORMONE SECRETION | Any process that modulates the rate, frequency, or extent of the regulated release of a peptide hormone from secretory granules. | 205 | 3 | 0.00078 | 0.029999 | INS:IL1B:JAK2 |
| POSITIVE REGULATION OF VASCULAR SMOOTH MUSCLE CELL PROLIFERATION | Any process that activates or increases the frequency, rate or extent of vascular smooth muscle cell proliferation. | 47 | 2 | 0.000794 | 0.030247 | DDX39B:JAK2 |
| ACUTE PHASE RESPONSE | An acute inflammatory response that involves non-antibody proteins whose | 47 | 2 | 0.000794 | 0.030247 | INS:IL1B |

| | | | | | | |
|---|---|---|---|---|---|---|
| | concentrations in the plasma increase in response to infection or injury of homeothermic animals. | | | | | |
| CARBOHYDRATE BIOSYNTHETIC PROCESS | The chemical reactions and pathways resulting in the formation of carbohydrates, any of a group of organic compounds based of the general formula Cx(H2O)y. | 208 | 3 | 0.000813 | 0.030802 | IGF2:INS:GSK3A |
| OSTEOBLAST DIFFERENTIATION | The process whereby a relatively unspecialized cell acquires the specialized features of an osteoblast, a mesodermal or neural crest cell that gives rise to bone. | 210 | 3 | 0.000836 | 0.031504 | IGF2:MYBBP1A:ACVR2A |
| POSITIVE REGULATION OF SIGNALING | Any process that activates, maintains or increases the frequency, rate or extent of a signaling process. | 1813 | 7 | 0.000847 | 0.031761 | IGF2:INS:GSK3A:IL1B:ACVR2A:KDR:JAK2 |
| REGULATION OF KINASE ACTIVITY | Any process that modulates the frequency, rate or extent of kinase activity, the catalysis of the transfer of a phosphate group, usually from ATP, to a substrate molecule. | 857 | 5 | 0.000856 | 0.031924 | IGF2:INS:IL1B:CCNB1:JAK2 |
| REGULATION OF INTRACELLULAR SIGNAL TRANSDUCTION | Any process that modulates the frequency, rate or extent of intracellular signal transduction. | 1824 | 7 | 0.000878 | 0.032581 | IGF2:INS:NDUFS3:GSK3A:IL1B:KDR:JAK2 |
| CELL CYCLE | The progression of biochemical and morphological phases and events that occur in a cell during successive cell replication or nuclear replication events. Canonically, the cell cycle comprises the replication and segregation of genetic material followed by the division of the cell, but in endocycles or syncytial cells nuclear replication or nuclear division may not be followed by cell division. | 1834 | 7 | 0.000906 | 0.033478 | IGF2:INS:MYBBP1A:IL1B:PPAT:CCNB1:DDX39B |
| POSITIVE REGULATION OF HEART GROWTH | Any process that increases the rate or extent of heart growth. Heart growth is the increase in size or mass of the heart. | 51 | 2 | 0.000935 | 0.034351 | CCNB1:DDX39B |

| | | | | | | |
|---|---|---|---|---|---|---|
| REGULATION OF LIPID CATABOLIC PROCESS | Any process that modulates the frequency, rate, or extent of the chemical reactions and pathways resulting in the breakdown of lipids. | 52 | 2 | 0.000972 | 0.035296 | INS:IL1B |
| POSITIVE REGULATION OF REACTIVE OXYGEN SPECIES BIOSYNTHETIC PROCESS | Any process that activates or increases the frequency, rate or extent of reactive oxygen species biosynthetic process. | 52 | 2 | 0.000972 | 0.035296 | IL1B:JAK2 |
| REGULATION OF PROTEIN SERINE THREONINE KINASE ACTIVITY | Any process that modulates the rate, frequency, or extent of protein serine/threonine kinase activity. | 505 | 4 | 0.000975 | 0.035296 | IGF2:IL1B:CCNB1:JAK2 |
| MULTI MULTICELLULAR ORGANISM PROCESS | A multicellular organism process which involves another multicellular organism of the same or different species. | 222 | 3 | 0.000981 | 0.035351 | IL1B:ACVR2A:PPAT |
| NEGATIVE REGULATION OF APOPTOTIC SIGNALING PATHWAY | Any process that stops, prevents, or reduces the frequency, rate or extent of cell death by apoptotic process. | 223 | 3 | 0.000994 | 0.035637 | INS:NDUFS3:IL1B |
| EXTRINSIC APOPTOTIC SIGNALING PATHWAY | A series of molecular signals in which a signal is conveyed from the cell surface to trigger the apoptotic death of a cell. The pathway starts with either a ligand binding to a cell surface receptor, or a ligand being withdrawn from a cell surface receptor (e.g. in the case of signaling by dependence receptors), and ends when the execution phase of apoptosis is triggered. | 224 | 3 | 0.001007 | 0.035924 | GSK3A:IL1B:JAK2 |
| CARDIAC MUSCLE TISSUE DEVELOPMENT | The process whose specific outcome is the progression of cardiac muscle over time, from its formation to the mature structure. | 226 | 3 | 0.001033 | 0.036677 | GSK3A:CCNB1:DDX39B |

| | | | | | | |
|---|---|---|---|---|---|---|
| NEGATIVE REGULATION OF INTRACELLULAR SIGNAL TRANSDUCTION | Any process that stops, prevents or reduces the frequency, rate or extent of intracellular signal transduction. | 521 | 4 | 0.001094 | 0.038658 | INS:NDUFS3:GSK3A:IL1B |
| CELL CYCLE PROCESS | The cellular process that ensures successive accurate and complete genome replication and chromosome segregation. | 1370 | 6 | 0.001106 | 0.038792 | IGF2:INS:MYBBP1A:IL1B:PPAT:CCNB1 |
| MUSCLE CELL PROLIFERATION | The expansion of a muscle cell population by cell division. | 232 | 3 | 0.001114 | 0.038792 | CCNB1:DDX39B:JAK2 |
| PIGMENT BIOSYNTHETIC PROCESS | The chemical reactions and pathways resulting in the formation of a pigment, any general or particular coloring matter in living organisms, e.g. melanin. | 56 | 2 | 0.001126 | 0.038792 | PPAT:PAICS |
| POSITIVE REGULATION OF OXIDOREDUCTASE ACTIVITY | Any process that activates or increases the frequency, rate or extent of oxidoreductase activity, the catalysis of an oxidation-reduction (redox) reaction, a reversible chemical reaction in which the oxidation state of an atom or atoms within a molecule is altered. | 56 | 2 | 0.001126 | 0.038792 | INS:IL1B |
| PROTEIN AUTOPHOSPHORYLATION | The phosphorylation by a protein of one or more of its own amino acid residues (cis-autophosphorylation), or residues on an identical protein (trans-autophosphorylation). | 233 | 3 | 0.001128 | 0.038792 | INS:KDR:JAK2 |
| RESPONSE TO ORGANIC CYCLIC COMPOUND | Any process that results in a change in state or activity of a cell or an organism (in terms of movement, secretion, enzyme production, gene expression, etc.) as a result of an organic cyclic compound stimulus. | 912 | 5 | 0.001129 | 0.038792 | CYP1A2:IL1B:CCNB1:JAK2:GLDC |
| LIPOPOLYSACCHARIDE MEDIATED SIGNALING PATHWAY | A series of molecular signals initiated by the binding of a lipopolysaccharide (LPS) to a receptor on the surface of a target cell, and ending with regulation of a downstream cellular process, e.g. transcription. | 57 | 2 | 0.001166 | 0.039868 | IL1B:NFKBIL1 |

| | | | | | | |
|---|---|---|---|---|---|---|
| | Lipopolysaccharides are major components of the outer membrane of Gram-negative bacteria, making them prime targets for recognition by the immune system. | | | | | |
| POSITIVE REGULATION OF CATALYTIC ACTIVITY | Any process that activates or increases the activity of an enzyme. | 1397 | 6 | 0.001223 | 0.041616 | IGF2:INS:GSK3A:IL1B:CCNB1:JAK2 |
| POSITIVE REGULATION OF CARDIAC MUSCLE TISSUE DEVELOPMENT | Any process that activates or increases the frequency, rate or extent of cardiac muscle cell proliferation. | 60 | 2 | 0.001291 | 0.043734 | CCNB1:DDX39B |
| APOPTOTIC PROCESS | A programmed cell death process which begins when a cell receives an internal (e.g. DNA damage) or external signal (e.g. an extracellular death ligand), and proceeds through a series of biochemical events (signaling pathway phase) which trigger an execution phase. The execution phase is the last step of an apoptotic process, and is typically characterized by rounding-up of the cell, retraction of pseudopodes, reduction of cellular volume (pyknosis), chromatin condensation, nuclear fragmentation (karyorrhexis), plasma membrane blebbing and fragmentation of the cell into apoptotic bodies. When the execution phase is completed, the cell has died. | 1956 | 7 | 0.001321 | 0.044535 | INS:NDUFS3:MYBBP1A:GSK3A:IL1B:KDR:JAK2 |
| REGULATION OF CARDIOCYTE DIFFERENTIATION | Any process that modulates the frequency, rate or extent of cardiocyte differentiation. [GO_REF:0000058, GOC:bc, GOC:BHF, GOC:BHF_miRNA, GOC:TermGenie, | 61 | 2 | 0.001334 | 0.044618 | GSK3A:DDX39B |
| WOUND HEALING | The series of events that restore integrity to a damaged tissue, following an injury. | 550 | 4 | 0.001336 | 0.044618 | PAPSS2:INS:CCNB1:JAK2 |

| | | | | | | |
|---|---|---|---|---|---|---|
| PEPTIDE HORMONE SECRETION | The regulated release of a peptide hormone from a cell. | 248 | 3 | 0.001349 | 0.044829 | INS:IL1B:JAK2 |
| POSITIVE REGULATION OF BIOSYNTHETIC PROCESS | Any process that activates or increases the frequency, rate or extent of the chemical reactions and pathways resulting in the formation of substances. | 1966 | 7 | 0.001361 | 0.044829 | IGF2:INS:IL1B:ACVR2A:KDR:DDX39B:JAK2 |
| RESPONSE TO INORGANIC SUBSTANCE | Any process that results in a change in state or activity of a cell or an organism (in terms of movement, secretion, enzyme production, gene expression, etc.) as a result of an inorganic substance stimulus. | 553 | 4 | 0.001362 | 0.044829 | CYP1A2:GSS:KDR:CCNB1 |
| SIGNAL TRANSDUCTION BY PROTEIN PHOSPHORYLATION | The cellular process in which a signal is conveyed to trigger a change in the activity or state of a cell. Signal transduction begins with reception of a signal (e.g. a ligand binding to a receptor or receptor activation by a stimulus such as light), or for signal transduction in the absence of ligand, signal-withdrawal or the activity of a constitutively active receptor. Signal transduction ends with regulation of a downstream cellular process, e.g. regulation of transcription or regulation of a metabolic process. Signal transduction covers signaling from receptors located on the surface of the cell and signaling via molecules located within the cell. For signaling between cells, signal transduction is restricted to events at and within the receiving cell. | 952 | 5 | 0.001366 | 0.044829 | IGF2:INS:IL1B:KDR:JAK2 |
| POSITIVE REGULATION OF ORGAN GROWTH | Any process that activates or increases the frequency, rate or extent of growth of an organ of an organism. | 62 | 2 | 0.001378 | 0.045014 | CCNB1:DDX39B |
| REGULATION OF CATABOLIC PROCESS | Any process that modulates the frequency, rate, or extent of the chemical reactions and | 959 | 5 | 0.001411 | 0.045852 | INS:GSK3A:IL1B:KDR:ATP6V1G2 |

| | | | | | | |
|---|---|---|---|---|---|---|
| | pathways resulting in the breakdown of substances. | | | | | |
| CARBOHYDRATE METABOLIC PROCESS | The chemical reactions and pathways involving carbohydrates, any of a group of organic compounds based of the general formula Cx(H2O)y. Includes the formation of carbohydrate derivatives by the addition of a carbohydrate residue to another molecule. | 559 | 4 | 0.001418 | 0.045852 | IGF2:INS:ALDH2:GSK3A |
| RESPONSE TO CADMIUM ION | Any process that results in a change in state or activity of a cell or an organism (in terms of movement, secretion, enzyme production, gene expression, etc.) as a result of a cadmium (Cd) ion stimulus. | 63 | 2 | 0.001422 | 0.045852 | CYP1A2:GSS |
| CELLULAR CARBOHYDRATE METABOLIC PROCESS | The chemical reactions and pathways involving carbohydrates, any of a group of organic compounds based of the general formula Cx(H2O)y, as carried out by individual cells. | 260 | 3 | 0.001544 | 0.049341 | IGF2:INS:GSK3A |
| POSITIVE REGULATION OF DNA BINDING TRANSCRIPTION FACTOR ACTIVITY | Any process that activates or increases the frequency, rate or extent of activity of a transcription factor, any factor involved in the initiation or regulation of transcription. | 260 | 3 | 0.001544 | 0.049341 | INS:IL1B:JAK2 |
| POSITIVE REGULATION OF CARBOHYDRATE METABOLIC PROCESS | Any process that activates or increases the frequency, rate or extent of the chemical reactions and pathways involving carbohydrate. | 66 | 2 | 0.00156 | 0.049624 | IGF2:INS |
| POSITIVE REGULATION OF LOCOMOTION | Any process that activates or increases the frequency, rate or extent of locomotion of a cell or organism. | 575 | 4 | 0.001572 | 0.049798 | INS:IL1B:KDR:JAK2 |

Supplementary table 1: GO Biological Processes Database (MsigDB c5). N indicates the total number of genes in the set while n shows the number of genes that SNPs mapped to the 37 genes identified. Gene sets were queried using FUMA GWAS. The adjusted P is an FDR-corrected P-value based on the number of gene sets examined.

| Gene Set | N | n | P-value | adj. P-value | Genes |
|---|---|---|---|---|---|
| Length of menstrual cycle | 8 | 2 | 2.10E-05 | 0.023045 | IGF2:INS-IGF2 |
| Inflammatory bowel disease | 730 | 6 | 3.72E-05 | 0.023045 | ALDH2:ATP6V1G2-DDX39B:DDX39B:ATP6V1G2:NFKBIL1:JAK2 |
| Autism spectrum disorder or schizophrenia | 735 | 6 | 3.87E-05 | 0.023045 | MIR1307:NDUFA6:ATP6V1G2-DDX39B:DDX39B:ATP6V1G2:NFKBIL1 |
| Ulcerative colitis | 465 | 5 | 5.08E-05 | 0.023045 | ATP6V1G2-DDX39B:DDX39B:ATP6V1G2:NFKBIL1:JAK2 |
| Myositis | 15 | 2 | 7.85E-05 | 0.026427 | DDX39B:ATP6V1G2 |
| Estimated glomerular filtration rate | 534 | 5 | 9.75E-05 | 0.026427 | IGF2:INS-IGF2:ALDH2:ACVR2A:GSS |
| Birth weight | 278 | 4 | 0.000102 | 0.026427 | IGF2:INS-IGF2:INS:IL1B |

Supplementary table 2: GWAS Catalogue Reported Genes Database. N indicates the total number of genes in the set while n shows the number of genes that SNPs mapped to the 37 genes identified. Gene sets were queried using FUMA GWAS. The adjusted P is an FDR-corrected P-value based on the number of gene sets examined.

| Gene Set | Brief Description | N | n | P-value | adj. P-value | Genes | Publication | Species |
|---|---|---|---|---|---|---|---|---|
| MOOTHA VOXPHOS | Genes involved in oxidative phosphorylation; based on literature and sequence annotation resources and converted to Affymetrix HG-U133A probe sets. | 87 | 6 | 1.35E-10 | 4.47E-07 | SDHC:NDUFS3:COX8A:COX7A2L:NDUFA6:NDUFA5 | Pubmed 12808457 Authors: Mootha VK,Lindgren CM,Eriksson KF,Subramanian A,Sihag S,Lehar J,Puigserver P,Carlsson E,Ridderstråle M,Laurila E,Houstis N,Daly MJ,Patterson N,Mesirov JP,Golub TR,Tamayo P,Spiegelman B,Lander ES,Hirschhorn JN,Altshuler D,Groop LC | Homo sapiens |
| MOOTHA HUMAN MITODB 6 2002 | Mitochondrial genes; based on literature and sequence annotation resources and converted to Affymetrix HG-U133A probe sets. | 430 | 8 | 2.90E-09 | 4.80E-06 | SDHC:NDUFS3:COX8A:ALDH2:POLG2:COX7A2L:NDUFA6:NDUFA5 | Pubmed 12808457 Authors: Mootha VK,Lindgren CM,Eriksson KF,Subramanian A,Sihag S,Lehar J,Puigserver P,Carlsson E,Ridderstråle M,Laurila E,Houstis N,Daly MJ,Patterson N,Mesirov JP,Golub TR,Tamayo P,Spiegelman B,Lander ES,Hirschhorn JN,Altshuler D,Groop LC | Homo sapiens |
| MOOTHA MITOCHONDRIA | Mitochondrial genes | 449 | 7 | 1.07E-07 | 0.000118 | SDHC:NDUFS3:COX8A:ALDH2:POLG2:NDUFA6:NDUFA5 | Pubmed 12808457 Authors: Mootha VK,Lindgren CM,Eriksson KF,Subramanian A,Sihag S,Lehar J,Puigserver P,Carlsson E,Ridderstråle M,Laurila E,Houstis N,Daly MJ,Patterson N,Mesirov JP,Golub TR,Tamayo P,Spiegelman B,Lander ES,Hirschhorn JN,Altshuler D,Groop LC | Homo sapiens |
| WONG MITOCHONDRIA GENE MODULE | Genes that comprise the mitochondria gene module | 218 | 5 | 1.31E-06 | 0.001079 | NDUFS3:COX8A:COX7A2L:NDUFA6:NDUFA5 | Pubmed 18199530 Authors: Wong DJ,Nuyten DS,Regev A,Lin M,Adler AS,Segal E,van de Vijver MJ,Chang HY | Homo sapiens |
| DANG BOUND BY MYC | Genes whose promoters are bound by the MYC gene, according to MYC Target Gene Database. | 1059 | 8 | 2.82E-06 | 0.001862 | ALDH2:COX7A2L:NDUFA6:PPAT:PAICS:CCNB1:ATP6V1G2:NFKBIL1 | Pubmed 14519204 Authors: Zeller KI,Jegga AG,Aronow BJ,O'Donnell KA,Dang CV | Homo sapiens |

| | | | | | | | | |
|---|---|---|---|---|---|---|---|---|
| YOSHIMURA MAPK8 TARGETS UP | Genes up-regulated in vascular smooth muscle cells (VSMC) by MAPK8 (JNK1) | 1171 | 8 | 5.91E-06 | 0.003254 | IGF2:INS:COX8A:POLG2:ACVR2A:GSS:NDUFA5:JAK2 | Pubmed 16311603 Authors: Yoshimura K,Aoki H,Ikeda Y,Fujii K,Akiyama N,Furutani A,Hoshii Y,Tanaka N,Ricci R,Ishihara T,Esato K,Hamano K,Matsuzaki M | Rattus norvegicus |
| TENEDINI MEGAKARYOCYTE MARKERS | Genes essential to the development of megakaryocytes, as expressed in normal cells and essential thrombocythemic cells (ET). | 61 | 3 | 2.16E-05 | 0.010193 | IL1B:CCNB1:JAK2 | Pubmed 15271793 Authors: Tenedini E,Fagioli ME,Vianelli N,Tazzari PL,Ricci F,Tagliafico E,Ricci P,Gugliotta L,Martinelli G,Tura S,Baccarani M,Ferrari S,Catani L | Homo sapiens |
| KIM BIPOLAR DISORDER OLIGODENDROCYTE DENSITY CORR UP | Genes whose expression significantly and positively correlated with oligodendrocyte density in layer VI of BA9 brain region in patients with bipolar disorder. | 681 | 6 | 2.52E-05 | 0.010418 | NDUFS3:ALDH2:COX7A2L:NDUFA6:NDUFA5:GLDC | Pubmed 18762803 Authors: Kim S,Webster MJ | Homo sapiens |
| MOOTHA PGC | Genes up-regulated in differentiating C2C12 cells (myoblasts) upon expression of PPARGC1A off an adenoviral vector. | 421 | 5 | 3.17E-05 | 0.011622 | NDUFS3:COX8A:GSS:NDUFA6:NDUFA5 | Pubmed 12808457 Authors: Mootha VK,Lindgren CM,Eriksson KF,Subramanian A,Sihag S,Lehar J,Puigserver P,Carlsson E,Ridderstråle M,Laurila E,Houstis N,Daly MJ,Patterson N,Mesirov JP,Golub TR,Tamayo P,Spiegelman B,Lander ES,Hirschhorn JN,Altshuler D,Groop LC | Homo sapiens |
| DUTERTRE ESTRADIOL RESPONSE 6HR UP | Genes up-regulated in MCF7 cells (breast cancer) at 6 h of estradiol treatment | 228 | 4 | 4.73E-05 | 0.01464 | CTPS1:MYBBP1A:PAICS:JAK2 | Pubmed 20406972 Authors: Dutertre M,Gratadou L,Dardenne E,Germann S,Samaan S,Lidereau R,Driouch K,de la Grange P,Auboeuf D | Homo sapiens |
| SCHUHMACHER MYC TARGETS UP | Genes up-regulated in P493-6 cells (Burkitt's lymphoma) induced to express MYC | 80 | 3 | 4.88E-05 | 0.01464 | CTPS1:PPAT:PAICS | Pubmed 11139609 Authors: Schuhmacher M,Kohlhuber F,Hölzel M,Kaiser C,Burtscher H,Jarsch M,Bornkamm GW,Laux G,Polack A,Weidle UH,Eick D | Homo sapiens |
| WINTER HYPOXIA METAGENE | Genes regulated by hypoxia, based on literature searches | 240 | 4 | 5.78E-05 | 0.015892 | IGF2:KDR:PPAT:PAICS | Pubmed 17409455 Authors: Winter SC,Buffa FM,Silva P,Miller C,Valentine HR,Turley H,Shah KA,Cox GJ,Corbridge | Homo sapiens |

| | | | | | | | | |
|---|---|---|---|---|---|---|---|---|
| | | | | | | | RJ,Homer JJ,Musgrove B,Slevin N,Sloan P,Price P,West CM,Harris AL | |
| BLALOCK ALZHEIMERS DISEASE DN | Genes down-regulated in brain from patients with Alzheimer's disease. | 1244 | 7 | 8.54E-05 | 0.021699 | NDUFS3:COX8A:ALDH2:COX7A2L:ATP6V1G2:NDUFA5:JAK2 | Pubmed 14769913 Authors: Blalock EM,Geddes JW,Chen KC,Porter NM,Markesbery WR,Landfield PW | Homo sapiens |
| FLECHNER BIOPSY KIDNEY TRANSPLANT REJECTED VS OK DN | Genes down-regulated in kidney biopsies from patients with acute transplant rejection compared to the biopsies from patients with well functioning kidneys more than 1-year post transplant. | 552 | 5 | 0.000114 | 0.026804 | SDHC:ALDH2:KDR:NDUFA5:GLDC | Pubmed 15307835 Authors: Flechner SM,Kurian SM,Head SR,Sharp SM,Whisenant TC,Zhang J,Chismar JD,Horvath S,Mondala T,Gilmartin T,Cook DJ,Kay SA,Walker JR,Salomon DR | Homo sapiens |
| TIEN INTESTINE PROBIOTICS 24HR UP | Genes up-regulated in Caco-2 cells (intestinal epithelium) after coculture with the probiotic bacteria L. casei for 24h. | 560 | 5 | 0.000122 | 0.026804 | SDHC:NDUFA6:CCNB1:NDUFA5:GLDC | Pubmed 16394013 Authors: Tien MT,Girardin SE,Regnault B,Bourhis Le L,Dillies MA,Coppée JY,Bourdet-Sicard R,Sansonetti PJ,Pédron T | Homo sapiens |
| TARTE PLASMA CELL VS PLASMABLAST DN | Genes down-regulated in mature plasma cells compared with plasmablastic B lymphocytes. | 307 | 4 | 0.000149 | 0.030085 | CTPS1:PAICS:CCNB1:GLDC | Pubmed 12663452 Authors: Tarte K,Zhan F,De Vos J,Klein B,Shaughnessy J Jr | Homo sapiens |
| DAIRKEE CANCER PRONE RESPONSE BPA E2 | 'Cancer prone response profile' (CPRP): genes common to estradiol and bisphenol A [PubChem=5757;6623] response of epithelial cell cultures from patients at high risk of breast cancer. | 118 | 3 | 0.000155 | 0.030085 | CTPS1:NDUFS3:COX8A | Pubmed 18381411 Authors: Dairkee SH,Seok J,Champion S,Sayeed A,Mindrinos M,Xiao W,Davis RW,Goodson WH | Homo sapiens |
| CAIRO HEPATOBLASTOMA CLASSES UP | Genes up-regulated in robust Cluster 2 (rC2) of hepatoblastoma samples compared to those in the robust Cluster 1 (rC1). | 608 | 5 | 0.000179 | 0.032757 | MYBBP1A:PFAS:PPAT:CCNB1:GLDC | Pubmed 19061838 Authors: Cairo S,Armengol C,Reyniès De A,Wei Y,Thomas E,Renard CA,Goga A,Balakrishnan A,Semeraro M,Gresh L,Pontoglio M,Strick-Marchand H,Levillayer F,Nouet Y,Rickman D,Gauthier F,Branchereau S,Brugières | Homo sapiens |

| | | | | | | | | |
|---|---|---|---|---|---|---|---|---|
| | | | | | | | L,Laithier V,Bouvier R,Boman F,Basso G,Michiels JF,Hofman P,Arbez-Gindre F,Jouan H,Rousselet-Chapeau MC,Berrebi D,Marcellin L,Plenat F,Zachar D,Joubert M,Selves J,Pasquier D,Bioulac-Sage P,Grotzer M,Childs M,Fabre M,Buendia MA | |
| BASSO B LYMPHOCYTE NETWORK | Genes which comprise the top 1% of highly interconnected genes (major hubs) that account for most of gene interactions in the reconstructed regulatory networks from expression profiles in B lymphocytes. | 145 | 3 | 0.000284 | 0.049385 | CTPS1:PAICS:CCNB1 | Pubmed 15778709 Authors: Basso K,Margolin AA,Stolovitzky G,Klein U,Dalla-Favera R,Califano A | Homo sapiens |

Supplementary table 3: Chemical and Genetic pertubation database(MsigDB c2). N indicates the total number of genes in the set while n shows the number of genes that SNPs mapped to the 37 genes identified. Gene sets were queried using FUMA GWAS. The adjusted P is an FDR-corrected P-value based on the number of gene sets examined.

| Gene Set | Brief description | N | n | P-value | adj. P-value | Genes |
|---|---|---|---|---|---|---|
| RESPIRASOME | The protein complexes that form the electron transport system (the respiratory chain), associated with a cell membrane, usually the plasma membrane (in prokaryotes) or the inner mitochondrial membrane (on eukaryotes). The respiratory chain complexes transfer electrons from an electron donor to an electron acceptor and are associated with a proton pump to create a transmembrane electrochemical gradient. | 89 | 6 | 1.56E-10 | 1.56E-07 | SDHC:NDUFS3:COX8A:COX7A2L:NDUFA6:NDUFA5 |
| RESPIRATORY CHAIN COMPLEX | Any protein complex that is part of a respiratory chain. | 75 | 5 | 6.29E-09 | 3.15E-06 | SDHC:NDUFS3:COX8A:NDUFA6:NDUFA5 |
| MITOCHONDRIAL PART | The double lipid bilayer enclosing the mitochondrion and separating its contents from the cell cytoplasm; includes the intermembrane space. | 1019 | 10 | 1.03E-08 | 3.42E-06 | SDHC:NDUFS3:COX8A:ALDH2:POLG2:COX7A2L:NDUFA6:CCNB1:NDUFA5:GLDC |
| OXIDOREDUCTASE COMPLEX | Any protein complex that possesses oxidoreductase activity. | 106 | 5 | 3.62E-08 | 9.01E-06 | SDHC:NDUFS3:NDUFA6:NDUFA5:GLDC |
| MITOCHONDRION | A semiautonomous, self replicating organelle that occurs in varying numbers, shapes, and sizes in the cytoplasm of virtually all eukaryotic cells. It is notably the site of tissue respiration. | 1552 | 11 | 4.50E-08 | 9.01E-06 | SDHC:NDUFS3:COX8A:ALDH2:POLG2:GSK3A:COX7A2L:NDUFA6:CCNB1:NDUFA5:GLDC |
| MITOCHONDRIAL MEMBRANE PART | Either of the lipid bilayers that surround the mitochondrion and form the mitochondrial envelope. | 221 | 5 | 1.40E-06 | 0.000233 | SDHC:NDUFS3:COX7A2L:NDUFA6:NDUFA5 |
| INNER MITOCHONDRIAL MEMBRANE PROTEIN COMPLEX | Any protein complex that is part of the inner mitochondrial membrane. | 126 | 4 | 4.60E-06 | 0.000657 | SDHC:NDUFS3:NDUFA6:NDUFA5 |
| ORGANELLE INNER MEMBRANE | The inner, i.e. lumen-facing, lipid bilayer of an organelle envelope; usually highly selective to most ions and metabolites. | 516 | 6 | 5.25E-06 | 0.000657 | SDHC:NDUFS3:COX8A:COX7A2L:NDUFA6:NDUFA5 |
| MITOCHONDRIAL RESPIRATORY | The aggregation, arrangement and bonding together of a set of components to form mitochondrial respiratory chain complex I. | 46 | 3 | 9.20E-06 | 0.001023 | NDUFS3:NDUFA6:NDUFA5 |

| | | | | | | |
|---|---|---|---|---|---|---|
| CHAIN COMPLEX I | | | | | | |
| MITOCHONDRIAL ENVELOPE | The double lipid bilayer enclosing the mitochondrion and separating its contents from the cell cytoplasm; includes the intermembrane space. | 725 | 6 | 3.58E-05 | 0.003585 | SDHC:NDUFS3:COX8A:COX7A2L:NDUFA6:NDUFA5 |
| MITOCHONDRIAL MATRIX | The gel-like material, with considerable fine structure, that lies in the matrix space, or lumen, of a mitochondrion. It contains the enzymes of the tricarboxylic acid cycle and, in some organisms, the enzymes concerned with fatty acid oxidation. | 465 | 5 | 5.08E-05 | 0.00442 | NDUFS3:ALDH2:POLG2:CCNB1:GLDC |
| MEMBRANE PROTEIN COMPLEX | Any protein complex that is part of a membrane. | 1153 | 7 | 5.30E-05 | 0.00442 | SDHC:NDUFS3:COX8A:ACVR2A:NDUFA6:ATP6V1G2:NDUFA5 |
| MITOCHONDRIAL PROTEIN COMPLEX | A protein complex that is part of a mitochondrion. | 254 | 4 | 7.19E-05 | 0.005539 | SDHC:NDUFS3:NDUFA6:NDUFA5 |
| CATALYTIC COMPLEX | A protein complex which is capable of catalytic activity. [GOC:bhm, GOC:TermGenie, | 1351 | 7 | 0.000143 | 0.01021 | SDHC:NDUFS3:ACVR2A:NDUFA6:CCNB1:NDUFA5:GLDC |
| ENDOSOME LUMEN | The volume enclosed by the membrane of an endosome. | 34 | 2 | 0.000415 | 0.027703 | INS:JAK2 |
| ENVELOPE | A multilayered structure surrounding all or part of a cell; encompasses one or more lipid bilayers, and may include a cell wall layer; also includes the space between layers. | 1166 | 6 | 0.000476 | 0.029777 | SDHC:NDUFS3:COX8A:COX7A2L:NDUFA6:NDUFA5 |

Supplementary table 4: GO Cellular Component Database (MsigDB c5). N indicates the total number of genes in the set while n shows the number of genes that SNPs mapped to the 37 genes identified. Gene sets were queried using FUMA GWAS. The adjusted P is an FDR-corrected P-value based on the number of gene sets examined.

| Gene Set | Brief Description | N | n | P-value | adj. P-value | Genes | Publications | Species |
|---|---|---|---|---|---|---|---|---|
| OXIDATIVE PHOSPHORYLATION | Genes encoding proteins involved in oxidative phosphorylation. | 200 | 6 | 2.06E-08 | 1.03E-06 | SDHC:NDUFS3:COX8A:COX7A2L:NDUFA6:NDUFA5 | Pubmed 26771021 Authors: Liberzon A,Birger C,Thorvaldsdóttir H,Ghandi M,Mesirov JP,Tamayo P. | Homo Sapiens |
| ADIPOGENESIS | Genes up-regulated during adipocyte differentiation (adipogenesis). | 200 | 5 | 8.56E-07 | 2.14E-05 | SDHC:NDUFS3:COX8A:ALDH2:NDUFA5 | Pubmed 26771021 Authors: Liberzon A,Birger C,Thorvaldsdóttir H,Ghandi M,Mesirov JP,Tamayo P. | Homo Sapiens |
| XENOBIOTIC METABOLISM | Genes encoding proteins involved in processing of drugs and other xenobiotics. | 200 | 4 | 2.84E-05 | 0.000473 | PAPSS2:ALDH2:CYP1A2:GSS | Pubmed 26771021 Authors: Liberzon A,Birger C,Thorvaldsdóttir H,Ghandi M,Mesirov JP,Tamayo P. | Homo Sapiens |
| ESTROGEN RESPONSE EARLY | Genes defining early response to estrogen. | 200 | 3 | 0.000726 | 0.007258 | PAPSS2:MYBBP1A:JAK2 | Pubmed 26771021 Authors: Liberzon A,Birger C,Thorvaldsdóttir H,Ghandi M,Mesirov JP,Tamayo P. | Homo Sapiens |
| ALLOGRAFT REJECTION | Genes up-regulated during transplant rejection. | 200 | 3 | 0.000726 | 0.007258 | IL1B:ACVR2A:JAK2 | Pubmed 26771021 Authors: Liberzon A,Birger C,Thorvaldsdóttir H,Ghandi M,Mesirov JP,Tamayo P. | Homo Sapiens |

Supplementary table 5: Hallmark Gene Sets (MsigDB h). N indicates the total number of genes in the set while n shows the number of genes that SNPs mapped to the 37 genes identified. Gene sets were queried using FUMA GWAS. The adjusted P is an FDR-corrected P-value based on the number of gene sets examined.

| Gene Set | Brief description | N | n | P-value | adj. P-value | Genes | Publications | Species |
|---|---|---|---|---|---|---|---|---|
| GSE42724 B1 BCELL VS PLASMABLAST UP | Genes up-regulated in B lymphocytes: B1 versus plasmablasts. | 198 | 5 | 8.14E-07 | 0.002084 | ALDH2:GSK3A:COX7A2L:GSS:ATP6V1G2 | Pubmed 23613519 Authors: Covens K,Verbinnen B,Geukens N,Meyts I,Schuit F,Lommel Van L,Jacquemin M,Bossuyt X | Homo Sapiens |
| GSE36476 CTRL VS TSST ACT 40H MEMORY CD4 TCELL YOUNG DN | Genes down-regulated in comparison of untreated CD4 memory T cells from young donors versus those treated with TSST at 40 h | 200 | 5 | 8.56E-07 | 0.002084 | CTPS1:ALDH2:PFAS:PAICS:CCNB1 | Pubmed 22434910 Authors: Yu M,Li G,Lee WW,Yuan M,Cui D,Weyand CM,Goronzy JJ. | Homo Sapiens |
| GSE28130 ACTIVATED VS INDUCEED TREG DN | Genes down-regulated in activated versus induced T reg cells. | 195 | 4 | 2.57E-05 | 0.009883 | IGF2:GSS:NDUFA6:ATP6V1G2 | Pubmed 21642545 Authors: Kuczma M,Lee JR,Kraj P | Mus musculus |
| GSE3982 EOSINOPHIL VS BCELL DN | Genes down-regulated in activated versus induced T reg cells. | 197 | 4 | 2.68E-05 | 0.009883 | MYBBP1A:PFAS:COX7A2L:GLDC | Pubmed 16474395 Authors: Jeffrey KL,Brummer T,Rolph MS,Liu SM,Callejas NA,Grumont RJ,Gillieron C,Mackay F,Grey S,Camps M,Rommel C,Gerondakis SD,Mackay CR. | Homo Sapiens |
| GSE21670 UNTREATED VS IL6 TREATED STAT3 KO CD4 TCELL DN | Genes down-regulated in CD4 T cells with STAT3 knockout medium versus IL6. | 197 | 4 | 2.68E-05 | 0.009883 | CTPS1:PFAS:IL1B:PAICS | Pubmed 20493732 Authors: Durant L,Watford WT,Ramos HL,Laurence A,Vahedi G,Wei L,Takahashi H,Sun HW,Kanno Y,Powrie F,O'Shea JJ | Homo Sapiens |
| GSE17721 LPS VS PAM3CSK4 12H BMDC DN | Genes down-regulated in comparison of dendritic cells (DC) stimulated with LPS (TLR4 agonist) at 12 h versus DC cells stimulated with Pam3Csk4 (TLR1/2 agonist) at 12 h. | 198 | 4 | 2.73E-05 | 0.009883 | SDHC:COX8A:IL1B:NDUFA6 | Pubmed 19729616 Authors: Amit I,Garber M,Chevrier N,Leite AP,Donner Y,Eisenhaure T,Guttman M,Grenier JK,Li W,Zuk O,Schubert LA,Birditt B,Shay T,Goren A,Zhang X,Smith Z,Deering R,McDonald RC,Cabili M,Bernstein BE,Rinn JL,Meissner A,Root DE,Hacohen N,Regev A. | Homo Sapiens |

| | | | | | | | | |
|---|---|---|---|---|---|---|---|---|
| GSE36476 CTRL VS TSST ACT 40H MEMORY CD4 TCELL YOUNG DN | Genes down-regulated in comparison of untreated CD4 memory T cells from young donors versus those treated with TSST at 40 h. | 199 | 4 | 2.78E-05 | 0.009883 | CTPS1:PFAS:PAICS:CCNB1 | Pubmed 22434910 Authors: Yu M,Li G,Lee WW,Yuan M,Cui D,Weyand CM,Goronzy JJ. | Homo Sapiens |
| GSE28726 NAIVE VS ACTIVATED CD4 TCELL DN | Genes down-regulated in CD4 T cells naïve versus activated. | 199 | 4 | 2.78E-05 | 0.009883 | CTPS1:COX8A:PAICS:CCNB1 | Pubmed 21632718 Authors: Constantinides MG,Picard D,Savage AK,Bendelac A | Homo Sapiens |
| GSE8685 IL2 STARVED VS IL15 ACT IL2 STARVED CD4 TCELL DN | Genes down-regulated in Sez-2 cells (T cell lymphoma): untreated versus IL5. | 200 | 4 | 2.84E-05 | 0.009883 | CTPS1:PFAS:GSK3A:GSS | Pubmed 18281483 Authors: Marzec M,Halasa K,Kasprzycka M,Wysocka M,Liu X,Tobias JW,Baldwin D,Zhang Q,Odum N,Rook AH,Wasik MA | Homo Sapiens |
| GSE36476 CTRL VS TSST ACT 40H MEMORY CD4 TCELL OLD DN | Genes down-regulated in comparison of untreated CD4 memory T cells from old donors versus those treated with TSST at 40 h. | 200 | 4 | 2.84E-05 | 0.009883 | CTPS1:ALDH2:PAICS:CCNB1 | Pubmed 22434910 Authors: Yu M,Li G,Lee WW,Yuan M,Cui D,Weyand CM,Goronzy JJ. | Homo Sapiens |
| GSE36476 CTRL VS TSST ACT 72H MEMORY CD4 TCELL OLD DN | Genes down-regulated in comparison of untreated CD4 memory T cells from old donors versus those treated with TSST at 72 h. | 200 | 4 | 2.84E-05 | 0.009883 | CTPS1:NDUFA6:PAICS:CCNB1 | Pubmed 22434910 Authors: Yu M,Li G,Lee WW,Yuan M,Cui D,Weyand CM,Goronzy JJ. | Homo Sapiens |
| GSE22886 UNSTIM VS IL15 STIM NKCELL DN | Genes down-regulated in comparison of unstimulated NK cells versus those stimulated with IL2 at 16 h. | 200 | 4 | 2.84E-05 | 0.009883 | CTPS1:NDUFA6:PPAT:PAICS | Pubmed 15789058 Authors: Abbas AR,Baldwin D,Ma Y,Ouyang W,Gurney A,Martin F,Fong S,van Lookeren Campagne M,Godowski P,Williams PM,Chan AC,Clark HF. | Homo Sapiens |
| GSE36476 CTRL VS TSST ACT 16H MEMORY CD4 TCELL YOUNG DN | Genes down-regulated in comparison of untreated CD4 memory T cells from young donors versus those treated with TSST at 16 h. | 200 | 4 | 2.84E-05 | 0.009883 | CTPS1:ALDH2:PFAS:PAICS | Pubmed 22434910 Authors: Yu M,Li G,Lee WW,Yuan M,Cui D,Weyand CM,Goronzy JJ. | Homo Sapiens |

| | | | | | | | | |
|---|---|---|---|---|---|---|---|---|
| GSE33292 WT VS TCF1 KO DN3 THYMOCYTE DN | Genes down-regulated in DN3 thymocytes wildtype versus TCF7 knockout. | 200 | 4 | 2.84E-05 | 0.009883 | GSS:PAICS:CCNB1:GLDC | Pubmed 23103132 Authors: Yu S,Zhou X,Steinke FC,Liu C,Chen SC,Zagorodna O,Jing X,Yokota Y,Meyerholz DK,Mullighan CG,Knudson CM,Zhao DM,Xue HH | Mus musculus |
| GSE3565 CTRL VS LPS INJECTED SPLENOCYTES DN | Genes down-regulated in spleen from wildtype mice: control versus LPS. | 189 | 3 | 0.000616 | 0.046527 | CTPS1:MYBBP1A:COX7A2L | Pubmed 16380512 Authors: Hammer M,Mages J,Dietrich H,Servatius A,Howells N,Cato AC,Lang R | Mus musculus |
| GSE29614 CTRL VS DAY7 TIV FLU VACCINE PBMC DN | Genes down-regulated in comparison of peripheral blood mononuclear cells (PBMC) from TIV influenza vaccinee pre-vaccination versus those from day 7 post-vaccination. | 192 | 3 | 0.000645 | 0.046527 | INS:CCNB1:GLDC | Pubmed 21743478 Authors: Nakaya HI,Wrammert J,Lee EK,Racioppi L,Marie-Kunze S,Haining WN,Means AR,Kasturi SP,Khan N,Li GM,McCausland M,Kanchan V,Kokko KE,Li S,Elbein R,Mehta AK,Aderem A,Subbarao K,Ahmed R,Pulendran B. | Homo Sapiens |
| GSE25846 IL10 POS VS NEG CD8 TCELL DAY7 POST CORONAVIRUS BRAIN DN | Genes down-regulated in CD8 T cells IL10+ versus IL10-. | 193 | 3 | 0.000655 | 0.046527 | NDUFS3:ALDH2:ATP6V1G2 | Pubmed 21317392 Authors: Trandem K,Zhao J,Fleming E,Perlman S | Mus musculus |
| GSE25087 TREG VS TCONV FETUS DN | Genes down-regulated in comparison of fetal regulatory T cell (Treg) versus fetal conventional T cells. | 193 | 3 | 0.000655 | 0.046527 | POLG2:ACVR2A:PAICS | Pubmed 21164017 Authors: Mold JE,Venkatasubrahmanyam S,Burt TD,Michaëlsson J,Rivera JM,Galkina SA,Weinberg K,Stoddart CA,McCune JM | Homo Sapiens |
| GSE19772 CTRL VS HCMV INF MONOCYTES AND PI3K INHIBITION UP | Genes up-regulated in monocytes pre-treated with Ly294002 control versus HCMV infection. | 193 | 3 | 0.000655 | 0.046527 | SDHC:MYBBP1A:GSK3A | Pubmed 20173022 Authors: Chan G,Nogalski MT,Bentz GL,Smith MS,Parmater A,Yurochko AD | Homo Sapiens |
| GSE27859 MACROPHAGE VS CD11C INT F480 INT DC DN | Genes down-regulated in macrophages versus dendritic cells sorted as ITGAX int and EMR1 int. | 195 | 3 | 0.000674 | 0.046527 | CTPS1:NDUFA6:DDX39B | Pubmed 22231304 Authors: Rivollier A,He J,Kole A,Valatas V,Kelsall BL | Mus musculus |

| | | | | | | | | |
|---|---|---|---|---|---|---|---|---|
| GSE19512 NAUTRAL VS INDUCED TREG UP | Genes up-regulated in T reg: natural versus induced cells. | 196 | 3 | 0.000685 | 0.046527 | IGF2:GSK3A:ATP6V1G2 | Pubmed 21723159 Authors: Haribhai D,Williams JB,Jia S,Nickerson D,Schmitt EG,Edwards B,Ziegelbauer J,Yassai M,Li SH,Relland LM,Wise PM,Chen A,Zheng YQ,Simpson PM,Gorski J,Salzman NH,Hessner MJ,Chatila TA,Williams CB | Mus musculus |
| GSE9988 ANTI TREM1 VS LPS MONOCYTE DN | Genes down-regulated in comparison of monocytes treated with anti-TREM1 versus monocytes treated with 5000 ng/ml LPS (TLR4 agonist). | 196 | 3 | 0.000685 | 0.046527 | IL1B:ACVR2A:CCNB1 | Pubmed 18292579 Authors: Dower K,Ellis DK,Saraf K,Jelinsky SA,Lin LL. | Homo Sapiens |
| GSE12845 IGD NEG BLOOD VS PRE GC TONSIL BCELL DN | Genes down-regulated in comparison of IgD- peripheral blood B cells versus pre-germinal center B cells. | 196 | 3 | 0.000685 | 0.046527 | ALDH2:PFAS:CCNB1 | Pubmed 19023113 Authors: Longo NS,Lugar PL,Yavuz S,Zhang W,Krijger PH,Russ DE,Jima DD,Dave SS,Grammer AC,Lipsky PE. | Homo Sapiens |
| GSE1460 CORD VS ADULT BLOOD NAIVE CD4 TCELL UP | Genes up-regulated in CD4 T cells from cord blood versus those from adult blood. | 196 | 3 | 0.000685 | 0.046527 | SDHC:MYBBP1A:PAICS | Pubmed 15210650 Authors: Lee MS,Hanspers K,Barker CS,Korn AP,McCune JM. | Homo Sapiens |
| GSE360 CTRL VS B MALAYI LOW DOSE MAC DN | Genes down-regulated in comparison of macrophages versus macrophages exposed to B. malayi (5 worms/well). | 197 | 3 | 0.000695 | 0.046527 | GSK3A:NDUFA6:PAICS | Pubmed 12663451 Authors: Chaussabel D,Semnani RT,McDowell MA,Sacks D,Sher A,Nutman TB. | Homo Sapiens |
| GSE36476 YOUNG VS OLD DONOR MEMORY CD4 TCELL 40H TSST ACT DN | Genes down-regulated in comparison of memory CD4 T cells from young donors treated with TSST at 40 h versus those from old donors treated with TSST at 40 h. | 197 | 3 | 0.000695 | 0.046527 | NDUFS3:COX8A:ALDH2 | Pubmed 22434910 Authors: Yu M,Li G,Lee WW,Yuan M,Cui D,Weyand CM,Goronzy JJ. | Homo Sapiens |
| GSE45365 WT VS IFNAR KO CD8A DC MCMV INFECTION UP | Genes up-regulated during primary acute viral infection in | 198 | 3 | 0.000705 | 0.046527 | COX8A:DDX39B:JAK2 | | Mus musculus |

| | | | | | | | | |
|---|---|---|---|---|---|---|---|---|
| | CD8A dendritic cells wildtype versus IFNAR1 knockout. | | | | | | | |
| GSE43955 1H VS 42H ACT CD4 TCELL WITH TGFB IL6 UP | Genes up-regulated in CD4 T helper cells Th17 treated with TGFB1 and IL6 1h versus 42h. | 198 | 3 | 0.000705 | 0.046527 | SDHC:ALDH2:COX7A2L | Pubmed 23467089 Authors: Yosef N,Shalek AK,Gaublomme JT,Jin H,Lee Y,Awasthi A,Wu C,Karwacz K,Xiao S,Jorgolli M,Gennert D,Satija R,Shakya A,Lu DY,Trombetta JJ,Pillai MR,Ratcliffe PJ,Coleman ML,Bix M,Tantin D,Park H,Kuchroo VK,Regev A | Mus musculus |
| GSE27434 WT VS DNMT1 KO TREG DN | Genes down-regulated in T reg wildtype versus DNMT1 knockout. | 198 | 3 | 0.000705 | 0.046527 | CTPS1:IL1B:JAK2 | Pubmed 23444399 Authors: Wang L,Liu Y,Beier UH,Han R,Bhatti TR,Akimova T,Hancock WW | Mus musculus |
| GSE42021 TCONV PLN VS TREG PRECURSORS THYMUS DN | Genes down-regulated in T conv from: peripheral lymph nodes versus thymic precursors. | 198 | 3 | 0.000705 | 0.046527 | CTPS1:IL1B:JAK2 | Pubmed 23420886 Authors: Toker A,Engelbert D,Garg G,Polansky JK,Floess S,Miyao T,Baron U,Düber S,Geffers R,Giehr P,Schallenberg S,Kretschmer K,Olek S,Walter J,Weiss S,Hori S,Hamann A,Huehn J | Mus musculus |
| GSE3982 NEUTROPHIL VS TH2 DN | Genes down-regulated in comparison of neutrophils versus Th2 cells. | 198 | 3 | 0.000705 | 0.046527 | PFAS:DDX39B:NDUFA5 | Pubmed 16474395 Authors: Jeffrey KL,Brummer T,Rolph MS,Liu SM,Callejas NA,Grumont RJ,Gillieron C,Mackay F,Grey S,Camps M,Rommel C,Gerondakis SD,Mackay CR. | Homo Sapiens |
| GSE42021 TREG PLN VS CD24INT TREG THYMUS UP | Genes up-regulated in T reg peripheral lymph nodes versus thymic CD24 int. | 198 | 3 | 0.000705 | 0.046527 | CTPS1:IL1B:JAK2 | Pubmed 23420886 Authors: Toker A,Engelbert D,Garg G,Polansky JK,Floess S,Miyao T,Baron U,Düber S,Geffers R,Giehr P,Schallenberg S,Kretschmer K,Olek S,Walter J,Weiss S,Hori S,Hamann A,Huehn J | Mus musculus |

| | | | | | | | | |
|---|---|---|---|---|---|---|---|---|
| GSE42021 TREG PLN VS CD24HI TREG THYMUS UP | Genes up-regulated in T reg: peripheral lymph nodes versus thymic CD24 high | 198 | 3 | 0.000705 | 0.046527 | CTPS1:IGF2:IL1B | Pubmed 23420886 Authors: Toker A,Engelbert D,Garg G,Polansky JK,Floess S,Miyao T,Baron U,Düber S,Geffers R,Giehr P,Schallenberg S,Kretschmer K,Olek S,Walter J,Weiss S,Hori S,Hamann A,Huehn J | Mus musculus |
| GSE17721 PAM3CSK4 VS GADIQUIMOD 8H BMDC UP | Genes up-regulated in comparison of dendritic cells (DC) stimulated with Pam3Csk4 (TLR1/2 agonist) at 8 h versus DC cells stimulated with Gardiquimod (TLR7 agonist) at 8 h. | 198 | 3 | 0.000705 | 0.046527 | GSS:PAICS:NDUFA5 | Pubmed 19729616 Authors: Amit I,Garber M,Chevrier N,Leite AP,Donner Y,Eisenhaure T,Guttman M,Grenier JK,Li W,Zuk O,Schubert LA,Birditt B,Shay T,Goren A,Zhang X,Smith Z,Deering R,McDonald RC,Cabili M,Bernstein BE,Rinn JL,Meissner A,Root DE,Hacohen N,Regev A. | Mus musculus |
| GSE24574 BCL6 HIGH TFH VS NAIVE CD4 TCELL UP | Genes up-regulated in BCL6 high follicular helper T cells versus naïve T CD4 cells. | 198 | 3 | 0.000705 | 0.046527 | PAPSS2:INS:ALDH2 | Pubmed 21636294 Authors: Kitano M,Moriyama S,Ando Y,Hikida M,Mori Y,Kurosaki T,Okada T | Mus musculus |
| GSE28726 NAIVE CD4 TCELL VS NAIVE NKTCELL UP | Genes up-regulated in naïve T cells: CD4 versus NK. | 199 | 3 | 0.000715 | 0.046527 | CTPS1:PFAS:PAICS | Pubmed 21632718 Authors: Constantinides MG,Picard D,Savage AK,Bendelac A | Homo Sapiens |
| GSE28726 NAIVE VS ACTIVATED NKTCELL UP | Genes up-regulated in NKT cells: naïve versus activated. | 199 | 3 | 0.000715 | 0.046527 | PAPSS2:INS:ALDH2 | Pubmed 21632718 Authors: Constantinides MG,Picard D,Savage AK,Bendelac A | Homo Sapiens |
| GSE39556 CD8A DC VS NK CELL UP | Genes up-regulated in CD8A dendritic versus NK cells. | 199 | 3 | 0.000715 | 0.046527 | GSS:PAICS:GLDC | Pubmed 23084923 Authors: Baranek T,Manh TP,Alexandre Y,Maqbool MA,Cabeza JZ,Tomasello E,Crozat K,Bessou G,Zucchini N,Robbins SH,Vivier E,Kalinke U,Ferrier P,Dalod M | Mus musculus |
| GSE32986 CURDLAN LOWDOSE VS GMCSF AND | Genes down-regulated in bone marrow-derived dendritic cells low dose of 1,3-beta-D- | 199 | 3 | 0.000715 | 0.046527 | CTPS1:IL1B:JAK2 | Pubmed 22250091 Authors: Min L,Isa SA,Fam WN,Sze SK,Beretta O,Mortellaro A,Ruedl C | Mus musculus |

| | | | | | | | | |
|---|---|---|---|---|---|---|---|---|
| CURDLAN LOWDOSE STIM DC DN | oligoglucan versus CSF2 and low dose of 1,3-beta-D-oligoglucan. | | | | | | | |
| GSE23114 PERITONEAL CAVITY B1A BCELL VS SPLEEN BCELL IN SLE2C1 MOUSE UP | Genes up-regulated in lupus susceptibility locus Sle2c1 B lymphocytes from: peritoneal cavity versus spleen. | 199 | 3 | 0.000715 | 0.046527 | PAPSS2:INS:ALDH2 | Pubmed 21543644 Authors: Xu Z,Potula HH,Vallurupalli A,Perry D,Baker H,Croker BP,Dozmorov I,Morel L | Mus musculus |
| GSE12845 IGD NEG BLOOD VS DARKZONE GC TONSIL BCELL DN | Genes down-regulated in comparison of IgD- peripheral blood B cells versus dark zone germinal center B cells. | 199 | 3 | 0.000715 | 0.046527 | COX8A:PFAS:PPAT | Pubmed 19023113 Authors: Longo NS,Lugar PL,Yavuz S,Zhang W,Krijger PH,Russ DE,Jima DD,Dave SS,Grammer AC,Lipsky PE. | Homo Sapiens |
| GSE43955 1H VS 10H ACT CD4 TCELL WITH TGFB IL6 UP | Genes up-regulated in CD4 T helper cells Th17 treated with TGFB1 and IL6 | 199 | 3 | 0.000715 | 0.046527 | NDUFS3:ALDH2:COX7A2L | Pubmed 23467089 Authors: Yosef N,Shalek AK,Gaublomme JT,Jin H,Lee Y,Awasthi A,Wu C,Karwacz K,Xiao S,Jorgolli M,Gennert D,Satija R,Shakya A,Lu DY,Trombetta JJ,Pillai MR,Ratcliffe PJ,Coleman ML,Bix M,Tantin D,Park H,Kuchroo VK,Regev A | Mus musculus |
| GSE32986 UNSTIM VS GMCSF AND CURDLAN HIGHDOSE STIM DC UP | Genes up-regulated in bone marrow-derived dendritic cells unstimulated versus CSF2 and high dose of 1,3-beta-D-oligoglucan. | 199 | 3 | 0.000715 | 0.046527 | CTPS1:COX8A:JAK2 | Pubmed 22250091 Authors: Min L,Isa SA,Fam WN,Sze SK,Beretta O,Mortellaro A,Ruedl C | Mus musculus |
| GSE22432 CONVENTIONAL CDC VS PLASMACYTOID PDC UP | Genes up-regulated in dendritic cells: common versus plasmacytoid. | 199 | 3 | 0.000715 | 0.046527 | CTPS1:PPAT:DDX39B | Pubmed 20881193 Authors: Felker P,Seré K,Lin Q,Becker C,Hristov M,Hieronymus T,Zenke M | Mus musculus |
| GSE3337 CTRL VS 4H IFNG IN CD8POS DC DN | Genes down-regulated in comparison of untreated CD8+ dendritic cells (DC) at 4 h versus those treated with IFNG at 4 h. | 199 | 3 | 0.000715 | 0.046527 | NDUFA6:KDR:PPAT | Pubmed 16339401 Authors: Orabona C,Puccetti P,Vacca C,Bicciato S,Luchini A,Fallarino F,Bianchi R,Velardi E,Perruccio | Mus musculus |

| | | | | | | | | |
|---|---|---|---|---|---|---|---|---|
| | | | | | | | K,Velardi A,Bronte V,Fioretti MC,Grohmann U. | |
| GSE26156 DOUBLE POSITIVE VS CD4 SINGLE POSITIVE THYMOCYTE DN | Genes down-regulated in thymocytes double positive versus CD4 single positive. | 199 | 3 | 0.000715 | 0.046527 | CTPS1:PAICS:CCNB1 | Pubmed 21551231 Authors: Ghisi M,Corradin A,Basso K,Frasson C,Serafin V,Mukherjee S,Mussolin L,Ruggero K,Bonanno L,Guffanti A,Bellis De G,Gerosa G,Stellin G,D'Agostino DM,Basso G,Bronte V,Indraccolo S,Amadori A,Zanovello P | Homo Sapiens |
| GSE18791 CTRL VS NEWCASTLE VIRUS DC 6H UP | Genes up-regulated in comparison of control conventional dendritic cells (cDC) at 0 h versus cDCs infected with Newcastle disease virus (NDV) at 6 h. | 199 | 3 | 0.000715 | 0.046527 | CTPS1:PPAT:NDUFA5 | Pubmed 20164420 Authors: Zaslavsky E,Hershberg U,Seto J,Pham AM,Marquez S,Duke JL,Wetmur JG,Tenoever BR,Sealfon SC,Kleinstein SH. | Homo Sapiens |
| GSE3982 EFF MEMORY CD4 TCELL VS TH1 DN | Genes down-regulated in comparison of effective memory CD4 T cells versus Th1 cells. | 199 | 3 | 0.000715 | 0.046527 | SDHC:NDUFA6:PAICS | Pubmed 16474395 Authors: Jeffrey KL,Brummer T,Rolph MS,Liu SM,Callejas NA,Grumont RJ,Gillieron C,Mackay F,Grey S,Camps M,Rommel C,Gerondakis SD,Mackay CR. | Homo Sapiens |
| GSE21670 STAT3 KO VS WT CD4 TCELL DN | Genes down-regulated in CD4 T cells STAT3 knockout versus wildtype. | 199 | 3 | 0.000715 | 0.046527 | CTPS1:IL1B:PAICS | Pubmed 20493732 Authors: Durant L,Watford WT,Ramos HL,Laurence A,Vahedi G,Wei L,Takahashi H,Sun HW,Kanno Y,Powrie F,O'Shea JJ | Mus musculus |
| GSE5503 MLN DC VS PLN DC ACTIVATED ALLOGENIC TCELL UP | Genes up-regulated in allogeneic T cells after stimulation with dendritic cells from lymph nodes: mesenteric (mLN) versus peripheral (pLN). | 199 | 3 | 0.000715 | 0.046527 | SDHC:COX8A:DDX39B | Pubmed 18178870 Authors: Kim TD,Terwey TH,Zakrzewski JL,Suh D,Kochman AA,Chen ME,King CG,Borsotti C,Grubin J,Smith OM,Heller G,Liu C,Murphy GF,Alpdogan O,Brink den van MR | Mus musculus |

| | | | | | | | | |
|---|---|---|---|---|---|---|---|---|
| GSE21927 BALBC VS C57BL6 MONOCYTE TUMOR UP | Genes up-regulated in CD11b Tumor from BALBc mouse versus CD11b Tumor from C57BL6 mouse. | 199 | 3 | 0.000715 | 0.046527 | COX8A:PFAS:PPAT | Pubmed 20605485 Authors: Marigo I,Bosio E,Solito S,Mesa C,Fernandez A,Dolcetti L,Ugel S,Sonda N,Bicciato S,Falisi E,Calabrese F,Basso G,Zanovello P,Cozzi E,Mandruzzato S,Bronte V | Mus musculus |
| GSE27786 NKTCELL VS MONO MAC UP | Genes up-regulated in comparison of NKT cells versus monocyte macrophages. | 200 | 3 | 0.000726 | 0.046527 | NDUFS3:PFAS:DDX39B | Pubmed 21540074 Authors: Konuma T,Nakamura S,Miyagi S,Negishi M,Chiba T,Oguro H,Yuan J,Mochizuki-Kashio M,Ichikawa H,Miyoshi H,Vidal M,Iwama A. | Mus musculus |
| GSE30083 SP1 VS SP2 THYMOCYTE UP | Genes up-regulated in comparison of SP1 thymocytes versus SP2 thymocytes. | 200 | 3 | 0.000726 | 0.046527 | ALDH2:GSS:ATP6V1G2 | Pubmed 22022412 Authors: Teng F,Zhou Y,Jin R,Chen Y,Pei X,Liu Y,Dong J,Wang W,Pang X,Qian X,Chen WF,Zhang Y,Ge Q. | Mus musculus |
| GSE27434 WT VS DNMT1 KO TREG UP | Genes up-regulated in T reg: wildtype versus DNMT1 knockout. | 200 | 3 | 0.000726 | 0.046527 | SDHC:NDUFS3:DDX39B | Pubmed 23444399 Authors: Wang L,Liu Y,Beier UH,Han R,Bhatti TR,Akimova T,Hancock WW | Mus musculus |
| GSE5679 CTRL VS PPARG LIGAND ROSIGLITAZONE TREATED DC UP | Genes up-regulated in monocyte-derived dendritic cells: untreated versus rosiglitazone | 200 | 3 | 0.000726 | 0.046527 | CTPS1:SDHC:PAICS | Pubmed 16982809 Authors: Szatmari I,Pap A,Rühl R,Ma JX,Illarionov PA,Besra GS,Rajnavolgyi E,Dezso B,Nagy L | Homo Sapiens |
| GSE36476 CTRL VS TSST ACT 16H MEMORY CD4 TCELL OLD DN | Genes down-regulated in comparison of untreated CD4 memory T cells from old donors versus those treated with TSST at 16 h. | 200 | 3 | 0.000726 | 0.046527 | CTPS1:ALDH2:PAICS | Pubmed 22434910 Authors: Yu M,Li G,Lee WW,Yuan M,Cui D,Weyand CM,Goronzy JJ. | Homo Sapiens |
| GSE22886 UNSTIM VS IL2 STIM NKCELL DN | Genes down-regulated in comparison of unstimulated NK cells versus those stimulated with IL2 at 16 h. | 200 | 3 | 0.000726 | 0.046527 | CTPS1:NDUFA6:PAICS | Pubmed 15789058 Authors: Abbas AR,Baldwin D,Ma Y,Ouyang W,Gurney A,Martin F,Fong S,van Lookeren Campagne M,Godowski P,Williams PM,Chan AC,Clark HF. | Homo Sapiens |

| | | | | | | | | |
|---|---|---|---|---|---|---|---|---|
| GSE42021 CD24LO TREG VS CD24LO TCONV THYMUS DN | Genes down-regulated in CD42 low cells from thymus T reg versus T conv. | 200 | 3 | 0.000726 | 0.046527 | SDHC:NDUFS3:DDX39B | Pubmed 23420886 Authors: Toker A,Engelbert D,Garg G,Polansky JK,Floess S,Miyao T,Baron U,Düber S,Geffers R,Giehr P,Schallenberg S,Kretschmer K,Olek S,Walter J,Weiss S,Hori S,Hamann A,Huehn J | Mus musculus |
| GSE22886 NAIVE CD4 TCELL VS 48H ACT TH1 DN | Genes down-regulated in comparison of naive CD4 T cells versus stimulated CD4 Th1 cells at 48 h. | 200 | 3 | 0.000726 | 0.046527 | CTPS1:NDUFS3:COX8A | Pubmed 15789058 Authors: Abbas AR,Baldwin D,Ma Y,Ouyang W,Gurney A,Martin F,Fong S,van Lookeren Campagne M,Godowski P,Williams PM,Chan AC,Clark HF. | Homo Sapiens |
| GSE43863 TH1 VS LY6C LOW CXCR5NEG EFFECTOR CD4 TCELL DN | Genes down-regulated in CD4 SMARTA effector T cells during acute infection of LCMV | 200 | 3 | 0.000726 | 0.046527 | NDUFS3:COX8A:COX7A2L | Pubmed 23583644 Authors: Hale JS,Youngblood B,Latner DR,Mohammed AU,Ye L,Akondy RS,Wu T,Iyer SS,Ahmed R | Mus musculus |
| GSE4142 GC BCELL VS MEMORY BCELL DN | Genes down-regulated in B lymphocytes: germinal center versus memory. | 200 | 3 | 0.000726 | 0.046527 | SDHC:COX8A:ACVR2A | Pubmed 16492737 Authors: Luckey CJ,Bhattacharya D,Goldrath AW,Weissman IL,Benoist C,Mathis D | Mus musculus |
| GSE24634 TREG VS TCONV POST DAY5 IL4 CONVERSION UP | Genes up-regulated in comparison of CD25+ T cells treated with IL4 versus CD25- T cells treated with IL4 at day 5. | 200 | 3 | 0.000726 | 0.046527 | PFAS:PPAT:PAICS | Pubmed 21347372 Authors: Prots I,Skapenko A,Lipsky PE,Schulze-Koops H. | Homo Sapiens |
| GSE16385 MONOCYTE VS 12H IL4 TREATED MACROPHAGE DN | Genes down-regulated in monocytes (12h) versus macrophages (12h) treated with IL4 | 200 | 3 | 0.000726 | 0.046527 | ALDH2:DDX39B:NFKBIL1 | Pubmed 21093321 Authors: Szanto A,Balint BL,Nagy ZS,Barta E,Dezso B,Pap A,Szeles L,Poliska S,Oros M,Evans RM,Barak Y,Schwabe J,Nagy L | Homo Sapiens |
| GSE26030 TH1 VS TH17 RESTIMULATED DAY5 POST POLARIZATION UP | Genes up-regulated in T helper cells 5 days post polarization and stimulated with anti-CD3 and anti-CD28: Th1 versus Th17. | 200 | 3 | 0.000726 | 0.046527 | NDUFS3:COX8A:NDUFA6 | Pubmed 22177921 Authors: Muranski P,Borman ZA,Kerkar SP,Klebanoff CA,Ji Y,Sanchez-Perez L,Sukumar M,Reger RN,Yu Z,Kern SJ,Roychoudhuri | Mus musculus |

| | | | | | | | | |
|---|---|---|---|---|---|---|---|---|
| | | | | | | | R,Ferreyra GA,Shen W,Durum SK,Feigenbaum L,Palmer DC,Antony PA,Chan CC,Laurence A,Danner RL,Gattinoni L,Restifo NP | |
| GSE1460 NAIVE CD4 TCELL ADULT BLOOD VS THYMIC STROMAL CELL DN | Genes down-regulated in naive CD4 T cells from adult blood versus thymic stromal cells. | 200 | 3 | 0.000726 | 0.046527 | COX8A:ALDH2:NDUFA6 | Pubmed 15210650 Authors: Lee MS,Hanspers K,Barker CS,Korn AP,McCune JM. | Homo Sapiens |
| GSE17721 0.5H VS 24H PAM3CSK4 BMDC UP | Genes up-regulated in comparison of dendritic cells (DC) stimulated with Pam3Csk4 (TLR1/2 agonist) at 0.5 h versus those stimulated at 24 h. | 200 | 3 | 0.000726 | 0.046527 | KDR:DDX39B:NDUFA5 | Pubmed 19729616 Authors: Amit I,Garber M,Chevrier N,Leite AP,Donner Y,Eisenhaure T,Guttman M,Grenier JK,Li W,Zuk O,Schubert LA,Birditt B,Shay T,Goren A,Zhang X,Smith Z,Deering R,McDonald RC,Cabili M,Bernstein BE,Rinn JL,Meissner A,Root DE,Hacohen N,Regev A. | Mus musculus |
| GSE24574 BCL6 HIGH VS LOW TFH CD4 TCELL DN | Genes down-regulated in follicular helper T cells: BCL6 high versus BCL6 low. | 200 | 3 | 0.000726 | 0.046527 | COX8A:PFAS:CCNB1 | Pubmed 21636294 Authors: Kitano M,Moriyama S,Ando Y,Hikida M,Mori Y,Kurosaki T,Okada T | Mus musculus |
| GSE16451 CTRL VS WEST EQUINE ENC VIRUS MATURE NEURON CELL LINE UP | Genes up-regulated in the mature neuron cell line: control versus infected with western equine encephalitis virus. | 200 | 3 | 0.000726 | 0.046527 | PFAS:GSS:JAK2 | Pubmed 20483728 Authors: Peltier DC,Simms A,Farmer JR,Miller DJ | Homo Sapiens |
| GSE15930 NAIVE VS 72H IN VITRO STIM CD8 TCELL UP | Genes up-regulated in comparison of CD8 T cells at 0 h versus those at 72 h. | 200 | 3 | 0.000726 | 0.046527 | PAPSS2:POLG2:ACVR2A | Pubmed 19592655 Authors: Agarwal P,Raghavan A,Nandiwada SL,Curtsinger JM,Bohjanen PR,Mueller DL,Mescher MF. | Mus musculus |
| GSE28726 NAIVE CD4 TCELL VS NAIVE NKTCELL DN | Genes down-regulated in naïve T cells: CD4 versus NK. | 200 | 3 | 0.000726 | 0.046527 | PAPSS2:INS:ALDH2 | Pubmed 21632718 Authors: Constantinides MG,Picard D,Savage AK,Bendelac A | Homo Sapiens |
| GSE28726 NAIVE CD4 TCELL VS NAIVE VA24NEG NKTCELL UP | Genes up-regulated in naïve T cells: CD4 versus Va24- NKT. | 200 | 3 | 0.000726 | 0.046527 | COX8A:PAICS:CCNB1 | Pubmed 21632718 Authors: Constantinides MG,Picard D,Savage AK,Bendelac A | Homo Sapiens |

| | | | | | | | | |
|---|---|---|---|---|---|---|---|---|
| GSE21927 GMCSF IL6 VS GMCSF GCSF TREATED BONE MARROW DN | Genes down-regulated in CD11b BoneMarrow from BALBc mouse incubated with GMCSF and IL-6 versus CD11b BoneMarrow from BALBc mouse incubated with GMCSF and GCSF. | 200 | 3 | 0.000726 | 0.046527 | COX8A:NDUFA6:PPAT | Pubmed 20605485 Authors: Marigo I,Bosio E,Solito S,Mesa C,Fernandez A,Dolcetti L,Ugel S,Sonda N,Bicciato S,Falisi E,Calabrese F,Basso G,Zanovello P,Cozzi E,Mandruzzato S,Bronte V | Mus musculus |
| GSE15930 NAIVE VS 72H IN VITRO STIM TRICHOSTATINA CD8 TCELL UP | Genes up-regulated in comparison of CD8 T cells at 0 h versus those at 72 h after treatment with trichostatin A (TSA) | 200 | 3 | 0.000726 | 0.046527 | PAPSS2:POLG2:ACVR2A | Pubmed 19592655 Authors: Agarwal P,Raghavan A,Nandiwada SL,Curtsinger JM,Bohjanen PR,Mueller DL,Mescher MF. | Mus musculus |
| GSE24726 WT VS E2 22 KO PDC DAY4 POST DELETION DN | Genes down-regulated in plasmacytoid dendritic cells (4 days after knockout): wildtype versus TCF4 knockout. | 200 | 3 | 0.000726 | 0.046527 | IL1B:DDX39B:JAK2 | Pubmed 21145760 Authors: Ghosh HS,Cisse B,Bunin A,Lewis KL,Reizis B | Mus musculus |
| GSE21360 SECONDARY VS TERTIARY MEMORY CD8 TCELL UP | Genes up-regulated in memory CD8 T cells: 2' versus 3'. | 200 | 3 | 0.000726 | 0.046527 | NDUFS3:COX8A:MYBBP1A | Pubmed 20619696 Authors: Wirth TC,Xue HH,Rai D,Sabel JT,Bair T,Harty JT,Badovinac VP | Mus musculus |
| GSE14308 TH2 VS NAIVE CD4 TCELL UP | Genes up-regulated in comparison of Th2 cells versus naive CD4 T cells. | 200 | 3 | 0.000726 | 0.046527 | CTPS1:SDHC:PAICS | Pubmed 19144320 Authors: Wei G,Wei L,Zhu J,Zang C,Hu-Li J,Yao Z,Cui K,Kanno Y,Roh TY,Watford WT,Schones DE,Peng W,Sun HW,Paul WE,O'Shea JJ,Zhao K. | Mus musculus |

Supplementary table 6: Immunologic Signatures Database (MsigDB c7). N indicates the total number of genes in the set while n shows the number of genes that SNPs mapped to the 37 genes identified. Gene sets were queried using FUMA GWAS. The adjusted P is an FDR-corrected P-value based on the number of gene sets examined.

| Gene Set | Gene sets from which the module was originally composed | N | n | P-value | adj. P-value | Genes |
|---|---|---|---|---|---|---|
| MODULE 62 | ion transporter activity<br>mitochondrial membrane<br>inner membrane<br>hydrogen ion transporter activity<br>Oxidative phosphorylation<br>monovalent inorganic cation transporter activity<br>sodium ion transporter activity<br>metal ion transporter activity<br>Electron Transport Chain | 89 | 5 | 1.50E-08 | 6.46E-06 | SDHC:NDUFS3:COX8A:<br>NDUFA6:NDUFA5 |
| MODULE 42 | NADH dehydrogenase activity<br>NADH dehydrogenase (ubiquinone) activity | 25 | 3 | 1.41E-06 | 0.000304 | NDUFS3:NDUFA6:NDU<br>FA5 |
| MODULE 278 -<br>Ubiquitin ligases | protein phosphatase type 2A activity<br>acid D amino acid ligase activity<br>ligase activity, forming carbon nitrogen bonds | 34 | 3 | 3.65E-06 | 0.000465 | CTPS1:GSS:PAICS |

| | | | | | | |
|---|---|---|---|---|---|---|
| MODULE 152 - Oxidative phosphorylation and ATP synthesis | nucleoside monophosphate phosphorylation<br>ATP Synthesis<br>proton transport<br>mitochondrial inner membrane<br>cytochrome c oxidase activity<br>cbb3 type cytochrome c oxidase<br>NADH dehydrogenase activity<br>NADH dehydrogenase (ubiquinone) activity<br>Ubiquinone biosynthesis<br>protein disulfide reduction<br>mitochondrial electron transport, NADH to ubiquinone<br>ATP synthesis coupled electron transport (sensu Eukarya)<br>oxidative phosphorylation<br>mitochondrion<br>ion transporter activity<br>mitochondrial membrane<br>inner membrane<br>hydrogen ion transporter activity<br>Oxidative phosphorylation<br>monovalent inorganic cation transporter activity<br>sodium ion transporter activity<br>metal ion transporter activity<br>Electron Transport Chain | 124 | 4 | 4.32E-06 | 0.000465 | NDUFS3:COX8A:NDUFA6:NDUFA5 |
| MODULE 22 | sodium ion transporter activity<br>metal ion transporter activity | 45 | 3 | 8.60E-06 | 0.000741 | NDUFS3:NDUFA6:NDUFA5 |

| | | | | | | |
|---|---|---|---|---|---|---|
| MODULE 126 - Lymphoma and immune response expression clusters | Rosenwald01 45<br>Alizadeh00 42<br>Boldrick02 28<br>Boldrick02 16<br>Boldrick02 12 | 182 | 4 | 1.96E-05 | 0.00141 | CTPS1:PPAT:PAICS:CCNB1 |
| MODULE 102 - Nucleotide (purine) biosynthesis | ribonucleoside monophosphate metabolism<br>nucleoside monophosphate biosynthesis<br>ribonucleoside monophosphate biosynthesis<br>purine ribonucleoside monophosphate metabolism<br>purine nucleoside monophosphate biosynthesis<br>purine ribonucleoside monophosphate biosynthesis<br>purine ribonucleotide biosynthesis<br>ribonucleotide biosynthesis<br>nucleotide biosynthesis | 19 | 2 | 0.000128 | 0.007856 | CTPS1:PAICS |
| MODULE 233 | protein phosphatase type 2A activity<br>acid_D_amino acid ligase activity | 22 | 2 | 0.000172 | 0.00927 | GSS:PAICS |
| MODULE 219 | ribonucleotide metabolism<br>nucleotide metabolism<br>purine nucleotide biosynthesis<br>ribonucleoside monophosphate metabolism<br>nucleoside monophosphate biosynthesis<br>ribonucleoside monophosphate biosynthesis<br>purine ribonucleoside monophosphate metabolism<br>purine nucleoside monophosphate biosynthesis<br>purine ribonucleoside monophosphate biosynthesis<br>purine ribonucleotide biosynthesis<br>ribonucleotide biosynthesis<br>nucleotide biosynthesis<br>Purine_metabolism | 27 | 2 | 0.000261 | 0.011587 | CTPS1:PAICS |

| | | | | | | |
|---|---|---|---|---|---|---|
| MODULE 77 | NADH dehydrogenase activity<br>NADH dehydrogenase (ubiquinone) activity<br>Ubiquinone biosynthesis<br>protein disulfide reduction<br>mitochondrial electron transport, NADH to ubiquinone<br>ATP synthesis coupled electron transport (sensu Eukarya)<br>oxidative phosphorylation | 28 | 2 | 0.000281 | 0.011587 | NDUFS3:NDUFA5 |
| MODULE 184 | Propanoate metabolism<br>Fatty Acid Degradation<br>acyl CoA dehydrogenase activity<br>oxidoreductase activity, acting on the CH CH group of donors | 29 | 2 | 0.000301 | 0.011587 | SDHC:ALDH2 |
| MODULE 221 - Fatty Acid Metabolism | Propanoate metabolism<br>Fatty Acid Degradation<br>acyl CoA dehydrogenase activity<br>oxidoreductase activity, acting on the CH CH group of donors<br>fatty acid oxidation<br>fatty acid beta oxidation | 30 | 2 | 0.000323 | 0.011587 | SDHC:ALDH2 |
| MODULE 105 | Shipp02 28<br>Ramaswamy01 28<br>Shipp02 5<br>Golub99 15 | 200 | 3 | 0.000726 | 0.024063 | CTPS1:PAICS:CCNB1 |
| MODULE 312 - Viral anti-apoptotic evasion mechanisms | viral host defense evasion<br>host pathogen interaction<br>Garber01 35 | 47 | 2 | 0.000794 | 0.024451 | IL1B:CCNB1 |

| | | | | | | |
|---|---|---|---|---|---|---|
| MODULE 61 | Rosenwald01 45<br>Alizadeh00 42 | 50 | 2 | 0.000899 | 0.025179 | CTPS1:PAICS |
| MODULE 85 | transmembrane receptor protein tyrosine kinase activity<br>transmembrane receptor protein kinase activity | 51 | 2 | 0.000935 | 0.025179 | ACVR2A:KDR |
| MODULE 234 - Bone Remodeling | skeletal development<br>bone remodeling | 54 | 2 | 0.001047 | 0.026553 | IGF2:JAK2 |
| MODULE 254 | viral host defense evasion<br>host pathogen interaction | 60 | 2 | 0.001291 | 0.030917 | IL1B:CCNB1 |

Supplementary table 7: Cancer gene modules Database (MsigDB c4). N indicates the total number of genes in the set while n shows the number of genes that SNPs mapped to the 37 genes identified. Gene sets were queried using FUMA GWAS. The adjusted P is an FDR-corrected P-value based on the number of gene sets examined.

| Gene Set | Brief Description/Gene Sets from which the module was originally composed | N | n | P-value | adj. P-value | Genes |
|---|---|---|---|---|---|---|
| MODULE 62 | ion transporter activity<br>mitochondrial membrane<br>inner membrane<br>hydrogen ion transporter activity<br>Oxidative phosphorylation<br>monovalent inorganic cation transporter activity<br>sodium ion transporter activity<br>metal ion transporter activity<br>Electron Transport Chain | 89 | 5 | 1.50E-08 | 1.29E-05 | SDHC:NDUFS3:COX8A:NDUFA6:NDUFA5 |
| MODULE 42 | NADH dehydrogenase activity<br>NADH dehydrogenase (ubiquinone) activity | 25 | 3 | 1.41E-06 | 0.000605 | NDUFS3:NDUFA6:NDUFA5 |
| MODULE 278 - Ubiquitin ligases | protein phosphatase type 2A activity<br>acid D amino acid ligase activity<br>ligase activity, forming carbon nitrogen bonds | 34 | 3 | 3.65E-06 | 0.000926 | CTPS1:GSS:PAICS |

| | | | | | | |
|---|---|---|---|---|---|---|
| MODULE 152 - Oxidative phosphorylation and ATP synthesis | nucleoside monophosphate phosphorylation<br>ATP Synthesis<br>proton transport<br>mitochondrial inner membrane<br>cytochrome c oxidase activity<br>cbb3 type cytochrome c oxidase<br>NADH dehydrogenase activity<br>NADH dehydrogenase (ubiquinone) activity<br>Ubiquinone biosynthesis<br>protein disulfide reduction<br>mitochondrial electron transport, NADH to ubiquinone<br>ATP synthesis coupled electron transport (sensu Eukarya)<br>oxidative phosphorylation<br>mitochondrion<br>ion transporter activity<br>mitochondrial membrane<br>inner membrane<br>hydrogen ion transporter activity<br>Oxidative phosphorylation<br>monovalent inorganic cation transporter activity<br>sodium ion transporter activity<br>metal ion transporter activity<br>Electron Transport Chain | 124 | 4 | 4.32E-06 | 0.000926 | NDUFS3:COX8A:NDUFA6:NDUFA5 |
| GNF2 RFC3 | Neighborhood of RFC3 replication factor C (activator 1) 3, 38kDa in the GNF2 expression compendium | 41 | 3 | 6.48E-06 | 0.001111 | CTPS1:PFAS:PAICS |
| MODULE 22 | sodium ion transporter activity<br>metal ion transporter activity | 45 | 3 | 8.60E-06 | 0.00123 | NDUFS3:NDUFA6:NDUFA5 |

| | | | | | | |
|---|---|---|---|---|---|---|
| MODULE 126 - Lymphoma and immune response expression clusters | Rosenwald01 45<br>Alizadeh00 42<br>Boldrick02 28<br>Boldrick02 16<br>Boldrick02 12 | 182 | 4 | 1.96E-05 | 0.002406 | CTPS1:PPAT:PAICS:CCNB1 |
| MORF DAP3 | Neighborhood of DAP3 death associated protein 3 in the MORF expression compendium | 195 | 4 | 2.57E-05 | 0.002759 | NDUFS3:COX8A:COX7A2L:DDX39B |
| MORF UBE2I | Neighborhood of UBE2I ubiquitin-conjugating enzyme E2I (UBC9 homolog, yeast) in the MORF expression compendium | 238 | 4 | 5.59E-05 | 0.00533 | NDUFS3:COX8A:COX7A2L:DDX39B |
| MORF RAN | Neighborhood of RAN RAN, member RAS oncogene family in the MORF expression compendium | 267 | 4 | 8.72E-05 | 0.007484 | NDUFS3:COX8A:COX7A2L:DDX39B |
| | ribonucleoside monophosphate metabolism<br>nucleoside monophosphate biosynthesis<br>ribonucleoside monophosphate biosynthesis<br>purine ribonucleoside monophosphate metabolism<br>purine nucleoside monophosphate biosynthesis<br>purine ribonucleoside monophosphate biosynthesis<br>purine ribonucleotide biosynthesis<br>ribonucleotide biosynthesis<br>nucleotide biosynthesis | 19 | 2 | 0.000128 | 0.009952 | CTPS1:PAICS |
| MODULE 233 | protein phosphatase type 2A activity<br>acid_D_amino acid ligase activity | 22 | 2 | 0.000172 | 0.012303 | GSS:PAICS |

| | | | | | | |
|---|---|---|---|---|---|---|
| | ribonucleotide metabolism<br>nucleotide metabolism<br>purine nucleotide biosynthesis<br>ribonucleoside monophosphate metabolism<br>nucleoside monophosphate biosynthesis<br>ribonucleoside monophosphate biosynthesis<br>purine ribonucleoside monophosphate metabolism<br>purine nucleoside monophosphate biosynthesis<br>purine ribonucleoside monophosphate biosynthesis<br>purine ribonucleotide biosynthesis<br>ribonucleotide biosynthesis<br>nucleotide biosynthesis<br>Purine_metabolism | 27 | 2 | 0.000261 | 0.016053 | CTPS1:PAICS |
| MORF PRKAR1A | Neighborhood of PRKAR1A protein kinase, cAMP-dependent, regulatory, type I, alpha (tissue specific extinguisher 1) in the MORF expression compendium | 142 | 3 | 0.000267 | 0.016053 | SDHC:COX8A:COX7A2L |
| MODULE 77 | NADH dehydrogenase activity<br>NADH dehydrogenase (ubiquinone) activity<br>Ubiquinone biosynthesis<br>protein disulfide reduction<br>mitochondrial electron transport, NADH to ubiquinone<br>ATP synthesis coupled electron transport (sensu Eukarya)<br>oxidative phosphorylation | 28 | 2 | 0.000281 | 0.016053 | NDUFS3:NDUFA5 |
| MODULE 184 | Propanoate metabolism<br>Fatty Acid Degradation<br>acyl CoA dehydrogenase activity<br>oxidoreductase activity, acting on the CH CH group of donors | 29 | 2 | 0.000301 | 0.016155 | SDHC:ALDH2 |

| | | | | | | |
|---|---|---|---|---|---|---|
| MODULE 221 - Fatty Acid Metabolism | Propanoate metabolism<br>Fatty Acid Degradation<br>acyl CoA dehydrogenase activity<br>oxidoreductase activity, acting on the CH CH group of donors<br>fatty acid oxidation<br>fatty acid beta oxidation | 30 | 2 | 0.000323 | 0.016282 | SDHC:ALDH2 |
| MORF RAD21 | Neighborhood of RAD21 RAD21 homolog (S. pombe) in the MORF expression compendium | 179 | 3 | 0.000526 | 0.02507 | NDUFS3:COX7A2L:NDUFA5 |
| MORF RAB1A | Neighborhood of RAB1A RAB1A, member RAS oncogene family in the MORF expression compendium | 191 | 3 | 0.000635 | 0.02868 | SDHC:COX8A:COX7A2L |
| MORF ANP32B | Neighborhood of ANP32B acidic (leucine-rich) nuclear phosphoprotein 32 family, member B in the MORF expression compendium | 198 | 3 | 0.000705 | 0.029553 | NDUFS3:COX7A2L:DDX39B |
| MODULE 105 | Shipp02 28<br>Ramaswamy01 28<br>Shipp02 5<br>Golub99 15 | 200 | 3 | 0.000726 | 0.029553 | CTPS1:PAICS:CCNB1 |
| MORF SKP1A | Neighborhood of SKP1A S-phase kinase-associated protein 1A (p19A) in the MORF expression compendium | 203 | 3 | 0.000758 | 0.029553 | NDUFS3:COX7A2L:NDUFA5 |
| MODULE 312 - Viral anti-apoptotic evasion mechanisms | viral host defense evasion<br>host pathogen interaction<br>Garber01 35 | 47 | 2 | 0.000794 | 0.029628 | IL1B:CCNB1 |
| MORF RAC1 | Neighborhood of RAC1 ras-related C3 botulinum toxin substrate 1 (rho family, small GTP binding protein Rac1) in the MORF expression compendium | 211 | 3 | 0.000847 | 0.030293 | SDHC:COX8A:COX7A2L |
| MODULE_61 | Rosenwald01 45<br>Alizadeh00 42 | 50 | 2 | 0.000899 | 0.030838 | CTPS1:PAICS |
| MODULE 85 | transmembrane receptor protein tyrosine kinase activity<br>transmembrane receptor protein kinase activity | 51 | 2 | 0.000935 | 0.030846 | ACVR2A:KDR |
| MODULE 234 - Bone Remodeling | skeletal development<br>bone remodeling | 54 | 2 | 0.001047 | 0.033283 | IGF2:JAK2 |

| | | | | | | |
|---|---|---|---|---|---|---|
| | Neighborhood of RFC4 replication factor C (activator 1) 4, 37kDa in the GNF2 expression compendium | 60 | 2 | 0.001291 | 0.036929 | CTPS1:PAICS |
| MORF RAB11A | Neighborhood of RAB11A RAB11A, member RAS oncogene family in the MORF expression compendium | 60 | 2 | 0.001291 | 0.036929 | SDHC:COX7A2L |
| MODULE 254 | viral host defense evasion<br>host pathogen interaction | 60 | 2 | 0.001291 | 0.036929 | IL1B:CCNB1 |
| MORF SMC1L1 | Neighborhood of SMC1L1 NULL in the MORF expression compendium | 62 | 2 | 0.001378 | 0.038139 | NDUFS3:DDX39B |
| MORF DEK | Neighborhood of DEK DEK oncogene (DNA binding) in the MORF expression compendium | 264 | 3 | 0.001613 | 0.043243 | NDUFS3:COX7A2L:DDX39B |
| MORF BUB3 | Neighborhood of BUB3 BUB3 budding uninhibited by benzimidazoles 3 homolog (yeast) in the MORF expression compendium | 277 | 3 | 0.00185 | 0.04809 | NDUFS3:PAICS:DDX39B |

Supplementary table 8: Computational Gene Set Database (MsigDB c4). N indicates the total number of genes in the set while n shows the number of genes that SNPs mapped to the 37 genes identified. Gene sets were queried using FUMA GWAS. The adjusted P is an FDR-corrected P-value based on the number of gene sets examined.

| Gene Set | Brief description | N | n | P-value | adj. P-value | Genes |
|---|---|---|---|---|---|---|
| ELECTRON TRANSFER ACTIVITY | Any molecular entity that serves as an electron acceptor and electron donor in an electron transport chain. An electron transport chain is a process in which a series of electron carriers operate together to transfer electrons from donors to any of several different terminal electron acceptors to generate a transmembrane electrochemical gradient. | 109 | 7 | 5.62E-12 | 9.24E-09 | SDHC:NDUFS3:COX8A:ALDH2:CYP1A2:COX7A2L:GLDC |
| OXIDOREDUCTASE ACTIVITY | Catalysis of an oxidation-reduction (redox) reaction, a reversible chemical reaction in which the oxidation state of an atom or atoms within a molecule is altered. One substrate acts as a hydrogen or electron donor and becomes oxidized, while the other acts as hydrogen or electron acceptor and becomes reduced. | 731 | 9 | 9.30E-09 | 7.65E-06 | SDHC:NDUFS3:COX8A:ALDH2:CYP1A2:COX7A2L:NDUFA6:NDUFA5:GLDC |
| LIGASE ACTIVITY FORMING CARBON NITROGEN BONDS | Catalysis of the joining of two molecules, or two groups within a single molecule, via a carbon-nitrogen bond, with the concomitant hydrolysis of the diphosphate bond in ATP or a similar triphosphate. | 43 | 4 | 5.97E-08 | 3.27E-05 | CTPS1:PFAS:GSS:PAICS |
| DRUG BINDING | Binding to a drug, a naturally occurring or synthetic substance, other than a nutrient, that, when administered or applied to an organism, affects the structure or functioning of the organism;typically used in the diagnosis, prevention, or treatment of disease | 1718 | 11 | 1.26E-07 | 5.19E-05 | CTPS1:PAPSS2:PFAS:GSK3A:ACVR2A:GSS:KDR:PAICS:DDX39B:JAK2:GLDC |
| COFACTOR BINDING | Interacting selectively with a cofactor, a substance that is required for the activity of an enzyme or other protein. Cofactors may be inorganic, such as the metal atoms zinc, iron, and copper in certain forms, or organic, in which case they are referred to as coenzymes. Cofactors may either be bound tightly to active sites or bind loosely with the substrate. | 495 | 7 | 2.07E-07 | 6.81E-05 | SDHC:ALDH2:CYP1A2:GSS:PPAT:JAK2:GLDC |
| ADENYL NUCLEOTIDE BINDING | Binding to an adenyl nucleotide, an adenosine esterified with (ortho)phosphate. | 1536 | 10 | 4.72E-07 | 0.00013 | CTPS1:PAPSS2:PFAS:GSK3A:ACVR2A:GSS:KDR:PAICS:DDX39B:JAK2 |

| | | | | | | |
|---|---|---|---|---|---|---|
| RIBONUCLEOTIDE BINDING | Binding to a ribonucleotide, any compound consisting of a ribonucleoside that is esterified with (ortho)phosphate or an oligophosphate at any hydroxyl group on the ribose moiety. | 1885 | 10 | 3.02E-06 | 0.00071 | CTPS1:PAPSS2:PFAS:GSK3A:ACVR2A:GSS:KDR:PAICS:DDX39B:JAK2 |
| NADH DEHYDROGENASE ACTIVITY | Catalysis of the reaction: NADH + H+ + acceptor = NAD+ + reduced acceptor. | 40 | 3 | 6.01E-06 | 0.001235 | NDUFS3:NDUFA6:NDUFA5 |
| SIGNALING RECEPTOR BINDING | Binding to one or more specific sites on a receptor molecule, a macromolecule that undergoes combination with a hormone, neurotransmitter, drug or intracellular messenger to initiate a change in cell function. | 1619 | 9 | 7.34E-06 | 0.001341 | IGF2:INS-IGF2:INS:GSK3A:IL1B:ACVR2A:KDR:CCNB1:JAK2 |
| LIGASE ACTIVITY | Catalysis of the joining of two molecules, or two groups within a single molecule, using the energy from the hydrolysis of ATP, a similar triphosphate, or a pH gradient. | 146 | 4 | 8.24E-06 | 0.001356 | CTPS1:PFAS:GSS:PAICS |
| IDENTICAL PROTEIN BINDING | Binding to an identical protein or proteins. | 1706 | 9 | 1.12E-05 | 0.001575 | CTPS1:INS:POLG2:GSS:KDR:PAICS:DDX39B:JAK2:GLDC |
| TRANSFERASE ACTIVITY TRANSFERRING PHOSPHORUS CONTAINING GROUPS | Catalysis of the transfer of a phosphorus-containing group from one compound (donor) to another (acceptor). | 908 | 7 | 1.15E-05 | 0.001575 | PAPSS2:POLG2:GSK3A:ACVR2A:KDR:CCNB1:JAK2 |
| OXIDOREDUCTASE ACTIVITY ACTING ON NAD P H QUINONE OR SIMILAR COMPOUND AS ACCEPTOR | Catalysis of an oxidation-reduction (redox) reaction in which NADH or NADPH acts as a hydrogen or electron donor and reduces a quinone or a similar acceptor molecule. | 53 | 3 | 1.41E-05 | 0.001788 | NDUFS3:NDUFA6:NDUFA5 |
| KINASE ACTIVITY | Catalysis of the transfer of a phosphate group, usually from ATP, to a substrate molecule. | 758 | 6 | 4.59E-05 | 0.005391 | PAPSS2:GSK3A:ACVR2A:KDR:CCNB1:JAK2 |
| GLYCINE BINDING | Binding to glycine, aminoethanoic acid. | 15 | 2 | 7.85E-05 | 0.008072 | GSS:GLDC |
| ACID AMINO ACID LIGASE ACTIVITY | Catalysis of the ligation of an acid to an amino acid via a carbon-nitrogen bond, with the concomitant hydrolysis of the diphosphate bond in ATP or a similar triphosphate. | 15 | 2 | 7.85E-05 | 0.008072 | GSS:PAICS |

| | | | | | | |
|---|---|---|---|---|---|---|
| INSULIN LIKE GROWTH FACTOR RECEPTOR BINDING | Binding to an insulin-like growth factor receptor. | 16 | 2 | 8.97E-05 | 0.008678 | IGF2:INS |
| OXIDOREDUCTASE ACTIVITY ACTING ON NAD P H | Catalysis of an oxidation-reduction (redox) reaction in which NADH or NADPH acts as a hydrogen or electron donor and reduces a quinone or a similar acceptor molecule. | 101 | 3 | 9.77E-05 | 0.008795 | NDUFS3:NDUFA6:NDUFA5 |
| HISTONE KINASE ACTIVITY | Catalysis of the transfer of a phosphate group to a histone. Histones are any of a group of water-soluble proteins found in association with the DNA of eukaroytic chromosomes. | 17 | 2 | 0.000102 | 0.008795 | CCNB1:JAK2 |
| PROTEIN KINASE ACTIVITY | Catalysis of the phosphorylation of an amino acid residue in a protein, usually according to the reaction: a protein + ATP = a phosphoprotein + ADP. | 588 | 5 | 0.000153 | 0.012573 | GSK3A:ACVR2A:KDR:CCNB1:JAK2 |
| HORMONE ACTIVITY | The action characteristic of a hormone, any substance formed in very small amounts in one specialized organ or group of cells and carried (sometimes in the bloodstream) to another organ or group of cells in the same organism, upon which it has a specific regulatory action. The term was originally applied to agents with a stimulatory physiological action in vertebrate animals (as opposed to a chalone, which has a depressant action). Usage is now extended to regulatory compounds in lower animals and plants, and to synthetic substances having comparable effects; all bind receptors and trigger some biological process. | 121 | 3 | 0.000167 | 0.012866 | IGF2:INS-IGF2:INS |
| INSULIN RECEPTOR BINDING | Binding to an insulin receptor. | 22 | 2 | 0.000172 | 0.012866 | IGF2:INS |
| INTEGRIN BINDING | Binding to an integrin. | 127 | 3 | 0.000192 | 0.013497 | IGF2:IL1B:KDR |
| PROTON TRANSMEMBRANE TRANSPORTER ACTIVITY | Enables the transfer of a proton from one side of a membrane to the other. | 128 | 3 | 0.000197 | 0.013497 | COX8A:COX7A2L:ATP6V1G2 |
| OXIDOREDUCTASE ACTIVITY ACTING ON A HEME GROUP OF DONORS | Catalysis of an oxidation-reduction (redox) reaction in which a heme group acts as a hydrogen or electron donor and reduces a hydrogen or electron acceptor. | 26 | 2 | 0.000242 | 0.015894 | COX8A:COX7A2L |

| | | | | | | |
|---|---|---|---|---|---|---|
| TETRAPYRROLE BINDING | Binding to a tetrapyrrole, a compound containing four pyrrole nuclei variously substituted and linked to each other through carbons at the alpha position. | 141 | 3 | 0.000262 | 0.016561 | SDHC:CYP1A2:JAK2 |
| PROTEIN CONTAINING COMPLEX BINDING | Binding to a macromolecular complex. | 1096 | 6 | 0.000342 | 0.020867 | IGF2:INS:IL1B:ACVR2A:KDR:DDX39B |

Supplementary table 9: Wikipathways Database. N indicates the total number of genes in the set while n shows the number of genes that SNPs mapped to the 37 genes identified. Gene sets were queried using FUMA GWAS. The adjusted P is an FDR-corrected P-value based on the number of gene sets examined.

| Gene Set | Brief description | N | n | P-value | adj. P-value | Genes |
|---|---|---|---|---|---|---|
| ELECTRON TRANSFER ACTIVITY | Any molecular entity that serves as an electron acceptor and electron donor in an electron transport chain. An electron transport chain is a process in which a series of electron carriers operate together to transfer electrons from donors to any of several different terminal electron acceptors to generate a transmembrane electrochemical gradient. | 109 | 7 | 5.62E-12 | 9.24E-09 | SDHC:NDUFS3:COX8A:ALDH2:CYP1A2:COX7A2L:GLDC |
| OXIDOREDUCTASE ACTIVITY | Catalysis of an oxidation-reduction (redox) reaction, a reversible chemical reaction in which the oxidation state of an atom or atoms within a molecule is altered. One substrate acts as a hydrogen or electron donor and becomes oxidized, while the other acts as hydrogen or electron acceptor and becomes reduced. | 731 | 9 | 9.30E-09 | 7.65E-06 | SDHC:NDUFS3:COX8A:ALDH2:CYP1A2:COX7A2L:NDUFA6:NDUFA5:GLDC |
| LIGASE ACTIVITY FORMING CARBON NITROGEN BONDS | Catalysis of the joining of two molecules, or two groups within a single molecule, via a carbon-nitrogen bond, with the concomitant hydrolysis of the diphosphate bond in ATP or a similar triphosphate. | 43 | 4 | 5.97E-08 | 3.27E-05 | CTPS1:PFAS:GSS:PAICS |
| DRUG BINDING | Binding to a drug, a naturally occurring or synthetic substance, other than a nutrient, that, when administered or applied to an organism, affects the structure or functioning of the organism;typically used in the diagnosis, prevention, or treatment of disease | 1718 | 11 | 1.26E-07 | 5.19E-05 | CTPS1:PAPSS2:PFAS:GSK3A:ACVR2A:GSS:KDR:PAICS:DDX39B:JAK2:GLDC |
| COFACTOR BINDING | Interacting selectively with a cofactor, a substance that is required for the activity of an enzyme or other protein. Cofactors may be inorganic, such as the metal atoms zinc, iron, and copper in certain forms, or organic, in which case they are referred to as coenzymes. Cofactors may either be bound tightly to active sites or bind loosely with the substrate. | 495 | 7 | 2.07E-07 | 6.81E-05 | SDHC:ALDH2:CYP1A2:GSS:PPAT:JAK2:GLDC |
| ADENYL NUCLEOTIDE BINDING | Binding to an adenyl nucleotide, an adenosine esterified with (ortho)phosphate. | 1536 | 10 | 4.72E-07 | 0.00013 | CTPS1:PAPSS2:PFAS:GSK3A:ACVR2A:GSS:KDR:PAICS:DDX39B:JAK2 |

| | | | | | | |
|---|---|---|---|---|---|---|
| RIBONUCLEOTIDE BINDING | Binding to a ribonucleotide, any compound consisting of a ribonucleoside that is esterified with (ortho)phosphate or an oligophosphate at any hydroxyl group on the ribose moiety. | 1885 | 10 | 3.02E-06 | 0.00071 | CTPS1:PAPSS2:PFAS:GSK3A:ACVR2A:GSS:KDR:PAICS:DDX39B:JAK2 |
| NADH DEHYDROGENASE ACTIVITY | Catalysis of the reaction: NADH + H+ + acceptor = NAD+ + reduced acceptor. | 40 | 3 | 6.01E-06 | 0.001235 | NDUFS3:NDUFA6:NDUFA5 |
| SIGNALING RECEPTOR BINDING | Binding to one or more specific sites on a receptor molecule, a macromolecule that undergoes combination with a hormone, neurotransmitter, drug or intracellular messenger to initiate a change in cell function. | 1619 | 9 | 7.34E-06 | 0.001341 | IGF2:INS-IGF2:INS:GSK3A:IL1B:ACVR2A:KDR:CCNB1:JAK2 |
| LIGASE ACTIVITY | Catalysis of the joining of two molecules, or two groups within a single molecule, using the energy from the hydrolysis of ATP, a similar triphosphate, or a pH gradient. | 146 | 4 | 8.24E-06 | 0.001356 | CTPS1:PFAS:GSS:PAICS |
| IDENTICAL PROTEIN BINDING | Binding to an identical protein or proteins. | 1706 | 9 | 1.12E-05 | 0.001575 | CTPS1:INS:POLG2:GSS:KDR:PAICS:DDX39B:JAK2:GLDC |
| TRANSFERASE ACTIVITY TRANSFERRING PHOSPHORUS CONTAINING GROUPS | Catalysis of the transfer of a phosphorus-containing group from one compound (donor) to another (acceptor). | 908 | 7 | 1.15E-05 | 0.001575 | PAPSS2:POLG2:GSK3A:ACVR2A:KDR:CCNB1:JAK2 |
| OXIDOREDUCTASE ACTIVITY ACTING ON NAD P H QUINONE OR SIMILAR COMPOUND AS ACCEPTOR | Catalysis of an oxidation-reduction (redox) reaction in which NADH or NADPH acts as a hydrogen or electron donor and reduces a quinone or a similar acceptor molecule. | 53 | 3 | 1.41E-05 | 0.001788 | NDUFS3:NDUFA6:NDUFA5 |
| KINASE ACTIVITY | Catalysis of the transfer of a phosphate group, usually from ATP, to a substrate molecule. | 758 | 6 | 4.59E-05 | 0.005391 | PAPSS2:GSK3A:ACVR2A:KDR:CCNB1:JAK2 |
| GLYCINE BINDING | Binding to glycine, aminoethanoic acid. | 15 | 2 | 7.85E-05 | 0.008072 | GSS:GLDC |
| ACID AMINO ACID LIGASE ACTIVITY | Catalysis of the ligation of an acid to an amino acid via a carbon-nitrogen bond, with the concomitant | 15 | 2 | 7.85E-05 | 0.008072 | GSS:PAICS |

| | | | | | | |
|---|---|---|---|---|---|---|
| | hydrolysis of the diphosphate bond in ATP or a similar triphosphate. | | | | | |
| INSULIN LIKE GROWTH FACTOR RECEPTOR BINDING | Binding to an insulin-like growth factor receptor. | 16 | 2 | 8.97E-05 | 0.008678 | IGF2:INS |
| OXIDOREDUCTASE ACTIVITY ACTING ON NAD P H | Catalysis of an oxidation-reduction (redox) reaction in which NADH or NADPH acts as a hydrogen or electron donor and reduces a quinone or a similar acceptor molecule. | 101 | 3 | 9.77E-05 | 0.008795 | NDUFS3:NDUFA6:NDUFA5 |
| HISTONE KINASE ACTIVITY | Catalysis of the transfer of a phosphate group to a histone. Histones are any of a group of water-soluble proteins found in association with the DNA of eukaroytic chromosomes. | 17 | 2 | 0.000102 | 0.008795 | CCNB1:JAK2 |
| PROTEIN KINASE ACTIVITY | Catalysis of the phosphorylation of an amino acid residue in a protein, usually according to the reaction: a protein + ATP = a phosphoprotein + ADP. | 588 | 5 | 0.000153 | 0.012573 | GSK3A:ACVR2A:KDR:CCNB1:JAK2 |
| HORMONE ACTIVITY | The action characteristic of a hormone, any substance formed in very small amounts in one specialized organ or group of cells and carried (sometimes in the bloodstream) to another organ or group of cells in the same organism, upon which it has a specific regulatory action. The term was originally applied to agents with a stimulatory physiological action in vertebrate animals (as opposed to a chalone, which has a depressant action). Usage is now extended to regulatory compounds in lower animals and plants, and to synthetic substances having comparable effects; all bind receptors and trigger some biological process. | 121 | 3 | 0.000167 | 0.012866 | IGF2:INS-IGF2:INS |
| INSULIN RECEPTOR BINDING | Binding to an insulin receptor. | 22 | 2 | 0.000172 | 0.012866 | IGF2:INS |
| INTEGRIN BINDING | Binding to an integrin. | 127 | 3 | 0.000192 | 0.013497 | IGF2:IL1B:KDR |
| PROTON TRANSMEMBRANE | Enables the transfer of a proton from one side of a membrane to the other. | 128 | 3 | 0.000197 | 0.013497 | COX8A:COX7A2L:ATP6V1G2 |

| | | | | | | |
|---|---|---|---|---|---|---|
| TRANSPORTER ACTIVITY | | | | | | |
| OXIDOREDUCTASE ACTIVITY ACTING ON A HEME GROUP OF DONORS | Catalysis of an oxidation-reduction (redox) reaction in which a heme group acts as a hydrogen or electron donor and reduces a hydrogen or electron acceptor. | 26 | 2 | 0.000242 | 0.015894 | COX8A:COX7A2L |
| TETRAPYRROLE BINDING | Binding to a tetrapyrrole, a compound containing four pyrrole nuclei variously substituted and linked to each other through carbons at the alpha position. | 141 | 3 | 0.000262 | 0.016561 | SDHC:CYP1A2:JAK2 |
| PROTEIN CONTAINING COMPLEX BINDING | Binding to a macromolecular complex. | 1096 | 6 | 0.000342 | 0.020867 | IGF2:INS:IL1B:ACVR2A:KDR:DDX39B |

Supplementary table 10: GO Molecular Functions Database (MsigDB c5). N indicates the total number of genes in the set while n shows the number of genes that SNPs mapped to the 37 genes identified. Gene sets were queried using FUMA GWAS. The adjusted P is an FDR-corrected P-value based on the number of gene sets examined.

| Gene Set | Brief description | N | n | P-value | adj. P-value | Genes |
|---|---|---|---|---|---|---|
| GNF2 RFC3 | Neighborhood of RFC3 replication factor C (activator 1) 3, 38kDa in the GNF2 expression compendium | 41 | 3 | 6.48E-06 | 0.002766 | CTPS1:PFAS:PAICS |
| MORF DAP3 | Neighborhood of DAP3 death associated protein 3 in the MORF expression compendium | 195 | 4 | 2.57E-05 | 0.005492 | NDUFS3:COX8A:COX7A2L:DDX39B |
| MORF UBE2I | Neighborhood of UBE2I ubiquitin-conjugating enzyme E2I (UBC9 homolog, yeast) in the MORF expression compendium | 238 | 4 | 5.59E-05 | 0.007958 | NDUFS3:COX8A:COX7A2L:DDX39B |
| MORF RAN | Neighborhood of RAN RAN, member RAS oncogene family in the MORF expression compendium | 267 | 4 | 8.72E-05 | 0.009312 | NDUFS3:COX8A:COX7A2L:DDX39B |
| MORF PRKAR1A | Neighborhood of PRKAR1A protein kinase, cAMP-dependent, regulatory, type I, alpha (tissue specific extinguisher 1) in the MORF expression compendium | 142 | 3 | 0.000267 | 0.022823 | SDHC:COX8A:COX7A2L |
| MORF RAD21 | Neighborhood of RAD21 RAD21 homolog (S. pombe) in the MORF expression compendium | 179 | 3 | 0.000526 | 0.035952 | NDUFS3:COX7A2L:NDUFA5 |
| MORF RAB1A | Neighborhood of RAB1A RAB1A, member RAS oncogene family in the MORF expression compendium | 191 | 3 | 0.000635 | 0.035952 | SDHC:COX8A:COX7A2L |
| MORF ANP32B | Neighborhood of ANP32B acidic (leucine-rich) nuclear phosphoprotein 32 family, member B in the MORF expression compendium | 198 | 3 | 0.000705 | 0.035952 | NDUFS3:COX7A2L:DDX39B |
| MORF SKP1A | Neighborhood of SKP1A S-phase kinase-associated protein 1A (p19A) in the MORF expression compendium | 203 | 3 | 0.000758 | 0.035952 | NDUFS3:COX7A2L:NDUFA5 |
| MORF RAC1 | Neighborhood of RAC1 ras-related C3 botulinum toxin substrate 1 (rho family, small GTP binding protein Rac1) in the MORF expression compendium | 211 | 3 | 0.000847 | 0.036182 | SDHC:COX8A:COX7A2L |
| MORF RAB11A | Neighborhood of RAB11A RAB11A, member RAS oncogene family in the MORF expression compendium | 60 | 2 | 0.001291 | 0.045261 | SDHC:COX7A2L |
| GNF2 RFC4 | Neighborhood of RFC4 replication factor C (activator 1) 4, 37kDa in the GNF2 expression compendium | 60 | 2 | 0.001291 | 0.045261 | CTPS1:PAICS |
| MORF SMC1L1 | Neighborhood of SMC1L1 NULL in the MORF expression compendium | 62 | 2 | 0.001378 | 0.045261 | NDUFS3:DDX39B |
| MORF DEK | Neighborhood of DEK DEK oncogene (DNA binding) in the MORF expression compendium | 264 | 3 | 0.001613 | 0.04919 | NDUFS3:COX7A2L:DDX39B |

Supplementary table 11: Cancer genes neighbourhood database (MsigDB c4). N indicates the total number of genes in the set while n shows the number of genes that SNPs mapped to the 37 genes identified. Gene sets were queried using FUMA GWAS. The adjusted P is an FDR-corrected P-value based on the number of gene sets examined.

| Gene Set | N | n | P-value | adj. P-value | Genes |
|---|---|---|---|---|---|
| OXIDATIVE PHOSPHORYLATION | 118 | 7 | 9.89E-12 | 1.84E-09 | SDHC:NDUFS3:COX8A:COX7A2L:NDUFA6:ATP6V1G2:NDUFA5 |
| ALZHEIMERS DISEASE | 159 | 7 | 8.16E-11 | 7.59E-09 | SDHC:NDUFS3:COX8A:COX7A2L:IL1B:NDUFA6:NDUFA5 |
| PARKINSONS DISEASE | 115 | 6 | 7.41E-10 | 4.60E-08 | SDHC:NDUFS3:COX8A:COX7A2L:NDUFA6:NDUFA5 |
| HUNTINGTONS DISEASE | 174 | 6 | 8.98E-09 | 4.18E-07 | SDHC:NDUFS3:COX8A:COX7A2L:NDUFA6:NDUFA5 |
| PURINE METABOLISM | 159 | 4 | 1.15E-05 | 0.000429 | PAPSS2:PFAS:PPAT:PAICS |
| TRYPTOPHAN METABOLISM | 40 | 2 | 0.000575 | 0.016061 | ALDH2:CYP1A2 |
| TYPE I DIABETES MELLITUS | 41 | 2 | 0.000604 | 0.016061 | INS:IL1B |
| CYTOKINE CYTOKINE RECEPTOR INTERACTION | 265 | 3 | 0.00163 | 0.03621 | IL1B:ACVR2A:KDR |
| LEISHMANIA INFECTION | 70 | 2 | 0.001752 | 0.03621 | IL1B:JAK2 |
| CARDIAC MUSCLE CONTRACTION | 74 | 2 | 0.001955 | 0.036368 | COX8A:COX7A2L |
| PROGESTERONE MEDIATED OOCYTE MATURATION | 85 | 2 | 0.002569 | 0.043437 | INS:CCNB1 |

Supplementary table 12: KEGG pathways. N indicates the total number of genes in the set while n shows the number of genes that SNPs mapped to the 37 genes identified. Gene sets were queried using FUMA GWAS. The adjusted P is an FDR-corrected P-value based on the number of gene sets examined.

| Gene Set | N | n | P-value | adj. P-value | Genes |
|---|---|---|---|---|---|
| RESPIRATORY ELECTRON TRANSPORT | 89 | 6 | 1.56E-10 | 2.33E-07 | SDHC:NDUFS3:COX8A:COX7A2L:NDUFA6:NDUFA5 |
| RESPIRATORY ELECTRON TRANSPORT ATP SYNTHESIS BY CHEMIOSMOTIC COUPLING AND HEAT PRODUCTION BY UNCOUPLING PROTEINS | 110 | 6 | 5.66E-10 | 4.24E-07 | SDHC:NDUFS3:COX8A:COX7A2L:NDUFA6:NDUFA5 |
| THE CITRIC ACID TCA CYCLE AND RESPIRATORY ELECTRON TRANSPORT | 161 | 6 | 5.64E-09 | 2.82E-06 | SDHC:NDUFS3:COX8A:COX7A2L:NDUFA6:NDUFA5 |
| PURINE RIBONUCLEOSIDE MONOPHOSPHATE BIOSYNTHESIS | 12 | 3 | 1.36E-07 | 5.10E-05 | PFAS:PPAT:PAICS |
| NUCLEOBASE BIOSYNTHESIS | 15 | 3 | 2.81E-07 | 8.42E-05 | PFAS:PPAT:PAICS |
| METABOLISM OF NUCLEOTIDES | 99 | 4 | 1.76E-06 | 0.000439 | CTPS1:PFAS:PPAT:PAICS |
| COMPLEX I BIOGENESIS | 49 | 3 | 1.11E-05 | 0.002386 | NDUFS3:NDUFA6:NDUFA5 |
| BIOLOGICAL OXIDATIONS | 221 | 4 | 4.19E-05 | 0.007855 | PAPSS2:ALDH2:CYP1A2:GSS |
| SIGNALING BY RECEPTOR TYROSINE KINASES | 468 | 5 | 5.24E-05 | 0.00872 | IGF2:INS:KDR:ATP6V1G2:JAK2 |
| PHASE II CONJUGATION OF COMPOUNDS | 109 | 3 | 0.000122 | 0.01836 | PAPSS2:CYP1A2:GSS |
| INSULIN RECEPTOR RECYCLING | 26 | 2 | 0.000242 | 0.032917 | INS:ATP6V1G2 |

Supplementary table 13: Reactome . N indicates the total number of genes in the set while n shows the number of genes that SNPs mapped to the 37 genes identified. Gene sets were queried using FUMA GWAS. The adjusted P is an FDR-corrected P-value based on the number of gene sets examined.

| Gene Set | N | n | P-value | adj. P-value | Genes |
|---|---|---|---|---|---|
| KEGG OXIDATIVE PHOSPHORYLATION | 118 | 7 | 9.89E-12 | 2.17E-08 | SDHC:NDUFS3:COX8A:COX7A2L:NDUFA6:ATP6V1G2:NDUFA5 |
| KEGG ALZHEIMERS DISEASE | 159 | 7 | 8.16E-11 | 8.97E-08 | SDHC:NDUFS3:COX8A:COX7A2L:IL1B:NDUFA6:NDUFA5 |
| REACTOME RESPIRATORY ELECTRON TRANSPORT | 89 | 6 | 1.56E-10 | 1.14E-07 | SDHC:NDUFS3:COX8A:COX7A2L:NDUFA6:NDUFA5 |
| REACTOME RESPIRATORY ELECTRON TRANSPORT ATP SYNTHESIS BY CHEMIOSMOTIC COUPLING AND HEAT PRODUCTION BY UNCOUPLING PROTEINS | 110 | 6 | 5.66E-10 | 3.11E-07 | SDHC:NDUFS3:COX8A:COX7A2L:NDUFA6:NDUFA5 |
| KEGG PARKINSONS DISEASE | 115 | 6 | 7.41E-10 | 3.26E-07 | SDHC:NDUFS3:COX8A:COX7A2L:NDUFA6:NDUFA5 |
| REACTOME THE CITRIC ACID TCA CYCLE AND RESPIRATORY ELECTRON TRANSPORT | 161 | 6 | 5.64E-09 | 2.07E-06 | SDHC:NDUFS3:COX8A:COX7A2L:NDUFA6:NDUFA5 |
| KEGG HUNTINGTONS DISEASE | 174 | 6 | 8.98E-09 | 2.82E-06 | SDHC:NDUFS3:COX8A:COX7A2L:NDUFA6:NDUFA5 |
| REACTOME PURINE RIBONUCLEOSIDE MONOPHOSPHATE BIOSYNTHESIS | 12 | 3 | 1.36E-07 | 3.74E-05 | PFAS:PPAT:PAICS |
| REACTOME NUCLEOBASE BIOSYNTHESIS | 15 | 3 | 2.81E-07 | 6.86E-05 | PFAS:PPAT:PAICS |
| REACTOME METABOLISM OF NUCLEOTIDES | 99 | 4 | 1.76E-06 | 0.000387 | CTPS1:PFAS:PPAT:PAICS |
| REACTOME COMPLEX I BIOGENESIS | 49 | 3 | 1.11E-05 | 0.002115 | NDUFS3:NDUFA6:NDUFA5 |
| KEGG PURINE METABOLISM | 159 | 4 | 1.15E-05 | 0.002115 | PAPSS2:PFAS:PPAT:PAICS |
| REACTOME BIOLOGICAL OXIDATIONS | 221 | 4 | 4.19E-05 | 0.007091 | PAPSS2:ALDH2:CYP1A2:GSS |
| REACTOME SIGNALING BY RECEPTOR TYROSINE KINASES | 468 | 5 | 5.24E-05 | 0.008224 | IGF2:INS:KDR:ATP6V1G2:JAK2 |
| REACTOME PHASE II CONJUGATION OF COMPOUNDS | 109 | 3 | 0.000122 | 0.017955 | PAPSS2:CYP1A2:GSS |
| NABA SECRETED FACTORS | 343 | 4 | 0.000228 | 0.02951 | IGF2:INS-IGF2:INS:IL1B |
| REACTOME INSULIN RECEPTOR RECYCLING | 26 | 2 | 0.000242 | 0.02951 | INS:ATP6V1G2 |
| PID IL27 PATHWAY | 26 | 2 | 0.000242 | 0.02951 | IL1B:JAK2 |

Supplementary table 14: All canonical pathways database (MsigDB c2). N indicates the total number of genes in the set while n shows the number of genes that SNPs mapped to the 37 genes identified. Gene sets were queried using FUMA GWAS. The adjusted P is an FDR-corrected P-value based on the number of gene sets examined.

| Gene Set | Brief Description | N | n | P-value | adj. P-value | Genes |
|---|---|---|---|---|---|---|
| chr4q12 | Ensemble 103 genes in cytogenetic band chr4q12 | 73 | 3 | 3.71E-05 | 0.011085 | KDR:PPAT:PAICS |

Supplementary table 15: All Positional Gene Set database (MsigDB c1). N indicates the total number of genes in the set while n shows the number of genes that SNPs mapped to the 37 genes identified. Gene sets were queried using FUMA GWAS. The adjusted P is an FDR-corrected P-value based on the number of gene sets examined.

| Gene Set | Brief Description | Publications | Species | N | n | P-value | adj. P-value | Genes |
|---|---|---|---|---|---|---|---|---|
| KEGG OXIDATIVE PHOSPHORYLATION | - | | | 118 | 7 | 9.89E-12 | 5.44E-08 | SDHC:NDUFS3:COX8A:COX7A2L:NDUFA6:ATP6V1G2:NDUFA5 |
| KEGG ALZHEIMERS DISEASE | - | | | 159 | 7 | 8.16E-11 | 2.14E-07 | SDHC:NDUFS3:COX8A:COX7A2L:IL1B:NDUFA6:NDUFA5 |
| MOOTHA VOXPHOS | Genes involved in oxidative phosphorylation; based on literature and sequence annotation resources and converted to Affymetrix HG-U133A probe sets. | Pubmed 12808457 Authors: Mootha VK,Lindgren CM,Eriksson KF,Subramanian A,Sihag S,Lehar J,Puigserver P,Carlsson E,Ridderstråle M,Laurila E,Houstis N,Daly MJ,Patterson N,Mesirov JP,Golub TR,Tamayo P,Spiegelman B,Lander ES,Hirschhorn JN,Altshuler D,Groop LC | Homo Sapiens | 87 | 6 | 1.35E-10 | 2.14E-07 | SDHC:NDUFS3:COX8A:COX7A2L:NDUFA6:NDUFA5 |
| REACTOME RESPIRATORY ELECTRON TRANSPORT | - | | | 89 | 6 | 1.56E-10 | 2.14E-07 | SDHC:NDUFS3:COX8A:COX7A2L:NDUFA6:NDUFA5 |
| REACTOME RESPIRATORY ELECTRON TRANSPORT ATP SYNTHESIS BY CHEMIOSMOTIC COUPLING AND HEAT PRODUCTION BY UNCOUPLING PROTEINS | - | | | 110 | 6 | 5.66E-10 | 6.23E-07 | SDHC:NDUFS3:COX8A:COX7A2L:NDUFA6:NDUFA5 |

| | | | | | | | | |
|---|---|---|---|---|---|---|---|---|
| KEGG PARKINSONS DISEASE | - | | | 115 | 6 | 7.41E-10 | 6.80E-07 | SDHC:NDUFS3:COX8A:COX7A2L:NDUFA6:NDUFA5 |
| MOOTHA HUMAN MITODB 6 2002 | Genes involved in oxidative phosphorylation; based on literature and sequence annotation resources and converted to Affymetrix HG-U133A probe sets. | Pubmed 12808457 Authors: Mootha VK,Lindgren CM,Eriksson KF,Subramanian A,Sihag S,Lehar J,Puigserver P,Carlsson E,Ridderstråle M,Laurila E,Houstis N,Daly MJ,Patterson N,Mesirov JP,Golub TR,Tamayo P,Spiegelman B,Lander ES,Hirschhorn JN,Altshuler D,Groop LC | Homo Sapiens | 430 | 8 | 2.90E-09 | 2.28E-06 | SDHC:NDUFS3:COX8A:ALDH2:POLG2:COX7A2L:NDUFA6:NDUFA5 |
| REACTOME THE CITRIC ACID TCA CYCLE AND RESPIRATORY ELECTRON TRANSPORT | - | | | 161 | 6 | 5.64E-09 | 3.88E-06 | SDHC:NDUFS3:COX8A:COX7A2L:NDUFA6:NDUFA5 |
| KEGG HUNTINGTONS DISEASE | - | | | 174 | 6 | 8.98E-09 | 5.49E-06 | SDHC:NDUFS3:COX8A:COX7A2L:NDUFA6:NDUFA5 |
| MOOTHA MITOCHONDRIA | Mitochondrial genes | Pubmed 12808457 Authors: Mootha VK,Lindgren CM,Eriksson KF,Subramanian A,Sihag S,Lehar J,Puigserver P,Carlsson E,Ridderstråle M,Laurila E,Houstis N,Daly MJ,Patterson N,Mesirov JP,Golub TR,Tamayo P,Spiegelman B,Lander | Homo Sapiens | 449 | 7 | 1.07E-07 | 5.89E-05 | SDHC:NDUFS3:COX8A:ALDH2:POLG2:NDUFA6:NDUFA5 |

| | | | | | | | | |
|---|---|---|---|---|---|---|---|---|
| | | ES,Hirschhorn JN,Altshuler D,Groop LC | | | | | | |
| REACTOME PURINE RIBONUCLEOSIDE MONOPHOSPHATE BIOSYNTHESIS | - | | | 12 | 3 | 1.36E-07 | 6.80E-05 | PFAS:PPAT:PAICS |
| REACTOME NUCLEOBASE BIOSYNTHESIS | - | | | 15 | 3 | 2.81E-07 | 0.000129 | PFAS:PPAT:PAICS |
| WONG MITOCHONDRIA GENE MODULE | Genes that comprise the mitochondria gene module | Pubmed 18199530 Authors: Wong DJ,Nuyten DS,Regev A,Lin M,Adler AS,Segal E,van de Vijver MJ,Chang HY | Homo Sapiens | 218 | 5 | 1.31E-06 | 0.000553 | NDUFS3:COX8A:COX7A2L:NDUFA6:NDUFA5 |
| REACTOME METABOLISM OF NUCLEOTIDES | - | | | 99 | 4 | 1.76E-06 | 0.000691 | CTPS1:PFAS:PPAT:PAICS |
| DANG BOUND BY MYC | Genes whose promoters are bound by MYC | Pubmed 14519204 Authors: Zeller KI,Jegga AG,Aronow BJ,O'Donnell KA,Dang CV | Homo Sapiens | 1059 | 8 | 2.82E-06 | 0.001034 | ALDH2:COX7A2L:NDUFA6:PPAT:PAICS:CCNB1:ATP6V1G2:NFKBIL1 |
| YOSHIMURA MAPK8 TARGETS UP | Genes up-regulated in vascular smooth muscle cells (VSMC) by MAPK8 (JNK1) | Pubmed 16311603 Authors: Yoshimura K,Aoki H,Ikeda Y,Fujii K,Akiyama N,Furutani A,Hoshii Y,Tanaka N,Ricci R,Ishihara T,Esato K,Hamano K,Matsuzaki M | Rattus norvegicus | 1171 | 8 | 5.91E-06 | 0.002033 | IGF2:INS:COX8A:POLG2:ACVR2A:GSS:NDUFA5:JAK2 |
| REACTOME COMPLEX I BIOGENESIS | - | | | 49 | 3 | 1.11E-05 | 0.003528 | NDUFS3:NDUFA6:NDUFA5 |
| KEGG PURINE METABOLISM | - | | | 159 | 4 | 1.15E-05 | 0.003528 | PAPSS2:PFAS:PPAT:PAICS |

| | | | | | | | | |
|---|---|---|---|---|---|---|---|---|
| TENEDINI MEGAKARYOCYTE MARKERS | Genes essential to the development of megakaryocytes, as expressed in normal cells and essential thrombocythemic cells (ET). | Pubmed 15271793 Authors: Tenedini E,Fagioli ME,Vianelli N,Tazzari PL,Ricci F,Tagliafico E,Ricci P,Gugliotta L,Martinelli G,Tura S,Baccarani M,Ferrari S,Catani L | Homo Sapiens | 61 | 3 | 2.16E-05 | 0.006256 | IL1B:CCNB1:JAK2 |
| KIM BIPOLAR DISORDER OLIGODENDROCYTE DENSITY CORR UP | Genes whose expression significantly and positively correlated with oligodendrocyte density in layer VI of BA9 brain region in patients with bipolar disorder. | Pubmed 18762803 Authors: Kim S,Webster MJ | Homo Sapiens | 681 | 6 | 2.52E-05 | 0.006942 | NDUFS3:ALDH2:COX7A2L:NDUFA6:NDUFA5:GLDC |
| MOOTHA PGC | Genes up-regulated in differentiating C2C12 cells (myoblasts) upon expression of PPARGC1A off an adenoviral vector. | Pubmed 12808457 Authors: Mootha VK,Lindgren CM,Eriksson KF,Subramanian A,Sihag S,Lehar J,Puigserver P,Carlsson E,Ridderstråle M,Laurila E,Houstis N,Daly MJ,Patterson N,Mesirov JP,Golub TR,Tamayo P,Spiegelman B,Lander ES,Hirschhorn JN,Altshuler D,Groop LC | Homo Sapiens | 421 | 5 | 3.17E-05 | 0.008298 | NDUFS3:COX8A:GSS:NDUFA6:NDUFA5 |
| REACTOME BIOLOGICAL OXIDATIONS | - | | | 221 | 4 | 4.19E-05 | 0.010482 | PAPSS2:ALDH2:CYP1A2:GSS |
| DUTERTRE ESTRADIOL RESPONSE 6HR UP | Genes up-regulated in MCF7 cells (breast cancer) at 6 h of estradiol treatment. | Pubmed 20406972 Authors: Dutertre M,Gratadou L,Dardenne E,Germann S,Samaan S,Lidereau R,Driouch K,de la Grange P,Auboeuf D | Homo Sapiens | 228 | 4 | 4.73E-05 | 0.011179 | CTPS1:MYBBP1A:PAICS:JAK2 |

| | | | | | | | | |
|---|---|---|---|---|---|---|---|---|
| SCHUHMACHER MYC TARGETS UP | Genes up-regulated in P493-6 cells (Burkitt's lymphoma) induced to express MYC. | Pubmed 11139609 Authors: Schuhmacher M,Kohlhuber F,Hölzel M,Kaiser C,Burtscher H,Jarsch M,Bornkamm GW,Laux G,Polack A,Weidle UH,Eick D | Homo Sapiens | 80 | 3 | 4.88E-05 | 0.011179 | CTPS1:PPAT:PAICS |
| REACTOME SIGNALING BY RECEPTOR TYROSINE KINASES | - | | | 468 | 5 | 5.24E-05 | 0.011521 | IGF2:INS:KDR:ATP6V1G2:JAK2 |
| WINTER HYPOXIA METAGENE | Genes regulated by hypoxia, based on literature searches. | Pubmed 17409455 Authors: Winter SC,Buffa FM,Silva P,Miller C,Valentine HR,Turley H,Shah KA,Cox GJ,Corbridge RJ,Homer JJ,Musgrove B,Slevin N,Sloan P,Price P,West CM,Harris AL | Homo Sapiens | 240 | 4 | 5.78E-05 | 0.012219 | IGF2:KDR:PPAT:PAICS |
| BLALOCK ALZHEIMERS DISEASE DN | Genes down-regulated in brain from patients with Alzheimer's disease. | Pubmed 14769913 Authors: Blalock EM,Geddes JW,Chen KC,Porter NM,Markesbery WR,Landfield PW | Homo Sapiens | 1244 | 7 | 8.54E-05 | 0.017405 | NDUFS3:COX8A:ALDH2:COX7A2L:ATP6V1G2:NDUFA5:JAK2 |
| FLECHNER BIOPSY KIDNEY TRANSPLANT REJECTED VS OK DN | Genes down-regulated in kidney biopsies from patients with acute transplant rejection compared to the biopsies from patients with well functioning kidneys more than 1-year post transplant. | Pubmed 15307835 Authors: Flechner SM,Kurian SM,Head SR,Sharp SM,Whisenant TC,Zhang J,Chismar JD,Horvath S,Mondala T,Gilmartin T,Cook DJ,Kay SA,Walker JR,Salomon DR | Homo Sapiens | 552 | 5 | 0.000114 | 0.022366 | SDHC:ALDH2:KDR:NDUFA5:GLDC |
| TIEN INTESTINE PROBIOTICS 24HR UP | Genes up-regulated in Caco-2 cells (intestinal epithelium) after | Pubmed 16394013 Authors: Tien MT,Girardin SE,Regnault B,Bourhis Le | Homo Sapiens | 560 | 5 | 0.000122 | 0.022459 | SDHC:NDUFA6:CCNB1:NDUFA5:GLDC |

| | | | | | | | | |
|---|---|---|---|---|---|---|---|---|
| | coculture with the probiotic bacteria L. casei for 24h. | L,Dillies MA,Coppée JY,Bourdet-Sicard R,Sansonetti PJ,Pédron T | | | | | | |
| REACTOME PHASE II CONJUGATION OF COMPOUNDS | - | | | 109 | 3 | 0.000122 | 0.022459 | PAPSS2:CYP1A2:GSS |
| TARTE PLASMA CELL VS PLASMABLAST DN | - | Pubmed 12663452 Authors: Tarte K,Zhan F,De Vos J,Klein B,Shaughnessy J Jr | Homo Sapiens | 307 | 4 | 0.000149 | 0.026477 | CTPS1:PAICS:CCNB1:GLDC |
| DAIRKEE CANCER PRONE RESPONSE BPA E2 | 'Cancer prone response profile' (CPRP): genes common to estradiol and bisphenol A [PubChem=5757;6623] response of epithelial cell cultures from patients at high risk of breast cancer. | Pubmed 18381411 Authors: Dairkee SH,Seok J,Champion S,Sayeed A,Mindrinos M,Xiao W,Davis RW,Goodson WH | Homo Sapiens | 118 | 3 | 0.000155 | 0.026626 | CTPS1:NDUFS3:COX8A |
| CAIRO HEPATOBLASTOMA CLASSES UP | Genes up-regulated in robust Cluster 2 (rC2) of hepatoblastoma samples compared to those in the robust Cluster 1 (rC1). | Pubmed 19061838 Authors: Cairo S,Armengol C,Reyniès De A,Wei Y,Thomas E,Renard CA,Goga A,Balakrishnan A,Semeraro M,Gresh L,Pontoglio M,Strick-Marchand H,Levillayer F,Nouet Y,Rickman D,Gauthier F,Branchereau S,Brugières L,Laithier V,Bouvier R,Boman F,Basso G,Michiels JF,Hofman P,Arbez-Gindre F,Jouan H,Rousselet-Chapeau MC,Berrebi D,Marcellin L,Plenat F,Zachar D,Joubert M,Selves J,Pasquier D,Bioulac-Sage P,Grotzer | Homo Sapiens | 608 | 5 | 0.000179 | 0.029766 | MYBBP1A:PFAS:PPAT:CCNB1:GLDC |

| | | | | | | | | |
|---|---|---|---|---|---|---|---|---|
| | | M,Childs M,Fabre M,Buendia MA | | | | | | |
| NABA SECRETED FACTORS | Genes encoding secreted soluble factors | Pubmed 22159717 Authors: Naba A,Clauser KR,Hoersch S,Liu H,Carr SA,Hynes RO | Homo Sapiens | 343 | 4 | 0.000228 | 0.03687 | IGF2:INS-IGF2:INS:IL1B |
| REACTOME INSULIN RECEPTOR RECYCLING | - | | | 26 | 2 | 0.000242 | 0.036911 | INS:ATP6V1G2 |
| PID IL27 PATHWAY | - | Pubmed 18832364 Authors: Schaefer CF,Anthony K,Krupa S,Buchoff J,Day M,Hannay T,Buetow KH | Homo Sapiens | 26 | 2 | 0.000242 | 0.036911 | IL1B:JAK2 |
| BASSO B LYMPHOCYTE NETWORK | Genes which comprise the top 1% of highly interconnected genes (major hubs) that account for most of gene interactions in the reconstructed regulatory networks from expression profiles in B lymphocytes. | Pubmed 15778709 Authors: Basso K,Margolin AA,Stolovitzky G,Klein U,Dalla-Favera R,Califano A | Homo Sapiens | 145 | 3 | 0.000284 | 0.042248 | CTPS1:PAICS:CCNB1 |
| BENPORATH ES 1 | Set 'ES exp1': genes overexpressed in human embryonic stem cells according to 5 or more out of 20 profiling studies. | Pubmed 18443585 Authors: Ben-Porath I,Thomson MW,Carey VJ,Ge R,Bell GW,Regev A,Weinberg RA | Homo Sapiens | 378 | 4 | 0.00033 | 0.047701 | PFAS:PAICS:CCNB1:GLDC |

Supplementary table 16: Curated gene sets. N indicates the total number of genes in the set while n shows the number of genes that SNPs mapped to the 37 genes identified. Gene sets were queried using FUMA GWAS. The adjusted P is an FDR-corrected P-value based on the number of gene sets examined.
